# Molecular and genetic mechanisms conferring dissolution of dioecy in *Diospyros oleifera* Cheng

**DOI:** 10.1101/2022.10.08.511238

**Authors:** Peng Sun, Soichiro Nishiyama, Huawei Li, Yini Mai, Weijuan Han, Yujing Suo, Chengzhi Liang, Huilong Du, Songfeng Diao, Yiru Wang, Jiaying Yuan, Yue Zhang, Ryutaro Tao, Fangdong Li, Jianmin Fu

## Abstract

Dioecy, a sexual system of single-sex (gynoecious/androecious) individuals, is rare in flowering plants. This rarity may be a result of the frequent transition from dioecy into systems with co-sex individuals. Here, we report potential molecular and genetic mechanisms that underlie the dissolution of dioecy to monoecy and andro(gyno)monoecy, based on multiscale genome-wide investigations of 150 accessions of *Diospyros oleifera*. All co-sex *D. oleifera* plants, including monoecious and andro(gyno)monoecious individuals, possessed the male determinant gene *OGI*, implying that genetic factors control gynoecia development in genetically male *D. oleifera*. In both single- and co-sex plants, female function was expressed in the presence of a genome-wide decrease in methylation levels, along with sexually distinct regulatory networks of smRNAs and their targets. Furthermore, a genome-wide association study (GWAS) identified a genomic region and a *DUF247* gene cluster strongly associated with the monoecious phenotype, as well as several regions that may contribute to andromonoecy. Collectively, our findings imply stable breakdown of the dioecious system in *D. oleifera*, presumably a result of the genomic features of the sex-linked region.

## Introduction

Dioecy—male and female reproductive organs in separate individuals—is found in only approximately 6% of angiosperm species, but in diverse plant lineages (Renner and Ricklefs, 1995; Weiblen *et al*., 2000; Renner, 2014). This low rate of dioecy may indicate an evolutionary dead-end scenario (Heilbuth, 2000); however, it has been proposed that the dissolution of dioecy, which leads to re-evolution of the co-sex systems, occurs during sexual-system evolution (Kafer *et al*., 2017). Multiple empirical studies have yielded results consistent with this hypothesis (Badouin *et al*., 2020; Masuda *et al*., 2022).

Sex expression in the genus *Diospyros* is diverse, and several key genetic components controlling sexual expression have been identified (Akagi *et al*., 2014). Diploid *D. lotus* is dioecious, exhibiting separate male and female plants; polyploid *D. kaki* encompasses gynoecious (female flower only), androecious (male flower only), monoecious (male and female flowers), androgynomonoecious (male, female, and hermaphroditic flowers), and andromonoecious (male and hermaphroditic flowers) individuals (Li *et al*., 2016; Akagi *et al*., 2016a; Wang *et al*., 2021; Li *et al*., 2021). Similar sexual diversity was observed in diploid *D. oleifera* (Fig. 1; Figs. S1-3). In *Diospyros*, male flowers are typically observed in the form of a three-flower cyme, but female flowers are always solitary (Fig. 1). Dioecy is considered an ancestral form in *Diospyros*; therefore, it is reasonable to assume that the polygamous system in *D. kaki* and *D. oleifera* evolved from the dioecious system (Henry *et al*., 2018). In diploid dioecious *D. lotus*, the small RNA (smRNA)-coding gene *OGI* determines the formation of male trees via repression of the feminising gene *MeGI* (Akagi *et al*., 2014). In contrast, an additional layer of regulation in the form of DNA methylation of the *MeGI* promoter may contribute to monoecy in hexaploid *D. kaki* (Akagi *et al*., 2016a).

**Fig. 1.**
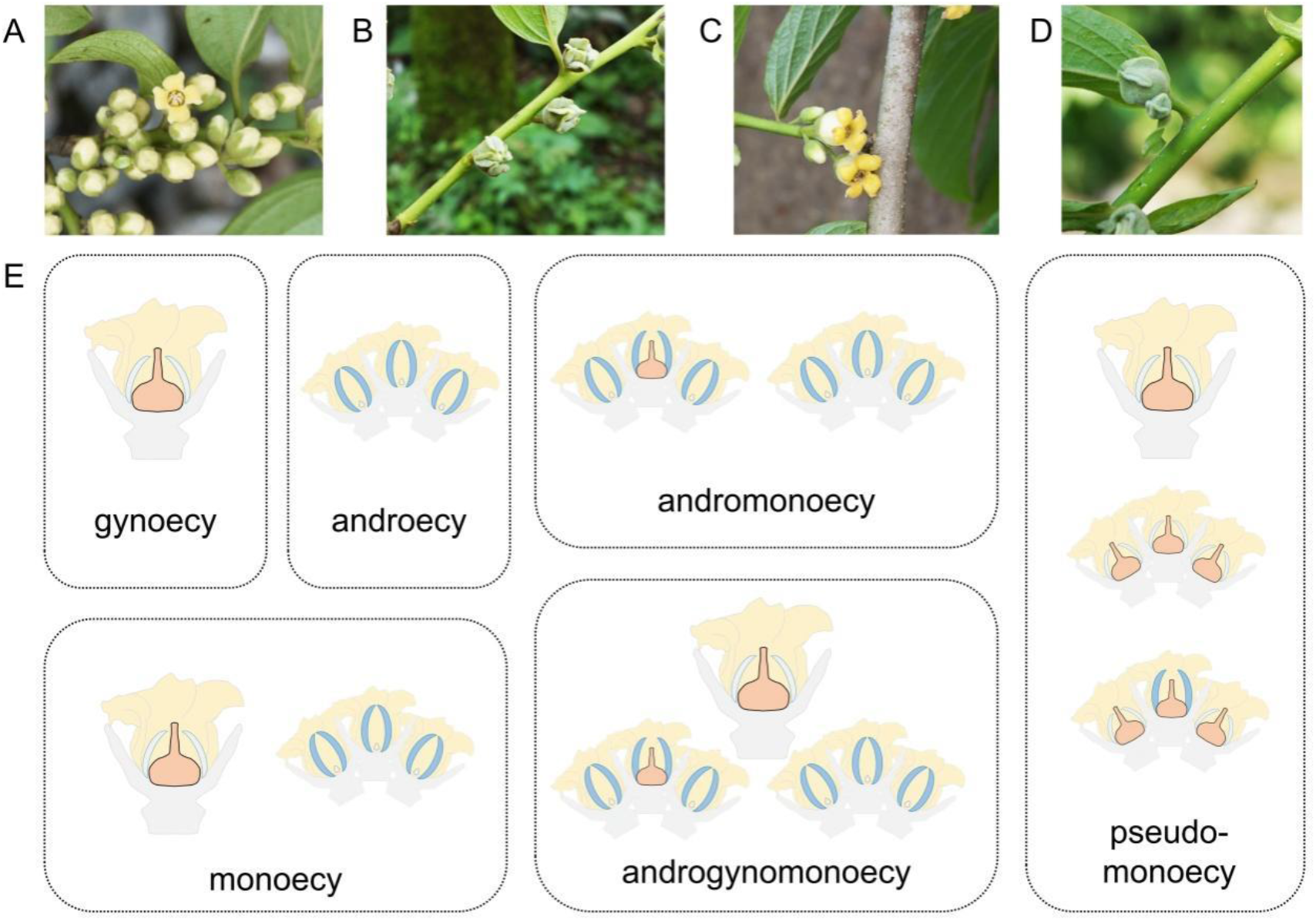
Diverse sex expression in *D. oleifera*. **(A)** male floral buds, **(B)** female floral buds, **(C)** hermaphroditic floral buds, and **(D)** female flower cyme in pseudo-monoecy. **(E)** Schematic representation of individual sexual types. Details of the sexual types are shown in Figs. S1-3.

Although several important molecular mechanisms underlying the sexuality determination of *D*. spp. have been identified, the molecular mechanisms that underlie the sexual diversity of some *D*. spp. (*e*.*g*., *D. kaki* and *D. oleifera*) are unclear. In this study, we used a genetic approach to characterise a *D. oleifera* population. The diploidy of *D. oleifera* with all types of sex expression was confirmed (Text S1; Fig. S4). The genome of diploid *D. oleifera* is simpler than the genome of cultivated polyploid *D. kaki*, and a new *D. oleifera* genome was assembled at the chromosome scale (Text S2; Figs. S5-S17; Tables S1-S12; Sun and Fu, 2022a). The development of *D. oleifera* floral buds is synchronous to the development of *D. kaki*. Therefore, *D. oleifera* is an excellent genetic model for the study of sex diversity in *Diospyros*.

In mid-April 2019 and 2021, we surveyed the sex phenotype of *D. oleifera* obtained from a natural population in Guangxi Zhuang Autonomous Region, China (Fig. S18). Two hundred eight plants with various types of sex expression were sampled in total (Table 1; Table S13). The genetic components of the *OGI*/*MeGI* system were examined in the *D. oleifera* population. Subsequently, comparative-methylome and whole-transcriptome analyses were performed to identify the molecular mechanism responsible for sexual diversity. Moreover, a genome-wide association study (GWAS) was conducted to determine candidate genomic regions and genes that contribute to the sexual diversity.

**Table 1.** Sex expression in a *D. oleifera* natural population.

| Individual sexual types | Number | DNA sampled | *OGI* genotype |  | Male expression |
| --- | --- | --- | --- | --- | --- |
|  |  |  | *OGI*<sup>+</sup> | *OGI*<sup>-</sup> |  |
| Gynoecy | 116 | 58 | 4 | 54 | - |
| Pseudo-monoecy | 2 | 2 | 0 | 2 | - |
| Androecy | 45 | 45 | 45 | 0 | + |
| Monoecy | 28 | 28 | 27 | 1 | + |
| Androgynomonoecy | 10 | 10 | 10 | 0 | + |
| Andromonoecy | 7 | 7 | 7 | 0 | + |
| Total | 208 | 150 | 93 | 57 |  |

## Methods

### Plant material and phenotyping

A *D. oleifera* collection from Guilin, Guangxi Zhuang Autonomous Region, China, was evaluated for sexual expression in the 2019 and 2021 seasons. Two hundred eight *D. oleifera* trees were found in the natural population. In *D. oleifera*, the flower sex on a single shoot is uniform. Thus, for simplicity and accuracy, large flowering mother branches (approximately 1.5 m in height × 1.5 m in width) containing ≥ 20 flowering shoots (each uniformly bearing female, male, or hermaphroditic floral buds) were used to calculate the proportions of female, male, and hermaphroditic shoots in monoecious, androgynomonoecious, and andromonoecious trees. At least five large flowering mother branches of each tree were selected; the highest female and hermaphroditic shoot proportions were measured and used for GWAS as an indication of feminising ability on the co-sex tree.

### Reference sequence construction

The female plant ‘*D. oleifera* 1’ was used to optimise the published version of the *D. oleifera* genome (Suo *et al*., 2020) using a BioNano optical mapping-assisted assembly (Method S2). The male-specific region (MSR) was absent in this genome; thus, the unmapped resequencing reads of 14 androecious, 4 andromonoecious, 15 monoecious, 7 androgynomonoecious, and 2 pseudo-monoecious *D. oleifera* individuals in the mapping with the Burrows-Wheeler Aligner mem option and the paired-end model (Li and Durbin, 2010) were extracted using SAMtools (Li *et al*., 2009). The reads were assembled using SoapDenovo (Li *et al*., 2010) to construct the male-unmapped sequences (Method S3). The *D. oleifera* main genome (Sun and Fu, 2022a), male-unmapped sequences (Sun and Fu, 2022b), and the *D. oleifera* chloroplast genome (Fu *et al*., 2017) were combined as a reference genome for methylome, whole-transcriptome, and resequencing analyses.

### Methylome and whole-transcriptome analyses

Floral bud and immature stem tissues of flowering shoots were sampled in mid-April (April 15–17th), 2019, which is a key period for the differentiation of flower sex types (Methods S5). Those samples (Tables S14 and S15) were used for the whole-genome bisulphite sequencing, whole-transcriptome sequencing, and small RNA (smRNA) sequencing. All reads obtained were mapped to the reference *D. oleifera* genome. Details of library construction, sequencing, and analysis are provided in Methods S6-S9.

The smRNAs-Seq reads were also mapped to the *OGI* sequence from the *D. lotus* genome (Akagi *et al*., 2020), *MeGI* sequence from the *D. oleifera* reference genome, and the ‘*Kali*’ sequence (Akagi *et al*., 2016a), using the method established by Akagi *et al*. (2016a). Here, the smRNAs-Seq reads mapped onto the *MeGI* gene body were referred to as *smMeGI*. The accumulation levels of *smMeGI* and each fragment were recorded as reads per million reads.

### Resequencing analysis

A set of 150 *D. oleifera* (Table 1; Table S13) was selected from the collection and used for resequencing analysis. Short read resequencing (PE150) by the Illumina NovaSeq 6000 platform yielded 3.03 Tb of raw data with ∼20-fold genomic coverage for each sample. All reads were mapped to the *D. oleifera* reference genome, the *D. lotus OGI* genomic sequence, and the *D. lotus* MSR sequence (Akagi *et al*., 2020) using the Bwa mem option and the paired-end model. SAMtools and the Genome Analysis Toolkit (version 2.4-7-g5e89f01) were used to label SNPs and insertion-deletions (indels). Polymorphisms that matched the following four criteria were filtered out: > 2 alleles, variants beyond the read depth between half and twice the genome-wide average, missing rates ≥ 0.25, and minor allele frequency < 0.05. The linkage disequilibrium (LD) was evaluated using the pairwise squared Pearson’s correlation coefficient (r^2^) calculated by PLINK version 1.9 (Slifer, 2018). LD pruning was conducted by a standard method (PLINK –indep 50 5 2).

Using the filtered variant sets, the familial relationships and sample uniqueness were evaluated based on the PI_HAT value computed by PLINK. Furthermore, the population structure of the 150 trees was estimated using principal component analysis (PCA) in EIGENSTRAT software (Price *et al*., 2006). The maximum-likelihood phylogenetic tree was constructed using MEGA-X (Kumar *et al*., 2016) with 1000 replicates using the following parameters: gaps/missing data, partial deletion; site coverage cut-off, 90%; general time reversible model; and rates among sites, uniform.

### GWAS

GWAS was performed using the linear mixed model in the R package rrBLUP (Endelman, 2011). A kinship (K) matrix (generated with the A.mat function of rrBLUP) was included in the linear mixed model, along with 6 principal components (PCs) for 90 individuals with male production (Fig. S55B) and 4 PCs for all 150 individuals (Fig. S57). Bonferroni correction (corrected *P* < 0.05) was used to determine the genome-wide significance thresholds. The LD patterns surrounding GWAS peaks were visualised using the R package LDheatmap (Shin *et al*., 2006) for chromosome 7, and using Haploview (http://www.broadinstitute.org/haploview) for chromosomes 4, 2, 11, and 14. The regions with pairwise r^2^ > 0.5 were regarded as candidate LD blocks.

## Results

### Characterisation of the male-specific region (MSR) in *D. oleifera*

We analysed the presence of MSR (Akagi *et al*., 2020) and the *OGI* gene in 150 *D. oleifera* individuals (Table 1). We found that 54 of 58 gynoecious individuals did not contain MSR and were *OGI*-negative (*OGI* ^-^). In contrast, all 45 androecious individuals contained MSR and were *OGI*-positive (*OGI*^+^). All co-sex plants (monoecious, androgynomonoecious, and andromonoecious individuals), except for one monoecious individual, contained MSR and were *OGI*^+^; these findings strongly suggested that *OGI* is required for male tissue production in general sex determination of *D. oleifera*, as in monoecious *D. kaki* (Akagi *et al*., 2016b). Among the co-sex plants, the ability to produce male flowers was variable (Table S13), and thus the four exceptional gynoecious plants possessing *OGI* were considered female-biased monoecious plants. We did not include plants with inconsistent phenotypes (four gynoecious with *OGI*) in subsequent analyses.

Several individuals showed distinct flower sexes and morphologies, which comprised pseudo-monoecy. The pseudo-monoecious trees, lacking MSR and *OGI*, form two- or three-flower cymes with a female (or occasionally hermaphroditic) flower in the middle and one or two abnormal small female flowers at the sides (Fig. 1D; Fig. S3).

None of the *OGI*^+^ individuals harboured *Kali* (a short interspersed nuclear element [SINE]-like insertion) on the promoter of *OGI* (Methods S4; Fig. S19), which is presumed to silence *OGI* in the hexaploid monoecious *D. kaki* (Akagi *et al*., 2016a). Therefore, previous molecular genetic knowledges for monoecious production in *D. kaki* might be not directly applicable to *D. oleifera*.

### Inconsistency between *smMeGI* abundance and *MeGI* transcripts in monoecious *D. oleifera*

We investigated the accumulation patterns of smRNAs on *OGI* and *MeGI* regions (*smMeGI*) in *D. oleifera* (Fig. S20); such patterns coincide with flower sex in *D. lotus* and *D. kaki* (Akagi *et al*., 2014; 2016a). In the single-sex (gynoecious and androecious plants) *D. oleifera*, greater accumulation of *smMeGI* was detected in floral buds from androecious plants (androecious male; A_M) than in floral buds from gynoecious plants (gynoecious female; G_F) (Fig. 2A), similar to findings in *D. lotus* and *D. kaki* (Akagi *et al*., 2016a). A similar pattern was observed in the immature stem tissues of flowering shoots (Fig. 2B). Consistent with the level of *smMeGI*, the level of *MeGI* transcripts was significantly higher in G_F than in A_M in mid-April (Fig. 2F), similar to findings in *D. lotus* and *D. kaki* (Akagi *et al*., 2014; 2016a).

**Fig. 2.**
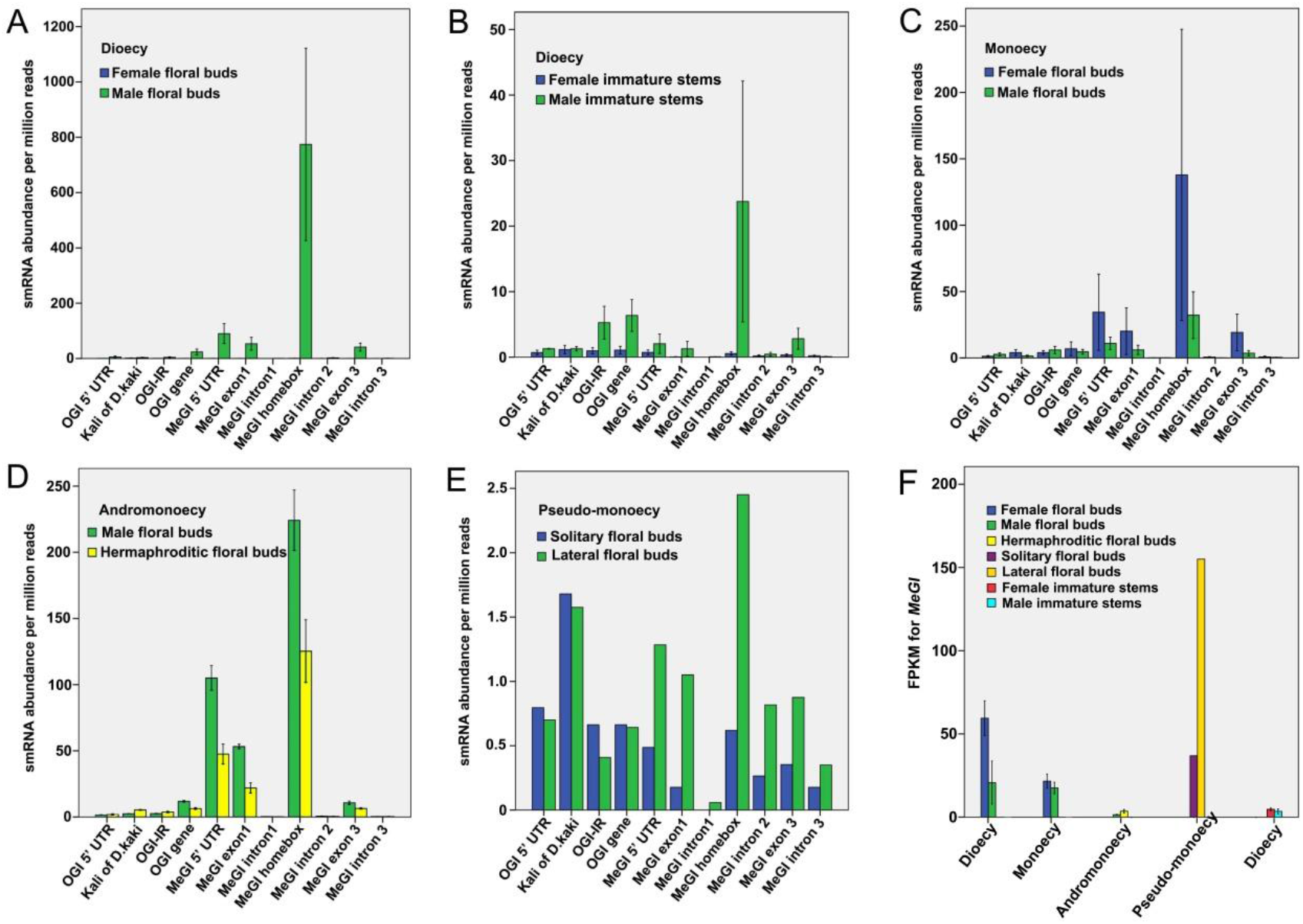
*MeGI* expression and smRNA accumulation in *D. oleifera*. **(A-E)** smRNA accumulation on *OGI*/*MeGI* genomic sequences in **(A)** female and male floral buds in dioecy; **(B)** stems of female and male shoots in dioecy; **(C)** female and male floral buds in monoecy; **(D)** male and hermaphroditic floral buds in andromonoecy; and **(E)** solitary floral buds and lateral floral buds of flower cymes in pseudo-monoecy. **(F)** Fragments per kilobase of transcript per million mapped reads (FPKM) values of *MeGI* in floral buds and stems. Data are means ± standard errors (three biological replicates) except for floral buds in pseudo-monoecious trees, for which no biological replicates were available.

In contrast, the *smMeGI* patterns in the co-sex types were distinct from the findings in previous reports. In the monoecious *D. oleifera*, where individuals exhibited separate male and female flowers, *smMeGI* accumulation was not significantly different between the female (monoecious female; M_F) and male floral buds (monoecious male; M_M) (Fig. 2C). Similarly, the level of *MeGI* expression was not significantly different between M_F and M_M; it was comparable with the level in male flower buds of androecious plants (A_M) (Fig. 2F).

In the hermaphroditic flowers, we observed an intermediate level of *smMeGI*. Among andromonoecious plants, the level of *smMeGI* accumulation was significantly lower in hermaphroditic floral buds (andromonoecious hermaphroditic; AM_H) than in male floral buds (andromonoecious male; AM_M) (Fig. 2D). In contrast to female flowers (G_F), where *smMeGI* was nearly absent, a substantial amount of *smMeGI* accumulation was detected in hermaphroditic flowers (AM_H) (Fig. 2A and D). The levels of *MeGI* expression were not significantly different between AM_M and AM_H (Fig. 2F).

In the *OGI*^―^ pseudo-monoecious plants, low *smMeGI* accumulation was observed in both solitary female floral buds (pseudo-monoecious solitary female; PM_SF) and lateral floral buds of the three flower cymes (pseudo-monoecious lateral female; PM_LF) (Fig. 2E). *MeGI* expression was higher in PM_SF and PM_LF than in male floral buds from androecious, monoecious, and andromonoecious plants, consistent with the level of gynoecia development (Fig. 2F).

### DNA methylation in *D. oleifera* floral buds and stems

In monoecious *D. kaki*, a lower methylation level of the *MeGI* promoter is associated with female flower formation in genetically male plants (Akagi *et al*., 2016a). We evaluated whether this mechanism is active in the co-sex *D. oleifera*, where sex expression is not controlled by the *OGI*/*MeGI* system. The DNA methylation levels of floral buds and stems were analysed by whole-genome bisulphite sequencing (Text S3; Tables S16-S21; Figs. S21-S23).

Unexpectedly, the DNA methylation patterns of the *MeGI* region in diverse *D. oleifera* matched the DNA methylation patterns of *D. kaki* and *D. lotus*, even in the co-sex types. The methylation levels of the 5’-untranslated region and exon of *MeGI* were significantly lower in female floral buds than in male floral buds obtained from monoecious plants (Fig. 3A). This pattern was also observed in andromonoecious and single-sex (gynoecious and androecious individuals) plants, which showed lower methylation levels in female tissues (AM_H, G_F, and G_S) than in male tissues (AM_M, A_M, and A_S) (Fig. 3A; Figs. S24 and S25).

**Fig. 3.**
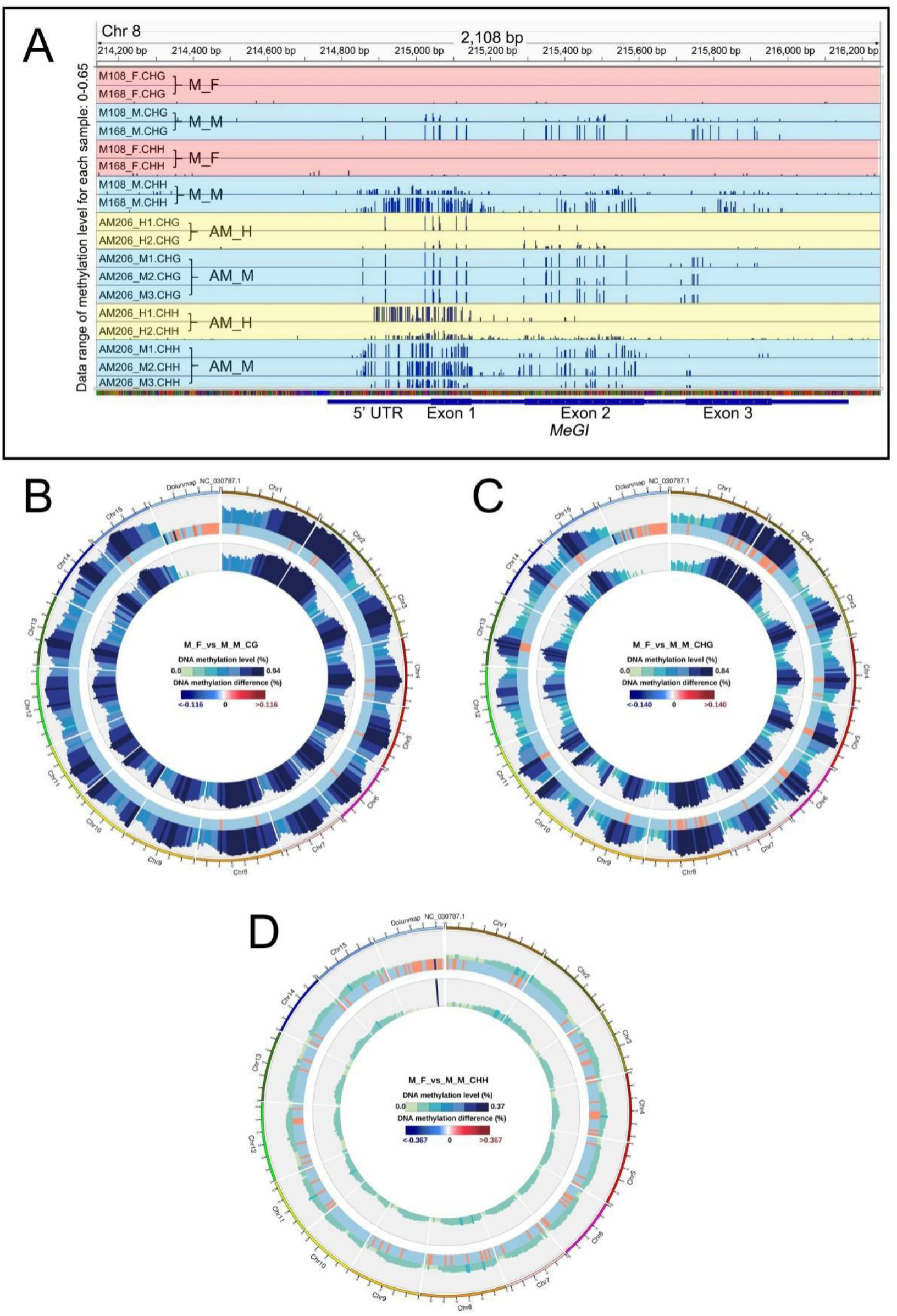
DNA methylation in flower buds from co-sex *D. oleifera*. **(A)** Methylation levels of the *MeGI* genomic region in floral buds from monoecious and andromonoecious *D. oleifera*. **(B-D)** Whole-genome comparison of methylation levels in the **(B)** CG, **(C)** CHG, and **(D)** CHH subcontexts between M_F and M_M. Tracks from outside to inside: methylation level of M_F; different methylation levels between M_F and M_M, where red and blue represent higher and lower methylation levels in M_F than in M_M, respectively; methylation level of M_M.

We next speculated that the discrepancy between the *smMeGI* level and DNA methylation in the co-sex plants is related to changes in the global methylation pattern. Notably, the DNA methylation levels of floral buds in the CG, CHG, and CHH subcontexts were lower in female tissues (M_F) than in male tissues (M_M) in almost all genomic regions (Fig. 3B, C, and D). The DNA methylation levels in the CG and CHG subcontexts in every part of gene body, as well as the surrounding regions, were generally lower in M_F than in M_M (Fig. S26A and B). Most of the differentially methylated regions were distant from genes, although there were numerous differentially methylated regions in the promoter, exon, and intronic regions (Fig. S26C, D, and E). A similar pattern was observed in the single-sex plants; the genome-wide DNA methylation levels in the CG, CHG, and CHH subcontexts were generally lower in females (G_F) than in males (A_M) (Fig. S27). The same trend was observed in stems of flowering shoots (Fig. S28). This finding was also observed for hermaphrodite flower formation. In andromonoecious plants, the DNA methylation levels of floral buds in all three subcontexts were lower in hermaphrodites (AM_H) than in males (AM_M) (Fig. S29). Therefore, female tissues showed lower global DNA methylation levels than male tissues in all sexual types, implying that a genome-wide decrease in DNA methylation promotes the development of gynoecia in both co- and single-sex systems.

As a potential regulator of DNA methylation, we investigated the expression levels of DNA methyltransferase/demethylase genes by RNA-Seq. A demethylase gene (evm.model.Chr7.1501), homologous to REPRESSOR OF SILENCING1 (*ROS1*) (Gong *et al*., 2002), was downregulated in AM_M compared with AM_H (Table S22), possibly explaining the dynamic modulation of genome-wide methylation levels in andromonoecious *D. oleifera*.

Many GO terms enriched in genes exhibited overlap with differentially methylated regions; however, most GO terms were not shared by the single- and co-sex systems (Table S23). This finding is consistent with the notion that mechanisms other than the active mechanisms in single-sex systems regulate sex differentiation in co-sex systems.

### Overlap of mRNA-miRNA functional modules between co- and single-sex systems

To characterise the functional overlap of gynoecia/androecia development in different sexual expression systems, whole-transcriptome analyses were conducted using 26 samples of floral buds and stems of flowering shoots (Text S4; Figs. S30-S54; Tables S24-S30). We found common sexually distinct regulatory networks of microRNAs (miRNAs) and their targets, as well as common functional enrichments for gynoecia development in single- and co-sex systems.

Based on the interaction network analysis and functional annotation, highly expressed miRNAs in female (or hermaphroditic) and their mRNA targets downregulated in female (or hermaphroditic) floral buds, and *vice versa* (*i*.*e*., low-level miRNA and high-level mRNA targets in females), were identified in single- and co-sex systems (Fig. 4; Tables S28 and S29). At least two of the three male-active networks (low-level miRNA and high-level mRNA targets in male tissues) (Fig. 4A, B, and C) included the exonuclease mut-7 homolog, *NOZZLE, GAMYB*, auxin response factor 18 family members, cinnamoyl-CoA reductase 2, myosin-11, and evm.model.Chr12.1832.1. Specifically, *NOZZLE* and *GAMYB* were detected in all three networks. In contrast, at least two of the three female-active networks (low-level miRNA and high-level mRNA targets in female tissues) (Fig. 4D, E, and F) included squamosa promoter-binding-like protein gene *7* (*SPL7*), *SPL9, SPL16, SPL17*, lysine-specific demethylase (*JMJ25*), *BHLH25, PCS1*, MAR-binding filament-like protein 1-1 (*MFP1-1*), origin of replication complex subunit 1A (*ORCS-1A*), Fanconi anaemia group M protein homolog, and cucumisin, growth-regulating factor 6 (*GRF6*), *ORCS-1A*, ATP sulphurylase 1-chloroplastic, and *SOBIR1*.

**Fig. 4.**
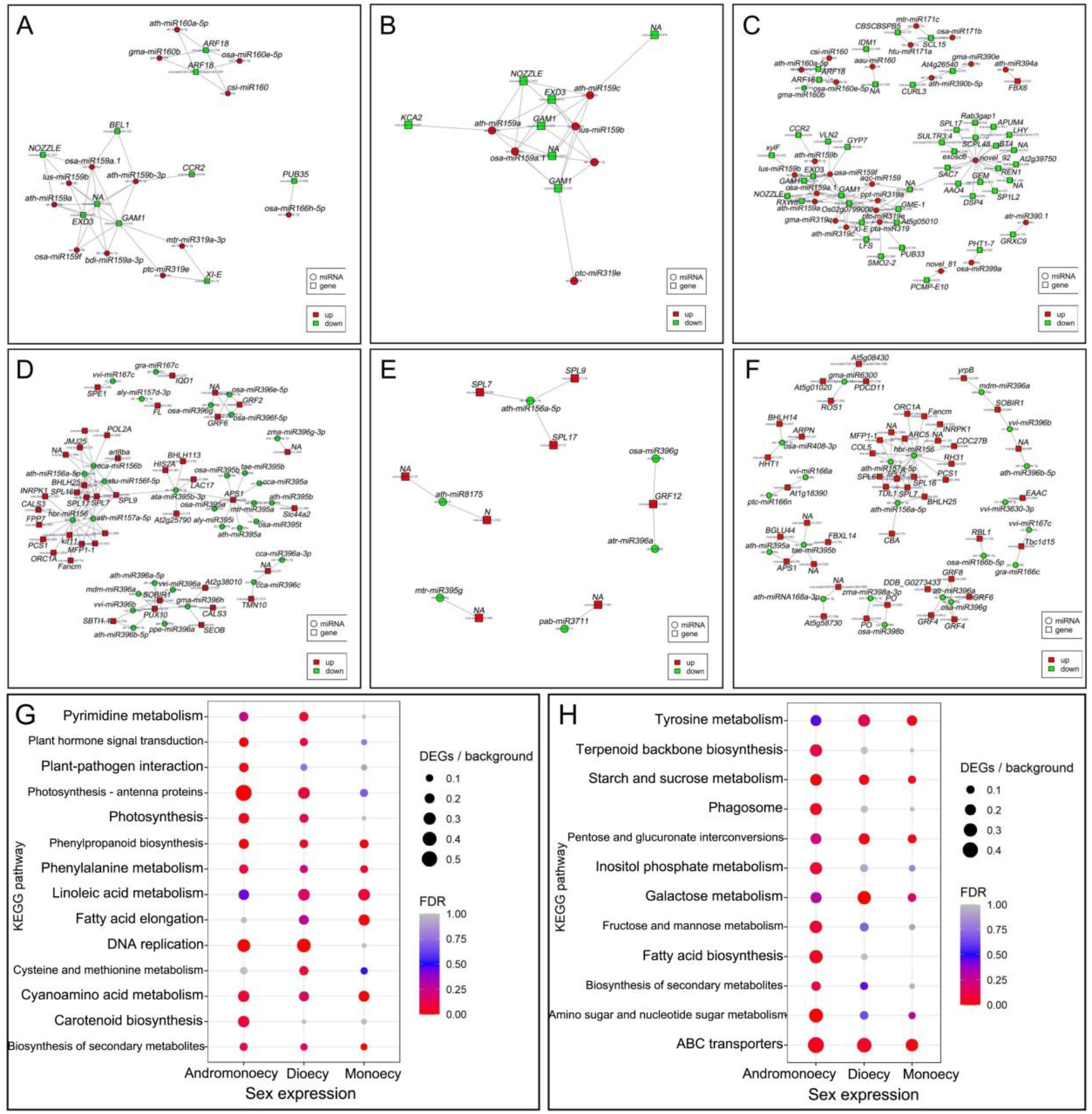
Networks constructed based on differentially expressed miRNAs and their mRNA targets, as well as KEGG pathway enrichments of differentially expressed genes. miRNAs with higher expression and their downregulated targets **(A)** in G_F compared with A_M, **(B)** in M_F compared with M_M, **(C)** in AM_H compared with AM_M, **(D)** in A_M compared with G_F, **(E)** in M_M compared with M_F, and **(F)** in AM_M compared with AM_H. KEGG pathway enrichments of **(G)** up- and **(H)** downregulated genes in female (or hermaphroditic) floral buds compared with male floral buds.

Comparisons of female (or hermaphroditic) and male tissues revealed that numerous KEGG pathways were commonly enriched in both single- and co-sex systems, including the female-active pathways phenylpropanoid biosynthesis, biosynthesis of secondary metabolites, and plant hormone signal transduction (Fig. 4G), as well as the male-active pathways starch and sucrose metabolism and galactose metabolism (Fig. 4H). Therefore, single- and co-sex systems use the same basic functional modules for flower sex determination.

### Population structure in *D. oleifera* according to sexual expression

We performed whole-genome sequencing of 150 *D. oleifera* individuals with a mean depth of 20×. After filtering, 3,545,359 SNPs and 318,863 indels were obtained for further analysis. The mean PI_HAT value was 0.056, indicating a low level of familial relationship in the population. PCA showed that the sampled individuals could be divided into three clusters (Fig. 5A). Single-sex plants and monoecious plants were found in all three clusters, whereas andromonoecious and androgynomonoecious trees were found only in groups 2 and 3. Pseudo-monoecious plants were found only in group 2, and they are distributed in close proximity. The results imply that the monoecious genetic factor prevails in *D. oleifera*, whereas andromonoecious, androgynomonoecious, and pseudo-monoecious individuals may be rare and develop only under certain conditions. The maximum-likelihood phylogenetic tree supported this notion; individuals obtained in close areas tended to have a close genetic relationship (Fig. 5B). The decay of LD with physical distance between SNPs occurred at 200 bp (r^2^ = 0.2) (Fig. S55A).

**Fig. 5.**
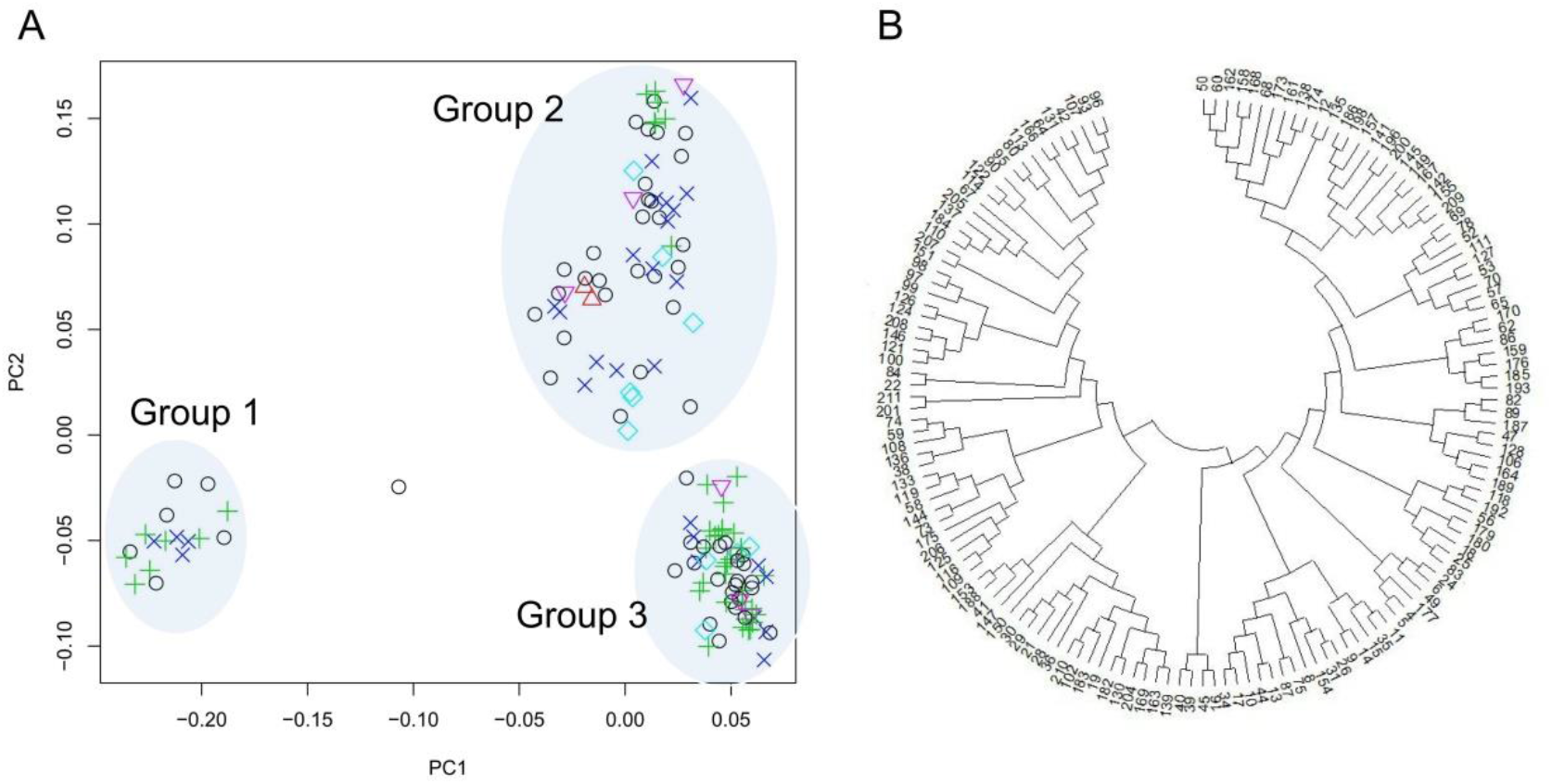
Phylogeny of the 150 *D. oleifera* plants. **(A)** Scatter clustering diagram based on the first two principal components after PCA of whole-genome sequence data. PC1 and PC2 explained 7.42% and 5.59% of the total variance, respectively. Black circle, gynoecy; red triangle, pseudo-monoecy; green +, androecy; blue cross, monoecy; light-blue diamond, androgynomonoecy; pink upside-down triangle, andromonoecy. **(B)** Maximum-likelihood phylogenetic tree of the 150 *D. oleifera* plants using MEGA-X labelled in order of sampling time. Therefore, similar numbers indicate similar distributions.

### GWAS of the co-sex phenotypes

To identify the genomic factors that confer the monoecious phenotype, we performed GWAS for the proportion of female shoots in genetically male plants. Ninety individuals with male flower production were selected; and 3,502,197 SNPs and 311,512 indels were retained for further analysis after LD pruning (Table S31).

GWAS for the proportion of female shoots (Table S13) detected a significant peak on chromosome 7 (Fig. 6A). Individuals with a heterozygous genotype at the locus with the strongest association signal showed greater proportions of female shoots (Fig. 6B; Table S32). A haplotype block spanning 29.0-29.4 Mb on chromosome 7 was strongly associated with phenotype (Fig. 6A). Seven genes of the *DUF247* family were identified in this block (Fig. 6C). Most were upregulated in female tissue compared with male tissue; one of these genes, evm.model.Chr7.983, was significantly upregulated in female floral buds compared with male floral buds in both monoecious and single-sex plants (Fig. 6D; Table S33). Most of the variants with the highest peak association were distributed upstream of evm.model.Chr7.983 (Fig. 6C), which may contribute to the differential expression of this gene. miRNA pab-miR3711, located within the block spanning 29.0-29.4 Mb, was downregulated in female floral buds compared with male tissues in monoecious plants (Table S34), possibly in relation to monoecious phenotype development.

**Fig. 6.**
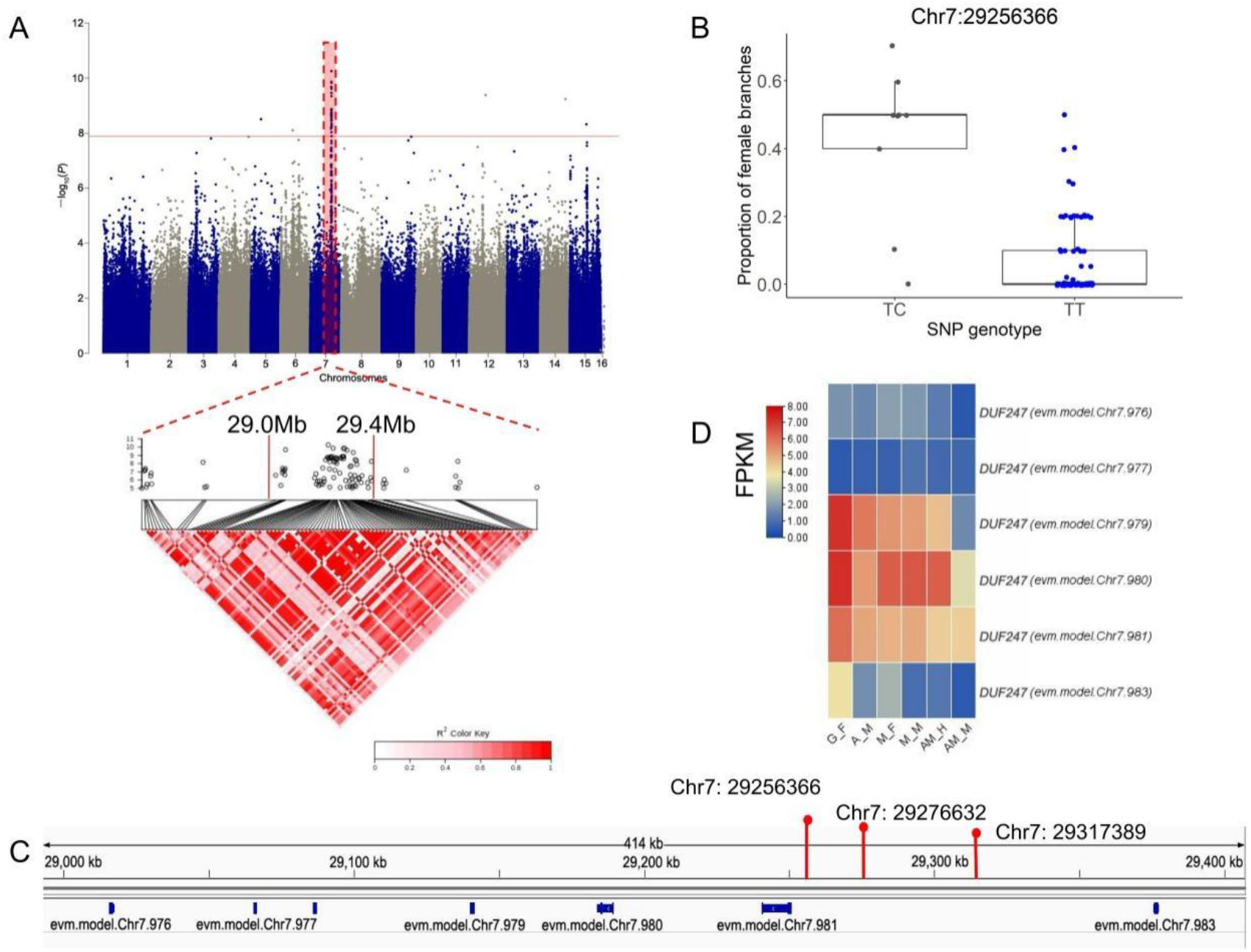
GWAS for the monoecious phenotype in *D. oleifera*. **(A)** Manhattan plot for the proportion of female shoots in 90 plants with male flower production (top), along with a local Manhattan plot and LD heatmap (bottom) of the associated region on chromosome 7. **(B)** Proportion of female shoots based on genotype at the most significant locus, Chr7: 29256366. **(C)** Schematic representation of gene position in the 29.0-29.4 Mb region of Chr7. **(D)** Expression pattern of genes in the 29.0-29.4 Mb region of Chr7 in female (or hermaphroditic) and male floral buds in single- and co-sex systems. Red line in **(A)** represents the Bonferroni-corrected *P*-value of 0.05, as shown in Figs. 7A and 8A.

We also recorded the proportion of hermaphroditic floral branches in the 90 plants. GWAS analysis for the proportion of hermaphrodite shoots revealed strong signals on chromosomes 2, 11, and 14 (Fig. 7A). According to the LD analysis, five, two, and three LD blocks with strong associations were detected on chromosomes 2, 11, and 14, respectively (Fig. S56). Individuals with heterozygous genotypes at the loci with the strongest association signals showed greater proportions of hermaphroditic shoots for all detected blocks (Fig. 7B and C; Table S35). In total, 65 genes were identified on the blocks, among which 3 and 2 genes were up- and downregulated in male floral buds compared with hermaphroditic floral buds in andromonoecious plants, respectively (Fig. 7D; Table S36). Additionally, genes encoding 31 lncRNAs (Table S37), 9 miRNAs (Table S38), and 39 circRNAs (Table S39) were identified in these regions. Further analysis of these sequences may identify genetic events linked to hermaphroditic flower development in genetically male *D*. spp.

**Fig. 7.**
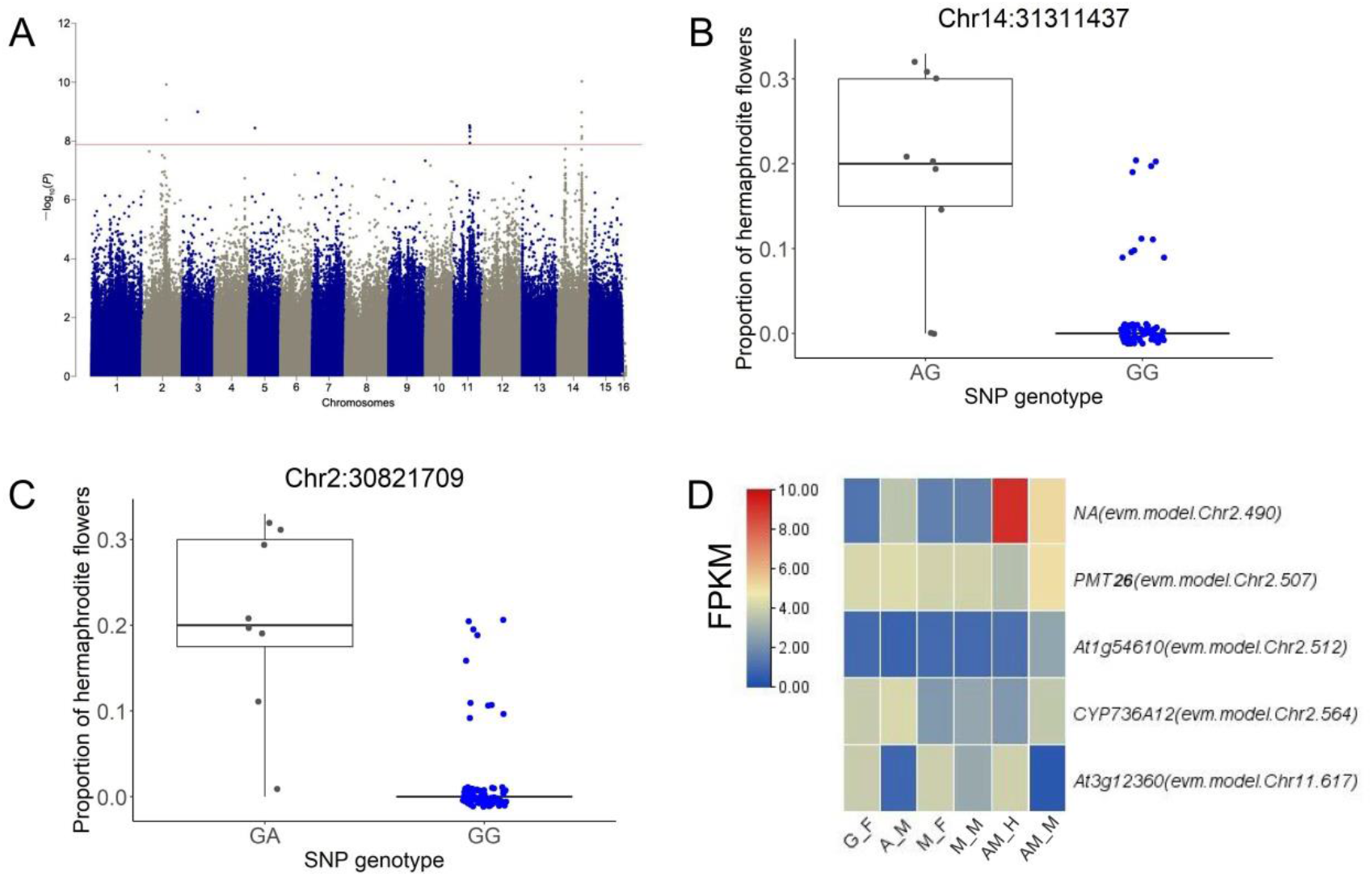
GWAS for hermaphroditic flower development in *D. oleifera*. **(A)** Manhattan plot for the proportion of hermaphroditic shoots among 90 samples with male flower production. **(B)** Proportion of hermaphroditic shoots based on genotype at the most significant locus, Chr14: 31311437, and **(C)** the second most significant locus, Chr2: 30821709. **(D)** Genes with significantly different expression in AM_M compared with AM_H in the association regions.

### Absence of the YY genotype in the *D. oleifera* population

The emergence of co-sex phenotype enables crossing among genetically male plants. A model-based analysis showed that the stability of dissolution of dioecy depends on the viability of the YY genotype (Crossman and Charlesworth, 2014), which is reduced by loss of function of Y-linked genes. Therefore, we characterised the sex-linked region. GWAS for the male expression (or *OGI*^+^) (Table S13) of 150 plants identified a sex-linked region at 22.0-32.0 Mb on chromosome 4 (Figs. 8A and S58; Table S40). This region corresponds to the male-specific region in *D. lotus* (Akagi *et al*., 2020) (Fig. 8C). The genotype of the SNPs with the strongest association signals in this region showed that most of the 54 gynoecious individuals (*OGI*^-^) were homologous (XX type), whereas most of the 89 male-functional individuals were heterozygous (XY type) (Fig. 8B; Table S41). Genotyping analysis indicated that the YY genotype was not supported by > 2 successive variant loci (Table S41). The occasional YY genotype may be attributed to recombination or genotyping error associated with the highly repetitive nature of this region. Therefore, the YY genotype is either present at a negligible level or absent from the population.

**Fig. 8.**
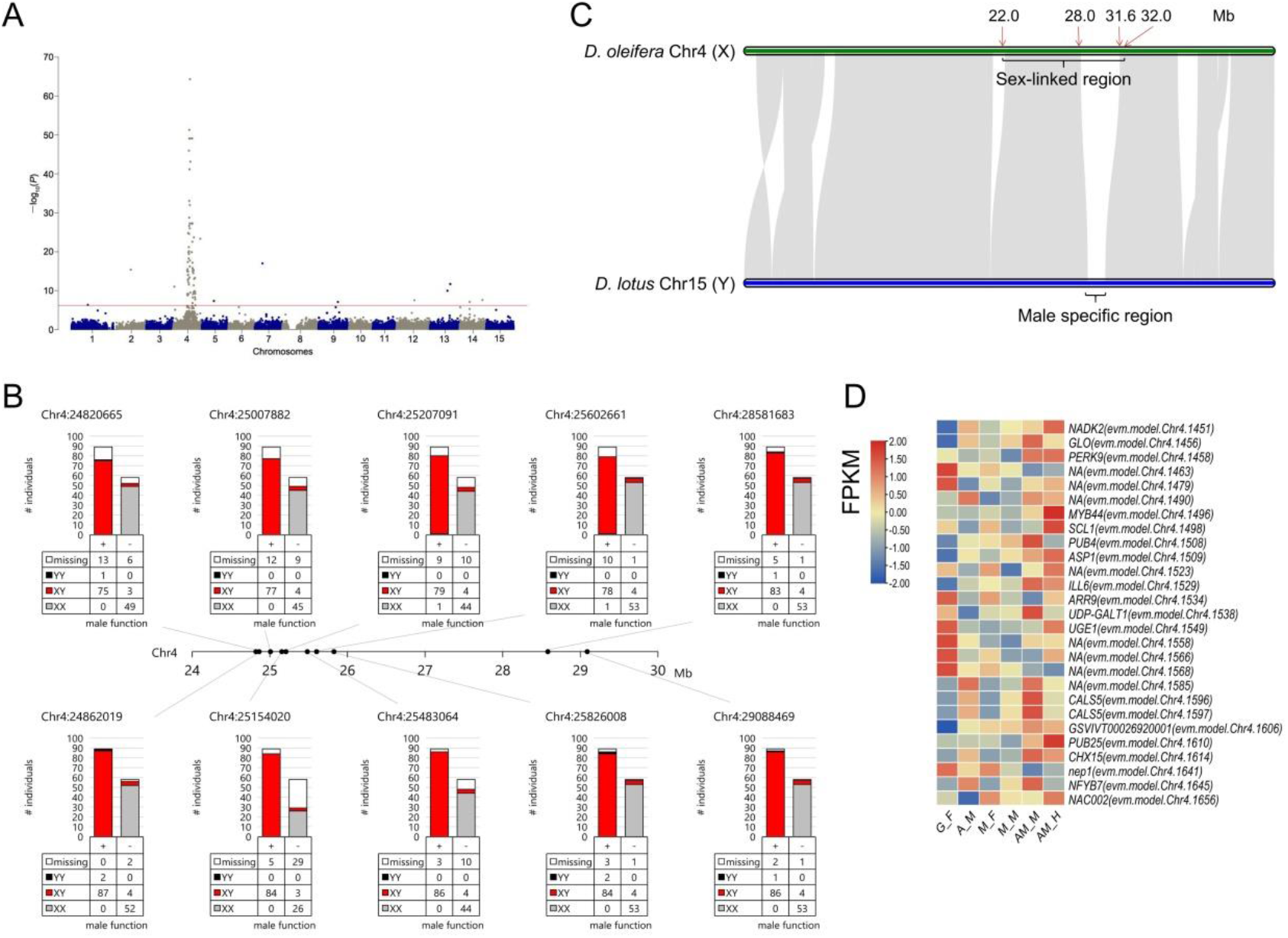
Genetic characterisation of the sex-linked region in the *D. oleifera* population. **(A)** GWAS for male expression. **(B)** Alignment of Chr4 of *D. oleifera* and Chr15 of *D. lotus*. **(C)** Genotype fractions of the SNPs with strong association signals in the sex-linked region. **(D)** Differentially expressed genes in the sex-linked region.

We detected a potential X-specific region and putative sex-related genes in this region, in addition to the male determinant *OGI* (Text S5; Table S42). Twenty-seven genes were differentially expressed between female (or hermaphroditic) and male floral buds in single- or co-sex systems, or both (Fig. 8D). Specifically, two genes, *GLO* and *ARR9*, which have masculinising functions in *Antirrhinum majus* (Perbal *et al*., 1996) and feminising functions in the genus *Populus* (Müller *et al*., 2020), respectively, were differentially expressed in this region (Fig. 8D).

## Discussion

### A sex-linked region contributing to sex dimorphism and diversity

All *D. oleifera* trees with male function, but exceptional one monoecious individual, had MSR and *OGI* (Table 1), and they showed many heterozygous genotypes in the sex-linked region on chromosome 4 (Fig. 8), consistent with the male heterogametic (XY) system. *smMeGI* and *MeGI* transcript analyses implied that the *OGI*/*MeGI* system (Akagi *et al*., 2014) functions in the sex determination of single-sex *D. oleifera* plants (Fig. 2).

In the *D. oleifera* population, two types of co-sex systems were identified: monoecy and hermaphroditic flower formation. The genotyping results (Table 1) indicate that all individuals can be divided into two groups: a genetically female (XX) group that includes gynoecious and pseudo-monoecious plants, and a genetically male (XY) group that includes androecious, monoecious, androgynomonoecious, and andromonoecious plants. Thus, the co-sex types (monoecious, androgynomonoecious, and andromonoecious) may represent evolutionary results of the breakdown of dioecy, where androecious trees acquire gynoecia development functions as demonstrated empirically in various species, including grape and papaya (Crossman and Charlesworth, 2014; Kafer *et al*., 2017; Cossard *et al*., 2021; Massonnet *et al*., 2020; Van Buren *et al*., 2015).

The lack of YY individuals in this study (Fig. 8) was surprising for us because it was inconsistent with the potential for crossing among genetically male individuals. This finding may be attributed to the low viability of the YY genotype (*i*.*e*., genetic degeneration). Genetic modelling has shown that under such conditions, stable coexistence of single- and co-sex plants in a population (subdioecy) can be achieved (Crossman and Charlesworth, 2014). In the present study, genomic and transcriptional analyses identified putative Y-/X-specific regions (Fig. 8C), as well as several genes in the sex-linked region that are differentially expressed between female and male tissues and may function in sexual expression, such as *GLO* and *ARR9* (Text S5); these findings imply functional divergence of the X and Y chromosomes. *GLO* is reportedly essential for stamen development in *Antirrhinum majus* (Perbal *et al*., 1996), whereas *ARR* proteins function in gynoecia development and are regarded as master regulators of sex expression in the genus *Populus* (Müller *et al*., 2020). The sex-linked region and the essential genes within that region presumably cooperate with the *OGI*/*MeGI* system to regulate the sex differentiation and diversity of *D*. spp.

### Feminising mechanisms that contribute to the dissolution of dioecy in co-sex *D. oleifera*

We identified three potential feminising mechanisms in both single- and co-sex systems: increased expression of the feminising gene *MeGI* and decreased abundance of *smMeGI*; genome-wide decrease in methylation levels; and sexual distinct regulatory networks of smRNAs and their targets. However, the first mechanism was inconsistent in the monoecious system; *MeGI* expression and *smMeGI* accumulation were not significantly different between M_F and M_M (Fig. 2). This is inconsistent with the pattern in monoecious hexaploid *D. kaki*, which has lower *smMeGI* levels and higher *MeGI* levels in the female flower buds of genetically male plants (Akagi *et al*., 2016a). Considering the similar *MeGI* expression in monoecious (both male and female floral buds) and androecious (male floral buds) plants (Fig. 2F), monoecious *D. oleifera* may develop gynoecia independently of *MeGI* regulation.

The mechanism that underlies gynoecia development in monoecious plants may be global regulation of DNA methylation (Fig. 3). Treatment of male flower buds with the DNA methylation inhibitor zebularine and 5-azacytidine induces pistil development and reduces pollen fertility in some *D. kaki* cultivars (Akagi *et al*., 2016a; Wang *et al*., 2022). The DNA demethylase gene *ROS1* (Gong *et al*., 2002) was upregulated in hermaphroditic floral buds compared with male floral buds in andromonoecious *D. oleifera* (Table S22), which may explain the genome-wide decrease in methylation level in hermaphroditic floral buds compared with male floral buds.

We also identified several putative key relationships between miRNA expression and target expression. Decreased expression of *GAMYBs* (evm.model.Chr13.1162 and evm.model.Chr11.1032) and increased expression of putative miRNA regulators were identified in female (or hermaphroditic) floral buds compared with male floral buds. *GAMYB* is a *trans*-activator of GA signalling (Gubler *et al*., 1995; Haseneyer *et al*., 2008) and functions in flower development (Gocal *et al*., 1999; Murray *et al*., 2003). The upregulation of *GAMYB* in male floral buds implies that GA signalling promotes the development of androecious tissues, consistent with our previous findings that GA promotes the male function in monoecious (Sun *et al*., 2017; Li *et al*., 2019) and andromonoecious (Li *et al*., 2021) *D. kaki*. Furthermore, *SPL* family genes (evm.model.Chr14.920, evm.model.Chr7.129, evm.model.Chr7.171, and evm.model.Chr12.765), *JMJ25*, and *GRFs* were commonly activated in female tissues, presumably through miRNA regulation. *SPL* family genes have diverse functions in plant development (Preston and Hileman, 2013); one of these genes acts a direct upstream activator of *LEAFY, FRUITFULL*, and *APETALA1* to control the timing of flower formation (Yamaguchi *et al*., 2009). *JMJ25* is a histone H3K9 demethylase gene that reportedly affects DNA methylation (Miura *et al*., 2009; Fan *et al*., 2018). *GRFs* are plant-specific transcription factors, and the *miR396/GRF* regulatory network is required for the proper development of the pistil in *Arabidopsis* (Omidbakhshfard *et al*., 2015; Liang *et al*., 2014). The functions of these genes in model plants and their expression patterns in *D. oleifera* were consistent with the working hypothesis regarding sexual expression in *D. oleifera*.

### Genetic factors linked to the dissolution of dioecy

The monoecious phenotype in *D. oleifera* was unique and could not be explained by known mechanisms. Therefore, we evaluated the genetic mechanisms that underlie the sexuality of monoecious and andromonoecious types. The candidate region for the monoecious trait on chromosome 7 included a cluster of seven *DUF247* genes (Fig. 6). The Y-specific dominant female suppression gene, *SOFF*, in dioecious *Asparagus officinalis* is a member of this gene family (Harkess *et al*., 2017). In *A. officinalis*, knockout of the *SOFF* gene converts males to hermaphrodites, knockout of the Y-specific male-promoting *aspTDF1* converts males to neuters, and knockout of both *TDF1* and *SOFF* converts males to females (Harkess *et al*., 2020). A *DUF247* family gene was identified as a male component of the self-incompatibility *S*-locus in perennial ryegrass (Manzanares *et al*., 2016; Thorogood *et al*., 2017). We detected clusters of duplicated *DUF* family genes in at least five loci in the *D. oleifera* genome (data not shown), implying functional divergence. The monoecious determinant, as well as its molecular genetic control, must be identified in subsequent studies. Although a very strong signal was obtained for chromosome 7, it could not explain all monoecious phenotypes (Fig 6B), implying that other loci and environmental factors affect female development.

The development of hermaphroditic flowers in dioecious systems because of mutations at the sex-determining locus has been observed in grape and papaya (Valleau, 1915; Massonnet *et al*., 2020; Van Buren *et al*., 2015). In contrast, our GWAS approach for the differentiation of hermaphroditic floral buds yielded candidate regions on chromosomes 2, 11, and 14, but not on chromosome 4 (which has the sex-linked region). We also observed decreased expression of the feminising gene *MeGI* in hermaphroditic flower buds (Fig. 2F). Therefore, the establishment of hermaphroditic flower development in *D. oleifera* is independent of direct activation/inactivation of the genetic regulation of sex dimorphism through the existing *OGI*/*MeGI* system, as implied in work regarding *D. kaki* (Masuda *et al*., 2022; Li *et al*., 2021).

Masuda *et al*. (2022) reported that *DkRAD* regulates gynoecia formation in hermaphroditic flowers of hexaploid *D. kaki*. This may also function in *D. oleifera* because the expression of the *DkRAD* homologue was higher in female and hermaphroditic tissues than in male tissues (Fig. S59), as in *D. kaki*. One discrepancy compared with the work of Masuda *et al*. (2022) is that our study revealed many diploid *D. oleifera* plants bearing hermaphroditic flowers, whereas Masuda *et al*. regarded the hermaphrodite mechanism in *Diospyros* as polyploid species-specific. Our results indicate that the evolution of hermaphroditic flower development in *Diospyros* is not ploidy-dependent; however, considering the similarity in sexual expression between *D. kaki* and *D. oleifera*, as well as their close phylogenetic relationship (Fu *et al*., 2017), a common evolutionary event and mechanism presumably led to sex expression diversity in both species. Further genetic analyses of sex expression in *D. kaki* and *D. oleifera* are needed.

### Summary and future perspectives

Based on our findings for *D. oleifera*, female shoots mainly develop from mixed dormant buds that developed on the tips of the flowering mother branches in monoecious plants which are genetically male with *OGI*. In contrast, male shoots mainly develop on the basal parts of the mother branches (Fig. 9). Floral primordia in *D. oleifera* are initiated in early summer, then experience a long dormant period until the following spring to break the buds (Li *et al*., 2016). Therefore, dormant floral buds on the tips of the flowering mother branches are more likely to activate the feminising scenario under natural conditions (Fig. 9). The same pattern was observed in cultivated hexaploid *D. kaki* (Fig. S60). Therefore, we recommend an investigation of the physiological and molecular statuses of floral buds on the top and basal parts of flowering mother branches in co-sex individuals at the beginning of flower bud differentiation.

**Fig. 9.**
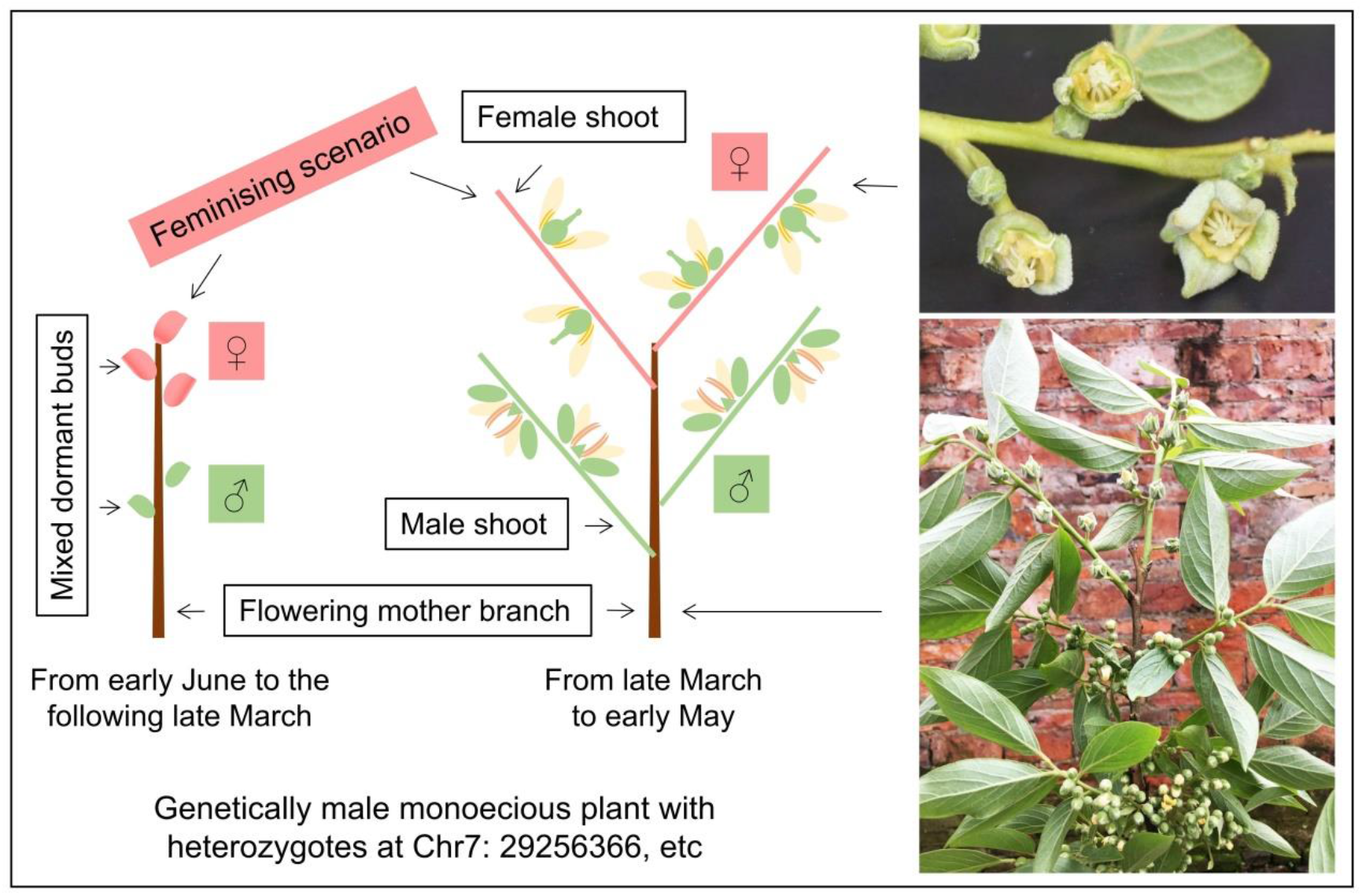
Schematic of the feminising scenario in monoecious *D. oleifera*.

## Acknowledgements

This study was supported by the National Key R&D Program of China (2018YFD1000606 and 2019YFD1000600) to JF and PS, the National Natural Science Foundation of China (32071801), and the Fundamental Research Funds for the Central Non-profit Research Institution of CAF (CAFYBB2017ZA005) to PS, and JSPS KAKENHI (21KK0269) to SN and (20K20454) to SN and RT. We thank Novogene Bioinformatics Institute for the technical assistance.

## Author contributions

PS, SN, RT, FL and JF designed the study; PS, HL, YM and WH surveyed and collected the plant materials; PS, SN, HL and YM performed the experiments and analyzed the data with help from YS, SD, YW, JY and YZ; CL and HD assembled the genome; PS and SN drafted the manuscript, RT, JF and CL revised the manuscript.

## Data availability

The repositories and accession numbers of raw sequence data can be found in the Supplementary Method.

