## Supplementary Method for "Molecular and genetic mechanisms conferring dissolution of dioecy in *Diospyros oleifera* Cheng"

#### Method S1: Ploidy determination for *D. oleifera* by flow cytometer

About 20–50 mg of fresh tissue were chopped using a razor blade in 1 ml of LB01 buffer (15 mM Tris, 2 mM EDTA-$Na_2$, 0.5 mM spermine tetrahydrochloride, 80 mM KCl, 20 mM NaCl, 0.1% (vol/vol) Triton X-100; pH 7.5 adjusted with NaOH; 15 mM 2-mercaptoethanol). The tissue was collected by gentle pipetting and filtered through a 400 μm mesh nylon strainer. The samples were stained with 100 μg/ml Propidium Iodide (PI) simultaneously with 100 μg/ml RNase in an ice bath for 10 min before the analysis.

The nuclear fluorescence was measured using the MoFlo XDP high-speed flow cytometer (Beckman Coulter Inc., USA) with a 70 μm ceramic nozzle at 60 psi sheath pressure. PI fluorescence was excited with a solid-state laser (488 nm) and detected with a 625/26 nm HQ band-pass filter. The FL3-Height/SSC-Height gate was used to eliminate debris, cell fragments, and dead cells. Single- and double-nuclei were discriminated using FL3-Height /FL3-Area.

#### Method S2: A BioNano optical mapping-assisted high-quality chromosome-level genome assembly of *D. oleifera*

##### S2.1 Plant materials and Illumina, PacBio sequencing

Genomic DNA was extracted from fresh leaves of a female *D. oleifera* using a DNAsecure Plant Kit (Tiangen Biotech, Beijing, China). A total of 104.02 Gb of Illumina raw sequence data (155.28-fold coverage of the *D. oleifera* genome) and 99.76 Gb of PacBio raw sequence data (148.90-fold coverage of the *D. oleifera* genome) from Suo *et al*. (2020) were used for genome assembly.

##### S2.2 BioNano library construction and sequencing

Approximately 2-3 g of young fresh leaf obtained from the same *D. oleifera* tree was transferred to a 50 ml BD tube and ground to a fine powder in liquid nitrogen. The sample was filtered through 100 μm and 40 μm diameter membranes. Using a density-gradient buffer, the nuclei in the interlayer were collected and embedded in a 60 μl low-melting-point gel. The embedded nuclei were digested overnight in 250 μl protease K, followed by digestion in 50 μl of RNase A for 1 h. DNA was extracted from the samples using BioNano Plant Tissue DNA Isolation Kit, followed by purification, and quantification with a Qubit 3.0 fluorometer. Approximately 600 ng genomic DNA was subjected to nicking (with the restriction enzyme DLE-1), labeling, repairing, and staining reactions, and incubated at room temperature for 30 min. After thorough mixing, the DNA concentration was quantified. The labeled library was loaded onto a Saphyr Chip for sequencing, which generated ~177Gb of high-quality BioNano molecules.

**S2.3 Hi-C library construction and sequencing**

A total of 98.24 Gb of the Dovetail Hi-C library reads, which were reported by Suo *et al.* (2020), were used for genome assembly in this present study.

**S2.4 Evaluation of genome size and heterozygosity**

The genome size was evaluated based on the *K*-mer distribution, as determined using Jellyfish ((http://www.genome.umd.edu/jellyfish.html, v2.2.3) with the following parameters: -C –m 17 –s 200M. The following equation was used to calculate genome size: Genome size = *K*-mer coverage/Mean *K*-mer depth. The heterozygosity ratio for the *D. oleifera* genome was estimated using GenomeScope (http://qb.cshl.edu/genomescope/).

**S2.5 Genome assembly and polishing**

The raw SMRT reads were self-corrected and assembled with CANU (genomeSize = 700m, min OverlapLength = 600, minReadLength = 1000, v1.7) (Koren *et al.*, 2017). The BioNano data were assembled with RefAligner and Assembler (https://bionanogenomics.com/) and used to generate a CANU assembled contigs. The gaps in the hybrid scaffold were filled using Highly Efficient Repeat Assembly (HERA) (Du and Liang, 2019) to form longer scaffolds with self-corrected reads under the following parameters: InterIncluded_Side = 30000, InterIncluded_Identity = 99; InterIncluded_Coverage = 99; MinIdentity = 97; MinCoverage = 90; MinLength = 5000; MinIdentity_Overlap = 97; MinOverlap_Overlap = 1000; MaxOverhang_Overlap = 100; MinExtend_Overlap = 500. Longer contigs were further connected using HERA with the same parameters as above.

##### S2.6 Heterozygosity filtration depending on similarity among the final contigs and Hi-C

Self-mapping among the final contigs was conducted by Burrows-Wheeler Aligner (Bwa) mem (Li and Durbin, 2010). Mapping similarity over 96% and matching lengths longer than 2 kb were documented. Then the coverage ratio between the two contigs was calculated. One of the two contigs with a coverage ratio higher than 99% was filtered out. The contigs, whose coverage ratio was lower than 99% and coverage region was longer than 30 kb, were combined according to the overlapping sequences. After filtration, the Hi-C sequencing data were aligned to the remaining genomic sequences to filter the heterozygous regions with a Hi-C heat map, resulting in *D.oleifera* main genome.

##### S2.7 Chromosome construction with Hi-C sequencing data

The Hi-C reads were aligned to the main genome with Juicer

(https://github.com/aidenlab/juicer). The genome was clustered, ordered, and oriented onto chromosome-level using 3D-DNA (Dudchenko *et al*., 2017).

**S2.8 Error correction for *D. oleifera* genome by Pilon**

The Illumina reads were aligned to the main genome and heterozygous sequences. The reads with aligned scores below Q30 were filtered by SAMtools (Li *et al*., 2009). Then Pilon (https://github.com/broadinstitute/pilon, v1.22) with default parameters was used to perform the error correction, resulting in the final 15 chromosomes with 89 gaps. Then the heterozygous sequences were aligned to the main genome to determine whether the heterozygous regions were well separated.

**S2.9 Assessment of completeness of the *D. oleifera* genome**

The completeness of the assembled genome was assessed using Benchmarking Universal Single-Copy Orthologs (BUSCO) (Simão *et al*., 2015). The Illumina reads reported by Suo *et al*., 2020 were also aligned to the *D. oleifera* main genome by Bwa mem to evaluate the completeness.

**S2.10 Repetitive elements identification and annotation of *D. oleifera* genome**

Repetitive elements identification, protein-coding genes, functional annotation, and non-coding RNAs annotation of the *D. oleifera* main genome were conducted following Suo *et al*. (2020).

**S2.11 Synteny analysis**

The current *D. oleifera* main genome, the version reported by Suo *et al*. (2020), and the *D. lotus* genome (Akagi *et al*., 2020) were aligned using Mummer (http://github.com/gmarcais/mummer).

**S2.12 Whole-genome duplication and macrosynteny analysis**

We used BLASTP (E-value < 1E−5) to perform homolog and paralog searches with *D. oleifera* and other genomes (*A. chinensis, C. canephora, C. sinensis*), and MCScanX (s = 5, e = 1e−5) (Wang *et al.*, 2012) was used to detect syntenic blocks. Then, transversion substitutions at 4-fold degenerate sites (4dtv) rates for all syntenic genes were calculated to identify putative whole-genome duplication (WGD) or species split events in *D. oleifera*.

#### Method S3: Assembly and annotation of male-unmapped sequences

Resequencing reads of 14 androecious, 4 andromonoecious, 15 monoecious, 6 androgynomonoecious, and 2 pseudo-monoecious *D. oleifera* individuals were mapped to the main *D. oleifera* reference genome constructed with Method S2 using the Bwa mem option and the paired-end model (Li and Durbin, 2010). Unmapped sequences were extracted using SAMtools (Li *et al.*, 2009), and assembled using SoapDenovo (Li *et al.*, 2010). The resultant contigs were referred to as male-unmapped sequences in the main text and below. Repetitive elements identification, protein-coding genes, functional annotation, and non-coding RNAs annotation of the male-unmapped sequences were conducted following Suo *et al.* (2020).

#### Method S4: Checking the existence of *Kali* on the 5' UTR region of *OGI* in *D. oleifera*

The primer sequences (5'-3', F: ACATCCAAAGTTCTGGAGAATCA R: ATTGGTGCTTGGTCAAACATATC) and KOD One™ PCR Master Mix (TOYOBO, Japan) were used to clone the 5' UTR region of *OGI* in *D. oleifera* and *D. kaki*.

#### Method S5: Plant tissues sampled for methylome and whole transcriptome analyses

Floral buds were sampled in natural populations in Guangxi Zhuang Autonomous Region, China (Fig. S18), in mid-April (April 15~17th). Additionally, stem tissues of immature flowering shoots, which exclusively bore female or male floral buds, were also sampled. Those samples were snap-frozen in liquid nitrogen and stored at −80 ℃ until DNA and RNA isolation for the whole-genome bisulfite sequencing (BS-seq) and whole transcriptome sequencing, respectively.

The flower differentiation and floral bud development in *Diospyros*, mainly in *D. kaki*, have been well characterized, and key stages for sex determination have been identified. Our previous observation indicated the development process of *D. oleifera* floral buds was synchronous to that of *D. kaki*. The development of floral primordia is initiated in early summer (mid-June in the experiment area) in both female and male floral buds of *D. kaki*. Then floral buds experience a long dormancy period until the following spring (March). The development of pistil and stamen primordia in both female and male floral buds is initiated at the bud break stage (late March). Subsequently, the arrest of stamen primordia leading to the formation of female flowers occurred after bud break (mid-April), while the arrest of pistil primordia leading to male flowers also occurred at this time (Li *et al.*, 2016). If neither pistil nor stamen primordia were arrested, the hermaphroditic flowers would develop (Li *et al.*, 2021). This result suggests that the developmental stage in mid-April is essential for the final sex expression of a flower, which is further supported by the facts: (1) accumulation levels of small RNAs (smRNA) on *MeGI* (*smMeGI*) were highest in early and late April compared with other stages in both *D. lotus* and *D. kaki* (Akagi *et al.*, 2016); (2) the largest divergence of *MeGI* expression between female and male floral buds in *D. kaki* was observed in mid-April (Li *et al.*, 2019).

### Method S6: Methylome detection

##### S6.1 Extraction of DNA and Bisulfite sequencing (BS-Seq)

Genomic DNA was extracted from floral bud and stem tissues collected in mid-April (Table. S14) using a DNeasy Plant Mini kit (QianGen, Shanghai, China) following the manufacturer's instructions and then checked the quality with 0.1% agarose gels and a NanoPhotometer spectrophotometer (Implen, Westlake Village, CA, USA). DNA concentrations were measured using Qubit DNA Assay kit on a Qubit 2.0 fluorometer (Life Technologies, Carlsbad, CA, USA).

Next, 5.2 μg of genomic DNA spiked with 26 ng of lambda DNA (negative control) was fragmented by sonication to 200-300 bp with Covaris S220 ultrasonicator, followed by end-repair, adenylation, and methyl-treated adapter ligation. Then these DNA fragments were treated twice with bisulfite using EZ DNA Methylation-Gold kit (Zymo Research, Irvine, CA, USA) to transform non-methylated cytosines into uracil, before the resulting single-strand DNA fragments were PCR amplificated using KAPA HiFi HotStart Uracil + ReadyMix (2×). Library concentration was quantified by Qubit 2.0 Fluorometer (Life Technologies) and quantitative PCR, and the insert size was assayed on Agilent Bioanalyzer 2100 system. Sequencing of the BS-seq library, which generated 150-bp paired-end reads, was performed on an Illumina NovaSeq 6000 platform followed by Illumina CASAVA pipeline analysis.

##### S6.2 Quality Assessment of Sequencing Data

The sequenced paired-end raw reads were checked for quality using FastQC (fastqc_v0.11.5) and stored in the FASTQ file format (Babraham Bioinformatics, http://www.bioinformatics.babraham.ac.uk/projects/fastqc/). The resulting files were pre-processed with Trimmomatic v0.36 software (Bolger *et al*., 2014) using the following parameters: SLIDINGWINDOW: 4:15; LEADING:3, TRAILING:3; ILLUMINACLIP: adapter.fa: 2:30:10; and MINLEN:36. Reads passing these filtering steps were counted as clean reads for use in subsequent analyses. Finally, basic quality statistics on the clean reads were obtained using FastQC.

##### S6.3 Reference Data Preparation and BS-seq data analysis

Here, we used the combined *D. oleifera* genome, including the *D. oleifera* main genome, male-unmapped sequences, and the Chloroplast genome, as reference. Then, we performed reverse complementation process (C to T, and G to A) using Bismarkv 0.16.3 software (Krueger and Andrews, 2011) and Bowtie2 (Langmead and Salzberg, 2012). In addition, a gene annotation file in gene transfer format, a Gene Ontology (GO) annotation file, a description file, and a gene region file in browser extensible data format were generated for subsequent annotation and function analyses.

The quality-checked clean reads generated by BS-seq were aligned against the two strands of the converted reference genome. The best unique alignment of these sequence reads was selected from the two sets of pairwise comparisons. To infer cytosine methylation states and positions, the sequences were then compared against the original genomic sequence. Identical sequences aligned to a unique genomic region were regarded as duplicates and used to estimate sequencing depth and coverage. To allow their visualization in the IGV browser, sequences were transformed into bigwig format (non-overlap) (Krueger and Andrews, 2011; Wang *et al*., 2014). The bisulfite non-conversion rate was defined as the number of sequenced cytosines at all of the cytosine reference positions divided by the number in the lambda genome.

**S6.4 Estimating methylation level**

To identify the methylation site, we modeled the sum mC of methylated counts as a binomial (Bln) random variable with methylation rate r: mC ~ Bln (mC+umC*r). In order to calculate the methylation level of the sequence, we divided the sequence into multiple bins, with a bin size of 10 Kb. The sum of methylated and unmethylated read counts in each window was calculated. Methylation level (ML) for each window or C site shows the fraction of methylated Cs, and is defined as: ML(mC) = reads(mC) / (reads(mC)+reads(C)). Calculated ML was further corrected with the bisulfite non-conversion rate according to previous studies (Lister *et al.*, 2013). Given the bisulfite non-conversion rate r, the corrected ML was estimated as: $ML_{(corrected)}$ = (ML(mC) - r) / (1 - r). Methylcytosine sequence contexts—mCG, mCHG, and mCHH (where H represents A, T, or C)—were analyzed. Methylation level densities and methylcytosine distributions in each chromosome and gene functional region (promoter, exon, intron, and 2-kb upstream and downstream regions) were also analyzed (Zhong *et al.*, 2013; Krzywinski *et al.*, 2009; Lister *et al.*, 2009). Differences in global methylation levels and methylcytosine distributions in gene structural regions (including 2-kb upstream and downstream) were compared in the comparative combinations of M_F versus (vs.) M_M, G_F vs. A_M, G_S vs. A_S, and AM_M vs. AM_H (Table S14) (Song *et al.*, 2013).

**S6.5 Correlation Analysis and DMR Detection**

Hierarchical clustering was conducted based on PCA analysis for the methylation levels of CG, CHG, and CHH subcontexts among all the samples. Differentially methylated regions (DMRs) were identified using the DSS software (Feng *et al.*, 2014; Wu *et al.*,

2015; Park and Wu, 2016) in the comparative combinations of M_F versus (vs.) M_M, G_F vs. A_M, G_S vs. A_S, and AM_M vs. AM_H(**Table S14)**. Information from neighboring cytosine sites (i.e., spatial correlation) and site read depths were analyzed to improve the accuracy of long cytosine reads. Variance among biological replicates was analyzed using a beta-binomial distribution model. DMRs were annotated, and DMR-related genes were defined as those having coding regions (from the transcriptional start site (TSS) to the transcriptional end site (TES)) or promoter regions (i.e., upstream 2-kb from the TSS) that overlapped with DMRs.

##### S6.6 GO and KEGG Enrichment Analysis of DMR-related Genes

GO enrichment analysis of DMR-related genes was performed using the GOseq R package (Young *et al.*, 2010). The resulting *P*-values were adjusted using Benjamini and Hochberg's approach for controlling the false discovery rate. A GO term was considered to be significantly enriched in DMR-related genes at a corrected *P*-value < 0.05. Statistical enrichment of KEGG pathways in DMR-related genes was tested using KOBAS software (Mao *et al.*, 2005).

#### Method S7: Long non-coding RNAs (lncRNAs) and mRNAs analyses

##### S7.1 RNA Isolation, quantification, and qualification

RNA was isolated from the samples shown in Table S15 using an RNeasy Plant Mini kit (QianGen). Extracted RNA was checked for degradation and contamination using 1% agarose gels and a NanoPhotometer spectrophotometer (Implen, Munich, Germany), respectively, and then quantified with a Qubit RNA Assay kit on a Qubit 2.0 fluorometer (Life Technologies). RNA integrity was assessed on a Bioanalyzer 2100 system (Agilent) using the supplied RNA Nano 6000 assay kit.

#### S7.2 Library preparation and sequencing

Three μg of total RNA per sample was used as input material for the RNA library preparations. Firstly, ribosomal RNA was removed by Epicentre Ribo-zero™ rRNA Removal kit (Epicentre, USA), and the rRNA-free residue was cleaned up by ethanol precipitation. Subsequently, sequencing libraries were generated using the rRNA-depleted RNA by NEBNext Ultra™ Directional RNA Library Prep kit for Illumina (NEB, USA) following the manufacturer's recommendations. Briefly, fragmentation was carried out using divalent cations under elevated temperature in NEBNext First Strand Synthesis Reaction Buffer (5×). First-strand cDNA was synthesized using random hexamer primer and M-MuLV Reverse Transcriptase, RNaseH minus (Promega). Second strand cDNA synthesis was subsequently performed using DNA Polymerase I and RNase H. In the reaction buffer, dNTPs with dTTP were replaced by dUTP. The remaining overhangs were converted into blunt ends via exonuclease/polymerase activities. After adenylation of 3' ends of DNA fragments, NEBNext Adaptor with hairpin loop structure was ligated to prepare for hybridization. In order to select cDNA fragments preferentially 250-300 bp in length, the library fragments were purified with the AMPure XP system (Beckman Coulter, Beverly, USA). Then 3 μl USER Enzyme (NEB, USA) was used with size-selected, adaptor-ligated cDNA at 37°C for 15 min followed by 5 min at 95°C before PCR. Then PCR was performed with Phusion High-Fidelity DNA polymerase, Universal PCR primers, and Index Primer. At last, products were purified with AMPure and library quality was assessed on the Agilent Bioanalyzer 2100 system. The libraries were sequenced on an Illumina NovaSeq 6000 platform and 150 bp paired-end reads were generated.

##### S7.3 Bioinformatics identification of lncRNAs

The pipeline used for the identification of lncRNAs is described in Fig. S32A. Clean data were obtained by removing reads containing adapters, reads containing poly-N, and low-quality reads from the raw data. The clean data of high quality mapped to the reference genome. An index of the reference genome was built using Bowtie v2.0.6 (Langmead *et al*., 2009) and paired-end clean reads were aligned to the reference genome using TopHat v2.0.9 (Trapnell *et al*., 2009). After the alignment, the mapped reads of each sample were assembled by both Scripture (beta2) (Guttman *et al*., 2010) and Cufflinks (v2.1.1) (Trapnell *et al*., 2010) in a reference-based approach. Both methods use spliced reads to determine exon connectivity, but with two different approaches.

Next, transcripts were predicted to have coding potential by either one of the four tools (CNCI (v2), CPC (Kong *et al*. 2007), Pfam Scan (v1.3) (Bateman *et al*., 2004; Punta *et al*., 2012), and PhyloCSF (v20121028) (Lin *et al*., 2011)) were filtered out, and those without coding potential were used as candidate set of lncRNAs. Briefly, five steps were used to screen lncRNAs. Transcripts with low confidence levels and low expression levels were filtered out in Step 1. Transcripts shorter than 200 bp were filtered out in Step 2. Transcripts overlapping with exons in the database were filtered out using the software of Cuffcompare in Step 3. Transcripts with FPKM lower than 0.5 were filtered out using Cuffquant in Step 4. Transcripts with uncertain coding potential (TUCP) were classified as TUCPs in Step 5.

Cuffdiff (v2.1.1) (Trapnell, *et al*., 2010) was used to calculate the FPKMs of lncRNAs, coding genes and TUCPs in each sample. Cuffdiff provides statistical routines for determining differential expression in digital transcript or gene expression data using a model based on the negative binomial distribution. Differential expression levels were identified in the comparative combinations of M_F vs. M_M, G_F vs. A_M, G_S vs. A_S, and AM_M vs. AM_H (Table S15) using Cuffdiff (Trapnell, et al., 2010). The resulting *P*-values were adjusted using Benjamini and Hochberg's approach for controlling the false discovery rate. The corrected *P*-value < 0.05 were defined as differentially expressed.

**S7.4 Target gene prediction**

The cis-acting mechanism of lncRNAs involves them acting on neighboring target genes. To identify such lncRNAs, a search was performed for coding genes 10–100 kb upstream and downstream of the lncRNAs; the function of those identified was then analyzed. The trans-acting mechanism of lncRNAs to identify each other by the expression level. We clustered the genes from different samples to search for common expression modules and then analyzed their function through functional enrichment analysis. The GO and KEGG enrichment analysis of differentially expressed genes or lncRNA (and TUCPs) target genes were performed as described in Method S6.

**S7.5 Construction of the co-expression network**

Core mRNAs and lncRNAs networks associated with sex differentiation between female and male floral buds (Table S30) were conducted with the method reported by Langfelder and Horvath (2008). In co-expression networking, genes were represented by nodes, and the correlation values (weight) between two genes were calculated by raising the Pearson's correlation coefficient. The genes in the same module were visualized with the Cytoscape program (Shannon *et al*., 2003).

#### Method S8: Circular RNAs (circRNAs) analyses

##### S8.1 circRNAs identification, quantification and differential expression analyses

RNA isolation, library preparation, sequencing, and mapping to the reference genome for circRNAs analyses were conducted in accordance with Method S7. The circRNAs were detected and identified using find_circ (Memczak *et al*., 2013) and CIRI2 (Gao *et al*., 2015). Circos software was used to construct the circos figure. The expression level of circRNAs was evaluated as transcripts per million (TPM) (Zhou *et al*., 2010).

Differentially expressed circRNAs (DEcircRNAs) in the comparative combinations of M_F vs M_M, G_F vs A_M, G_S vs A_S, and AM_M vs AM_H (Table S15) were identified using the DESeq R package (1.10.1) with corrected *P*-value < 0.05 (as described in Method S7). The GO and KEGG enrichment analyses for host genes of differentially expressed circRNAs were implemented as described above. The host genes were defined as genes that produce circRNAs by reverse splicing.

##### S8.2 MicroRNA (miRNA) target site analysis

The miRNA binding sites of circRNAs, and the target coding genes of predicted target miRNAs were identified using psRobot_tar in psRobot (Wu *et al*., 2012) for plants. Cytoscape software was used to construct the circRNA-miRNA-gene networks.

#### Method S9: Small RNA (smRNA) analyses

##### Library preparation for smRNA sequencing

RNA isolation, quantification, and qualification were performed as mentioned in Method S7. Three μg of total RNA per sample was used as input material for the smRNA library. Sequencing libraries were generated using NEBNext Multiplex smRNA Library Prep Set for Illumina® (NEB, USA.) following the manufacturer's recommendations and index codes were added to attribute sequences to each sample. Briefly, NEB 3' SR Adaptor was directly, and specifically ligated to 3' end of miRNA, siRNA, and piRNA. After the 3' ligation reaction, the SR RT Primer hybridized to the excess of 3' SR Adaptor (that remained free after the 3' ligation reaction) and transformed the single-stranded DNA adaptor into a double-stranded DNA molecule. Next, 5' ends adapter was ligated to 5' ends of miRNAs, siRNA, and piRNA. Then first-strand cDNA was synthesized using M-MuLV Reverse Transcriptase (RNase $H^-$). PCR amplification was performed using LongAmp Taq 2× Master Mix, SR Primer for Illumina, and index (X) primer. PCR products were purified on an 8% polyacrylamide gel (100V, 80 min). DNA fragments corresponding to 140~160 bp (the length of small noncoding RNA plus the 3' and 5' adaptors) were recovered and dissolved in 8 μL elution buffer. At last, library quality was assessed on the Agilent Bioanalyzer 2100 system using DNA High Sensitivity Chips. The library preparations were sequenced on an Illumina NovaSeq 6000 platform and 50 bp single-end reads were generated.

**Analysis of smRNA sequencing data**

After the removal of the reads containing poly-N, with 5' adapter contamination, without 3' adapter or insert tag, and low-quality reads from the raw data, the remaining reads were filtered with the size range of 18-30 bp, and mapped to the *D. oleifera* reference genome by Bowtie (Langmead *et al*., 2009) without mismatch for further analysis. Rfam 11.0 (Burge *et al*., 2013) was applied to remove tags originating from protein-coding genes, repeat sequences, rRNA, tRNA, snRNA, and snoRNA. Then, mapped smRNA tags were used to look for known miRNAs. MiRBase20.0 was used as a reference, while modified software miRDeep2 (Friedländer, *et al*., 2012) and sRNA-tools-cli were used to obtain potential miRNAs and draw the secondary structures. The characteristic hairpin structure of miRNA precursors could be used to predict novel miRNAs. The software miREvo (Wen *et al.*, 2012) and miRDeep2 (Friedländer, *et al.*, 2012) were integrated to predict novel miRNAs by exploring the secondary structure, the Dicer cleavage site, and the minimum free energy of the smRNA tags unannotated in the previous steps. The expression levels were represented by TPM (Zhou *et al.*, 2010). Differential expression analysis in the comparative combinations of M_F vs M_M, G_F vs A_M, G_S vs A_S, and AM_M vs AM_H was performed using the DESeq R package (1.8.3). The corrected *P*-value of < 0.05 as mentioned in Method S7 was set as the threshold for significant differential expression by default. The target coding genes of miRNAs were predicted using psRobot_tar in psRobot (Wu *et al.*, 2012) for plants. GO and KEGG enrichment analyses were conducted using the target genes of differentially expressed miRNAs. Cytoscape software was used to construct the miRNA-gene networks.

#### Data Availability

The raw sequence data of DNA resequencing, BS-seq, smRNA and BioNano molecules reported in this paper have been deposited in the Genome Sequence Archive (GSA, https://ngdc.cncb.ac.cn/gsa/) with accession numbers of CRA007608, CRA007609, CRA007611, CRA007613, respectively. The raw sequence data of lncRNA has been deposited in the Genome Sequence Archive (GSA, https://ngdc.cncb.ac.cn/gsa/) with an accession number of CRA007610 as reported by Mai *et al.* (2022).

**Punta M, Coggill PC, Eberhardt RY, Mistry J, Tate J, Boursnell C, Pang N, Forslund K, Ceric G, Clements J, Heger A, Holm L, Sonnhammer ELL, Eddy SR, Bateman**

**A, Finn RD. 2012.** The Pfam protein families database. *Nucleic Acids Research* **40**: D290–301.

**Shannon P, Markiel A, Ozier O, Baliga NS, Wang JT, Ramage D, *et al.* 2003.** Cytoscape: a software environment for integrated models of biomolecular interaction networks. *Genome Research* **13**: 2498–2504.

**Simão FA, Waterhouse RM, Ioannidis P, Kriventseva EV, Zdobnov EM. 2015.** BUSCO: assessing genome assembly and annotation completeness with single-copy orthologs. *Bioinformatics* **31(19)**: 3210–3212.

**Song QX, Lu X, Li QT, Chen H, Hu XY, Ma B, Zhang WK, Chen SY, Zhang JS. 2013.** Genome-wide analysis of DNA methylation in soybean. *Molecular Plant* **6**, 1961–1974.

**Suo YJ, Sun P, Cheng HH, Han WJ, Diao SF, Li HW, Mai YN, Zhao X, Li FD, Fu JM. 2020.** A high-quality chromosomal genome assembly of *Diospyros oleifera* Cheng. *GigaScience* **9**: 1–10.

**Trapnell C, Pachter L, Salzberg SL. 2009.** TopHat: discovering splice junctions with RNA-Seq. *Bioinformatics* **25**: 1105–1111.

**Trapnell C, Williams BA, Pertea G, Mortazavi A, Kwan G, van Baren M, Salzberg SL, Wold BJ, Pachter L. 2010.** Transcript assembly and quantification by RNA-Seq reveals unannotated transcripts and isoform switching during cell differentiation. *Nature Biotechnology* **28(5)**: 511–515.

**Wang YP, Tang HB, Debarry JD, Tan X, Li JP, Wang XY, Lee T, Jin HZ, Marler B, Guo H, Kissinger JC, Paterson AH. 2012.** MCScanX: a toolkit for detection and evolutionary analysis of gene synteny and collinearity. *Nucleic Acids Research* **40(7)**: e49.

**Wang L, Zhang J, Duan JL, Gao XX, Zhu W, Lu XY, Yang L, Zhang J, Li GQ, Ci WM, et al. 2014.** Programming and inheritance of parental DNA methylomes in mammals. *Cell* **157**: 979–991.

**Wen M, Shen Y, Shi SH, Tang T. 2012.** miREvo: an integrative microRNA evolutionary analysis platform for next-generation sequencing experiments. *BMC Bioinformatics* **13(1)**: 140.

**Wu HJ, Ma YK, Chen T, Wang M, Wang XJ. 2012.** PsRobot: a web-based plant small RNA meta-analysis toolbox. *Nucleic Acids Research* **40**: W22–28.

**Wu H, Xu TL, Feng H, Chen L, Li B, Yao B, Qin ZH, Jin P, Conneely K N. 2015.** Detection of differentially methylated regions from whole-genome bisulfite sequencing data without replicates. *Nucleic Acids Research* **43(21)**: e141.

**Young MD, Wakefield MJ, Smyth GK, Oshlack A. 2010.** Gene ontology analysis for RNA-seq: accounting for selection bias. *Genome Biology* **11(2)**: R14.

**Zhong SL, Fei ZJ, Chen YR, Zheng Y, Huang MY, Vrebalov J, Mcquinn R, Gapper NE, Liu B, Xiang J, *et al*. 2013.** Single-base resolution methylomes of tomato fruit development reveal epigenome modifications associated with ripening. *Nature Biotechnology* **31**: 154–159.

**Zhou L, Chen J, Li Z, Li X, Hu X, Huang Y, et al. 2010.** Integrated profiling of MicroRNAs and mRNAs: MicroRNAs Located on Xq27.3 associate with clear cell renal cell carcinoma. *PLoS One* **5**: e15224.
