## Supplementary Text for "Molecular and genetic mechanisms conferring dissolution of dioecy in *Diospyros oleifera* Cheng"

#### Text S1: Ploidy analysis of the *D. oleifera* collection with diverse sex expression

In Fig. S4, (A)-(E) show the DNA levels in gynoecious, monoecious, androgynomonoecious, andromonoecious, and pseudo-monoecious *D. oleifera* plants, respectively. (F) and (G) show the DNA levels in a diploid *D. lotus* plant and a hexaploid *D. kaki* plant. Peak R10 is DNA. The results indicated that *D. oleifera* trees with diverse types of sex expression were diploid, similar to findings in *D. lotus* (Fig. S4).

#### Text S2: BioNano optical mapping-assisted chromosomal genome assembly of *D. oleifera*

##### S2.1 BioNano optical mapping-assisted chromosomal genome assembly of *D. oleifera*

To assemble the *D. oleifera* genome, a female *D. oleifera* was sequenced, and 104.02 Gb of Illumina paired-end reads were generated. *K*-mer analysis revealed that the genome size and heterozygosity ratio were 670 Mb and 1.26%, respectively (Fig. S5). Subsequently, ~99.76 Gb (~148.9×) of PacBio single-molecule real-time reads (N50 = 14.823kb) and ~177 Gb (~264.2×) of BioNano optical maps (N50 = 14.0 Mb) were generated and integrated into 3790 contigs (~1370 Mb, N50 = 7.55 Mb) and 3353 scaffolds (~1400 Mb, scaffold N50 = 12.64 Mb). After heterozygosity filtration, a *D. oleifera* main genome with 104 contigs (~690 Mb, N50 = 14.94 Mb) and 73 scaffolds (~700 Mb, scaffold N50 = 20.82 Mb) (Table S1) was obtained (Sun and Fu, 2022a). The length of gaps in scaffolds at this stage was 10.01 Mb. Heterozygous sequences with 3735 contigs (~680 Mb, N50 = 1.11 Mb) were also obtained (Table S1; Sun and Fu, 2022b). Synteny analysis between the *D. oleifera* main genome and the heterozygous sequences confirmed that our allele-aware genome was complete; it also revealed several large indels (*e.g.,* on chromosome 7 and chromosome 14) (Fig. S6).

The Hi-C reads (~98.24 Gb, ~146.6×) were used to correct, order, and orient the scaffolds of the main genome into 15 chromosomes (Dudchenko *et al.*, 2017). The percentage of genomic sequences anchoring to Hi-C was 96.07%. Subsequently, the unanchored sequences were aligned to the chromosomes with a coverage ratio of 98.7% and identity ≥ 95%, indicating that most of the unanchored sequences were heterozygous or repetitive sequences. The alignment of the Hi-C and genomic sequences is shown in Fig. S7.

BUSCO analysis (Simão *et al.*, 2015) showed that 86.2% and 92.3% of complete BUSCOs were detected in the main genome and the whole genome (including both the main genome and the heterozygous sequences) (Table S2), respectively. Furthermore, mapping of Illumina short reads (Suo *et al.*, 2020) to the *D. oleifera* main genome yielded proportions of mapped reads and correct pairs of 98.9% and 94.18%, respectively, indicating that the assembled genome had a high level of completeness.

**S2.2 Repetitive sequence and gene annotation**

Repetitive sequences comprised 59.43% of the *D. oleifera* main genome, including 58.11% transposable elements (TEs). The most frequently detected TEs were long terminal repeat retrotransposons (49.11%), followed by DNA TEs (5.27%). Of the long terminal repeats, 29.46% and 17.10% were Ty3/Gypsy and Ty1/Copia, respectively (Table S3; Fig. S8).

The *D. oleifera* main genome contained 22,164 protein-coding genes with a mean transcript size of 6958.94 bp, a mean coding sequence size of 1044.05 bp, and a mean number of exons per gene of 4.66 (Table S4). This number of annotated genes is fewer than in the previous *D. oleifera* genome (Suo *et al.*, 2020) (30,539 genes) or *D. lotus* genome (40,532 genes). Additionally, 23,854 protein-coding genes (91.17% of the total 26,164 genes) had conserved functional motifs or functional terms in the NR, InterPro, and KEGG databases—90.95% (23,796), 76.96% (20,135), and 69.02% (18,058) of the genes, respectively (Table S5).

In terms of non-coding RNAs, 481 miRNAs, 504 tRNAs, 937 rRNAs, and 736 snRNAs were identified, with mean lengths of 115.04, 74.90, 201.50, and 112.99 bp, respectively (Table S6).

**S2.3 Genome synteny**

Good general synteny was observed between the current *D. oleifera* main genome and the genome reported by Suo *et al*. (2020), although chromosome 10 and chromosome 11 in the current main genome contained large reversions compared with chromosome 8 and chromosome 11, respectively, in the previous genome. Additionally, small reversion regions between the current main genome and the previous genome were detected in the end region of chromosome 1 (Fig. S9). Analysis of synteny between the current *D. oleifera* main genome and *D. lotus* genome (Akagi *et al*., 2020) revealed homologous chromosomes between the two genomes (Fig. S10). The male-linked region is located on chromosome 15 in *D. lotus* (Akagi *et al*., 2020), which corresponds to chromosome 4 in the current *D. oleifera* main genome. Compared with the current *D. oleifera* main genome, the *D. lotu*s genome lacked some regions on each chromosome, possibly because of genetic map-based anchoring during *D. lotus* genome sequencing (Akagi *et al*., 2020).

**S2.4 Assembly of the male-unmapped sequence**

Because a gynoecious *D. oleifera* tree was used for the assembly, a male-specific region was not included in the genome. To characterise the sex-linked region in *D. oleifera*, resequencing reads of genetically male individuals were used to reconstruct the sex-linked region. The male-specific reads were assembled into 2860 contigs (43.56 Mb, N50 = 3802) (Table S7), which were regarded as the male-unmapped sequence (Sun and Fu, 2022b). Repetitive sequences comprised 1.45% of the male-specific sequences, including 0.99% TEs (Table S8).

The male-unmapped sequence contained 2952 protein-coding genes with a mean length of 1238.19 bp, and a mean coding sequence of 1108.19 bp (Table S9). Additionally, 2729 protein-coding genes (92.45% of 2952 genes) had conserved functional motifs or functional terms in the NR, InterPro, and KEGG databases—92.38% (2,727), 90.65% (2,676), and 87.53% (2,584) of the genes, respectively (Table S10).

In terms of non-coding RNAs, 4 miRNAs, 147 tRNAs, and 27 snRNAs were identified, with mean lengths of 79.5, 92.32, and 148.15 bp, respectively (Table S11).

**S2.5 Gene family cluster, phylogenetic tree construction, and divergence time estimation**

The main genome combined with the male-unmapped sequence was used for gene family clustering, phylogenetic tree construction, and divergence time estimation. After clustering, 27,638 gene families were detected in *D. oleifera* and 11 other species (Table S12), among which 222 single-copy orthologs were shared by 12 species. Among the five Ericales species (*D. oleifera*, *D. lotus*, *A. chinensis*, *R. delavayi*, and *C. sinensis*), 9550 gene families were shared; 558 gene families consisting of 1026 genes were unique to *D. oleifera* (Fig. S11). The unique genes were significantly enriched in 18 biological process and 12 molecular function GO terms (Fig. S12).

A phylogenetic tree of the 12 plant species was constructed in accordance with the method used by Suo *et al*. (2020). The tree indicated that *D. oleifera* split from *D. lotus* 10.0 million years ago (Fig. S13).

**S2.6 Expansion and contraction of gene families**

We evaluated the expansion and contraction of the gene families in accordance with the method used by Suo *et al*. (2020). Compared with the common ancestor of *D. oleifera* and *D. lotus*, 41 gene families (155 genes) have expanded in *D. oleifera* (Fig. S14); these gene families were significantly enriched in 22 biological process and 8 molecular function GO terms (Fig. S15A). They were also enriched in several KEGG pathways, including ubiquinone and other terpenoid-quinone biosynthesis, protein processing in endoplasmic reticulum, peroxisome, N-Glycan biosynthesis, glyoxylate, and dicarboxylate metabolism (Fig. S16A). In contrast, 593 gene families (1143 genes) have contracted in *D. oleifera*; these gene families were significantly enriched in 14 biological process and 16 molecular function GO terms (Fig. S15B). They were also enriched in several KEGG pathways, including plant-pathogen interaction, biosynthesis of secondary metabolites, and phenylpropanoid biosynthesis (Fig. S16B).

**S2.7 Whole-genome duplication and macrosynteny analysis**

In addition to the ancient whole-genome duplication event that occurred in all dicot species, the $\gamma$ event (all core eudicots share an ancient whole-genome duplication, 4dtv = 0.6), a second whole-genome duplication event occurred in *D. oleifera* and *D. lotus* (4dtv = 0.32/0.35) and may have contributed to the divergence of Ebenaceae from *A. chinensis* and *C. sinensis* (Fig.

S17).

#### Text S3: Genome methylation landscape of *D. oleifera*

To investigate DNA methylation dynamics among floral buds and immature stems of flowering shoots from *D. oleifera* trees of different sexual types, single-base resolution maps of DNA methylation for 30 samples, including floral buds, stems of flowering shoots, and leaves were generated. In total, 2,934,174,248 clean reads were generated, corresponding to 805.14 Gb and > 30-fold coverage of the genome (~690 Mb). To confirm the quality of the sequences, QC20 (>96%) and QC30 (86.22-93.27%) values, GC contents (22.14-27.02%, average of 23.20%) (Table S16), and bisulphite conversion rates of total C (99.71%) (Table S17) were calculated. An average of 63.02% (57.67-66.44%) of clean reads were mapped to the reference genome, with a mean duplication rate of 21.52% (13.68-36.56%) (Table S18). After mapping, the mean coverage depth of the genome was 18.14% (Table S19). For instance, for the female floral buds of the monoecious #108 tree (M108_F), the coverage depths to the main genome, male-unmapped sequence, and chloroplast genome were 17.41%, 0.006%, and 363.15%, respectively (Fig. S21A). In stems of flowering shoots of the monoecious #108 tree (M108_S), the coverage depths were similar (Fig. S21B). Moreover, the coverage depth for the main genome was similar in other samples.

In M108_F, 87.28% of the main genome was covered, compared with 0.23% and 34.95% of the male-unmapped sequence and the chloroplast genome, respectively (Fig. S21A). In M108_S, 87.69% of the main genome was covered, compared with 0.36% and 32.63% of the male-unmapped sequence and the chloroplast genome, respectively (Fig. S21B). The genome coverage rates in other samples showed similar trends.

For all samples, 2.24%, 2.85%, 3.74%, and 1.88% of the total, CG, CHG, and CHH cytosines were methylated, respectively (Table S20); the global methylation level was 24.90% (Table S21). The mean CG, CHG, and CHH methylation levels were 76.39%, 56.11%, and 8.11%, respectively (Table S21). In M108_F, 10.97% mCG, 15.75% mCHG, and 73.72%

mCHH constituted the total mC (Fig. S22A). In M108_S, 10.20% mCG, 14.67% mCHG, and 75.13% mCHH constituted the total mC (Fig. S22B). The proportions of mCHH to total mC were > 50% in all samples.

Hierarchical clustering based on PCA of the methylation levels of CG (Fig. S23A) and CHG (Fig. S23B) subcontexts demonstrated that tissues (*i.e.,* floral buds, stems, and leaf) from the same tree had a close relationship, whereas clustering based on PCA of the methylation level of CHH showed that similar tissues had a close relationship among multiple trees (Fig. S23C).

### Text S4: Whole transcriptome analysis

#### S4.1 Identification of mRNAs, lncRNAs, and transcripts of uncertain coding potential (TUCPs)

To investigate sex-biased mRNAs, lncRNAs, and TUCPs in *D. oleifera*, we used the Illumina HiSeq2500 platform to perform RNA sequencing of the floral buds and stems of flowering shoots obtained from single- and co-sex *D. oleifera* trees. In total, we obtained 2331.62 million raw reads from 26 samples with mean Q20, Q30, and GC contents of 98.00%, 94.22%, and 47.20%, respectively. After the removal of adapter sequences and low-quality reads, 2.29 billion clean reads were generated. Subsequently, 71.21% and 62.82% of the clean reads were totally and uniquely mapped to the combined genome, respectively. The mean percentages of non-splice and splice reads were 44.66% and 18.16%, respectively (Table S24).

A similar distribution of reads on chromosomes between M108_F (Fig. S30A) and A13_S (Fig. S30B) was observed. Additionally, 82.3% and 78.2% of mapped reads were classified as mRNAs in M108_F and A13_S, respectively (Fig. S31A and B). Similar results were obtained for other samples.

In this study, 7880 lncRNAs were obtained (Fig. S32A). The lincRNAs, antisense_lncRNAs, and intronic_lncRNAs constituted 87.3%, 12.7%, and 0% of the total lncRNAs, respectively (Fig. S32B). The number of mRNA exons was larger than the number of lncRNA exons (Fig. S32C), and the transcript length of mRNAs was longer than the transcript length of lncRNAs

(Fig. S32D). The open reading frames of mRNAs were typically longer than the open reading frames of lncRNAs (Fig. S33E).

In this study, 2840 TUCPs were identified. The exon number, total length, and open reading frame length were similar for mRNAs and TUCPs (Fig. S32F-H). The FPKM values of mRNA were twofold greater than the FPKM values of lncRNA or TUCP (Fig. S33A). The square of the Pearson correlation coefficient between samples in the same group was > 0.8, indicating reliable biological replicates (Fig. S33B).

**S4.2 Clustering based on differentially expressed mRNAs, lncRNAs, and TUCPs**

Clustering of samples based on the FPKM values of all differentially expressed mRNAs (DEGs) indicated that the same tissue (*e.g.,* female floral buds or immature stems obtained from trees with the same sexual type) exhibited a close relationship (Fig. S34A). Clustering based on the FPKM values of all differentially expressed lncRNAs (DELs) and TUCPs (DETs) indicated that different tissues obtained from the same tree exhibited a close relationship (Fig. S34B and C).

**S4.3 Sex-biased expression of mRNA**

We identified 1840 DEGs between M_F and M_M. Compared with M_M, 1015 genes were upregulated and 825 genes were downregulated in M_F (Fig. S35A). The chromosomal distribution of DEGs is shown in Fig. S35C. Furthermore, 3523 DEGs were identified between G_F and A_M. Compared with A_M, 1788 genes were upregulated and 1735 genes were downregulated in G_F (Fig. S35B). The chromosomal distribution of DEGs is shown in Fig. S35D.

A GO category (copper ion binding [GO:0005507]) enriched in upregulated DEGs (Fig. S36A and B) and 18 GO categories enriched in downregulated DEGs (*e.g.,* transporter activity [GO:0005215], membrane [GO:0016020], transmembrane transporter activity [GO:0022857]) (Fig. S36C and D) were shared between monoecious plants and single-sex plants. These findings imply that a similar system regulates floral sex-type expression in monoecious and single-sex *D. oleifera*. Twelve GO categories were enriched in upregulated DEGs in monoecious plants, but not in single-sex plants (*e.g.,* terpene synthase activity [GO:0010333], carbon-oxygen lyase activity, acting on phosphates [GO:0016838]) (Fig. S36A). Five GO categories were enriched in downregulated DEGs in monoecious plants, but not in single-sex plants (hydrolase activity [GO:0016787], inorganic anion exchanger activity [GO:0005452], anion:anion antiporter activity [GO:0015301], anion transport [GO:0006820], and enzyme inhibitor activity [GO:0004857]) (Fig. S36C). These results imply that specific mechanisms regulate sex differentiation in monoecious *D. oleifera*.

In the andromonoecious plants, 4407 DEGs were identified between AM_M and AM_H. Compared with AM_H, 2230 and 2177 genes were up- and downregulated, respectively in AM_M. Upregulated DEGs were enriched in 96 GO categories, including membrane (GO:0016020), transporter activity (GO:0005215), pectinesterase activity (GO:0030599), carbohydrate metabolic process (GO:0005975), and catalytic activity (GO:0003824) (Fig. S37A). Downregulated DEGs were enriched in 112 GO categories, including regulation of transcription, DNA-dependent (GO:0006355), regulation of RNA metabolic process (GO:0051252), regulation of RNA biosynthetic process (GO:2001141), and regulation of macromolecule biosynthetic process (GO:0010556) (Fig. S37B).

##### S4.4 Sex-biased expression of lncRNAs

In total, 266 DELs were identified between M_F and M_M. Compared with M_M, 113 and 153 lncRNAs were up- and downregulated in M_F, respectively (**Fig. S38A**). The chromosomal distribution of DELs is shown in **Fig. S38C**. Additionally, 593 DELs were identified between G_F and A_M. Compared with A_M, 253 and 340 lncRNAs were up- and downregulated in G_F, respectively (**Fig. S38B**). The chromosomal distribution of DELs is shown in **Fig. S38D**.

Non-shared GO categories significantly enriched in mRNAs co-located with upregulated lncRNAs in the monoecious comparison (M_F compared with M_M), rather than the single-sex comparison (G_F compared with A_M), included cytochrome-c oxidase activity (GO:0004129), haem-copper terminal oxidase activity (GO:0015002), oxidoreductase activity, acting on a haem group of donors (GO:0016675), and oxidoreductase activity, acting on a haem group of donors, oxygen as acceptor (GO:0016676) (**Fig. S39A**).

mRNAs co-located with downregulated lncRNAs in the monoecious comparison were enriched in one GO category (DNA catabolic process [GO:0006308]) (**Fig. S39B**), compared with the 19 GO categories in the single-sex comparison (**Fig. S39C**).

We identified 35 common GO categories significantly enriched in mRNAs coexpressed with downregulated lncRNAs in the monoecious and single-sex comparisons, including enzyme regulator activity (GO:0030234), and pectinesterase activity (GO:0030599). Furthermore, we identified 15 non-shared GO categories significantly enriched in mRNAs coexpressed with downregulated lncRNAs in the monoecious comparison, rather than the single-sex comparison (*e.g.,* hydrolase activity, acting on ester bonds [GO:0016788], and single-organism process [GO:0044699]) (**Fig. S40A and B**).

In andromonoecious plants, 428 DELs were identified in the comparison of AM_M and AM_H. Compared with AM_H, 249 and 179 lncRNAs were up- and downregulated in AM_M, respectively. mRNAs coexpressed with upregulated lncRNAs were enriched in 81 GO categories, including pectinesterase activity (GO:0030599) and enzyme regulator activity (GO:0030234) (**Fig. S41A).** mRNAs coexpressed with downregulated lncRNAs were enriched in 10 GO categories, including oxidoreductase activity (GO:0016491) and haem binding (GO:0020037) (**Fig. S41B)**.

##### S4.5 Sex-biased expression of TUCPs

In total, 89 DETs were identified between M_F and M_M. Compared with M_M, 56 and 33 TUCPs were up- and downregulated in M_F, respectively (Fig. S42A). The chromosomal distribution of DETs is shown in Fig. S42C. Additionally, 163 DETs were identified between G_F and A_M. Compared with A_M, 73 and 90 TUCPs were up- and downregulated, respectively (Fig. S42B). The chromosomal distribution of TUCPs is shown in Fig. S42D.

We identified 61 common GO categories enriched in mRNAs coexpressed with downregulated TUCPs in the monoecious and single-sex comparisons, including pectinesterase activity (GO:0030599) and enzyme regulator activity (GO:0030234). Moreover, we identified 16 non-shared GO categories enriched in mRNAs coexpressed with downregulated TUCPs in the monoecious comparison, rather than the single-sex comparison (*e.g.,* phosphorylation [GO:0016310] and single-organism cellular process [GO:0044763]) (Fig. S43A and B).

In the andromonoecious *D. oleifera*, 181 DETs were identified in the comparison of AM_M and AM_H. Compared with AM_H, 91 and 90 TUCPs were up- and downregulated, respectively. mRNAs coexpressed with upregulated TUCPs were enriched in 125 GO categories, including enzyme regulator activity (GO:0030234), pectinesterase activity (GO:0030599), and small GTPase regulator activity (GO:0005083) (Fig. S44A). mRNAs coexpressed with downregulated TUCPs were enriched in 14 GO categories, including iron ion binding (GO:0005506), oxidation-reduction process (GO:0055114), and oxidoreductase activity (GO:0016491) (Fig. S44B).

The KEGG pathways with significant enrichment in mRNAs coexpressed with downregulated lncRNAs and TUCPs between female (or hermaphroditic) and male floral buds in both single- and co-sex systems included plant-pathogen interaction, pentose and glucuronate interconversions, starch and sucrose metabolism, and galactose metabolism (Fig. S45).

**S4.6 Identification of miRNAs**

In this study, 498.93 million 50 bp single-end reads were generated; after filtering, 324.29 million clean reads with a length of 18-30 nt were retained for analysis (Tables S25 and S26). The miRNA length distributions were similar among libraries, and 24-nt-long RNAs were most abundant (Fig. S46A). After length filtration, 88.42% of the clean reads (mean value) were mapped to the combined genome (Table S27). In the floral buds of M108, reads were equally distributed on the 15 chromosomes of the main genome, whereas few reads were aligned to the male-unmapped sequences (Fig. S46B). The distribution patterns were similar among samples. Repeat classification analysis indicated that most reads accumulated on reverse long terminal repeats and forward long terminal repeats (Fig. S46C). Annotation of unique small RNA reads in M108_F is shown in Fig. S46D. The results indicated that 402 conserved miRNAs in 70 miRNA families and 89 predicted novel miRNAs were present among the 26 small RNA libraries. The $\log_{10}$[transcripts per million + 1] values of most miRNAs were < 2 (Fig. S47A). Clustering of samples based on the transcripts per million values of all differentially expressed miRNAs (DEMs) indicated that the same tissues (*e.g.,* female floral buds or immature floral stems) obtained from trees of the same sexual type (*e.g.,* gynoecy) have a close relationship (Fig. S47B).

**S4.7 Sex-biased expression of miRNAs**

In total, 26 DEMs were identified in the monoecious comparison (M_F compared with M_M). Compared with M_M, 11 and 15 DEMs were up- and downregulated in M_F, respectively. The up- and downregulated DEMs included one and four novel miRNAs, respectively. Additionally, 70 DEMs were identified in the single-sex comparison (G_F compared with A_M). Compared with A_M, 26 and 44 DEMs were up- and downregulated in G_F. The up- and downregulated DEMs included seven and eight novel miRNAs, respectively.

mRNAs targeted by DEMs were significantly enriched in 11 GO categories in the single-sex comparison, including protein binding (GO:0005515), nucleus (GO:0005634), and response to endogenous stimulus (GO:0009719) (Fig. S48A).

In andromonoecious plants, 75 DEMs were identified in the comparison of AM_M and AM_H. Compared with AM_H, 40 and 35 DEMs were up- and downregulated in AM_M, respectively. The up- and downregulated DEMs included 10 and 9 novel miRNAs, respectively. mRNAs targeted by DEMs were significantly enriched in 13 GO categories, including nucleus (GO:0005634), protein binding (GO:0005515), and membrane-bounded organelle (GO:0043227) (Fig. S48B).

**S4.8 Identification of circRNAs**

In total, 12,893 novel circRNAs were identified in the 26 samples. Most circRNAs were generated from exon and intron sequences (Fig. S49A), and the length of most circRNAs was ≤ 3000 nt (Fig. S49B).

**S4.9 ceRNA networks**

If an RNA molecule class possesses at least one common miRNA response element that is accessible to miRNA binding, those molecules are regarded as competing endogenous RNAs (ceRNAs) (Salmena *et al*., 2011). mRNAs and ncRNAs, such as lncRNAs and circRNAs, communicate and co-regulate by competitively binding the same miRNAs; these are considered ceRNAs (Tay *et al*., 2014). We performed circRNA, miRNA, and mRNA association analyses to construct ceRNA networks.

No differentially expressed circRNAs were identified in M_F compared with M_M. In the single-sex comparison (G_F and A_M), a ceRNA network was constructed based on 6 downregulated circRNAs, 12 upregulated miRNAs, and their 11 downregulated targets (Fig. S50A). A second network was constructed based on 6 upregulated circRNAs, 30 downregulated miRNAs, and their 46 upregulated targets (Fig. S50B).

In andromonoecious plants, a ceRNA network was constructed based on 13 downregulated circRNAs, 20 upregulated miRNAs, and their 50 downregulated targets in AM_M compared with AM_H (Fig. S50C). A second network was constructed based on 19 upregulated circRNAs, 25 downregulated miRNAs, and their 49 upregulated targets in AM_M compared with AM_H (Fig. S50D).

ceRNA networks were constructed based on integrated analysis of lncRNAs, miRNAs, and mRNAs. Four upregulated DELs (LNC_006965, LNC_006631, LNC_003380, and LNC_003283), 3 downregulated DEMs (zma-miR396g-3p, hbr-miR156, and ata-miR395b-3p), and their 21 upregulated DEGs targets in the single-sex comparison (G_F compared with A_M) were used to construct a ceRNA network (Fig. S51A). In andromonoecious plants, a ceRNA network was constructed based on one upregulated DEL (LNC_002635), five downregulated DEMs (ath-miR319c, gma-miR319q, pta-miR319, ppt-miR319a, and ptc-miR319e) and their nine upregulated DEG targets in AM_M compared with AM_H (Fig. S51B).

##### S4.10 Core gene networks correlated with sex differentiation in female and male floral buds

To evaluate the regulatory paths of sex differentiation in female and male floral buds in co- and single-sex systems, coexpression patterns were visualised by weighted correlation network analysis using all female and male floral buds that had been subjected to RNA-Seq analysis

(Table S30). A scale-free topology model fit soft threshold of 7 (Fig. S52A and B) was applied and the coexpression pattern was clustered into 21 modules (Fig. S52C and D).

The midnight-blue and yellow modules showed the strongest positive correlations with sex dimorphism, and they may promote the development of male floral buds (Fig. S52E). The midnight-blue module contained 156 genes, which were significantly (corrected $P < 0.05$) enriched in five GO terms: regulation of transcription DNA-templated, regulation of primary metabolic process, regulation of gene expression, regulation of cellular macromolecule biosynthetic process, and copper ion binding (Fig. S53A). The yellow module contained 378 genes, which were significantly enriched in the organic substance transport GO term (Fig. S53B).

The pink and green modules showed the strongest negative correlations with sex dimorphism, and they may promote the development of female floral buds. The pink module contained 285 genes, which were significantly enriched in five GO terms: UDP-glycosyltransferase activity, transferase activity, transferring glycosyl groups, single-organism process, single-organism metabolic process, and oxidoreductase activity (Fig. S53C). Genes in the pink module were also significantly enriched in two KEGG pathways: phenylpropanoid biosynthesis and linoleic acid metabolism (Fig. S53D).

Genes with respective connection weights of > 0.2, 0.25, 0.2, and 0.23 in the midnight-blue, yellow, pink, and green modules were used to construct core gene networks (Fig. S54A-D).

#### Text S5: Key candidate genes in the sex-linked region of chromosome 4 complement functional sex dimorphism

Two genes located in the sex-linked region of chromosome 4 (Fig. 8A; Fig. S58) were notable. The first gene was a MADS-box transcription factor *GLO* (evm.model.Chr4.1456), which was downregulated in female floral buds compared with male floral buds in single- and co-sex plants. *GLO* was upregulated in male floral buds (AM_M) compared with hermaphroditic floral buds (AM_H) in andromonoecious plants (Fig. 8D), implying a function in male promotion. The second gene was a two-component response regulator *ARR9* (evm.model.Chr4.1534), which was upregulated in female tissues compared with male tissues in single- and co-sex plants (Fig. 8D), implying a function in female promotion.

Several other genes, including callose synthase 5 (*CALS5*; evm.model.Chr4.1596 and evm.model.Chr4.1597) and aspartic proteinase nepenthesin-1 (*nep1*; evm.model.Chr4.1641), were differentially expressed between male and female tissues in the single- and co-sex types; they were also functionally associated with sexual expression (Fig. 8D; Table S42). Collectively, the results imply the presence of genes that function in sexual expression in the sex-linked region, which may contribute to the differentiation of the X and Y chromosomes.
