## Supplementary Figures for "Molecular and genetic mechanisms conferring dissolution of dioecy in *Diospyros oleifera* Cheng"

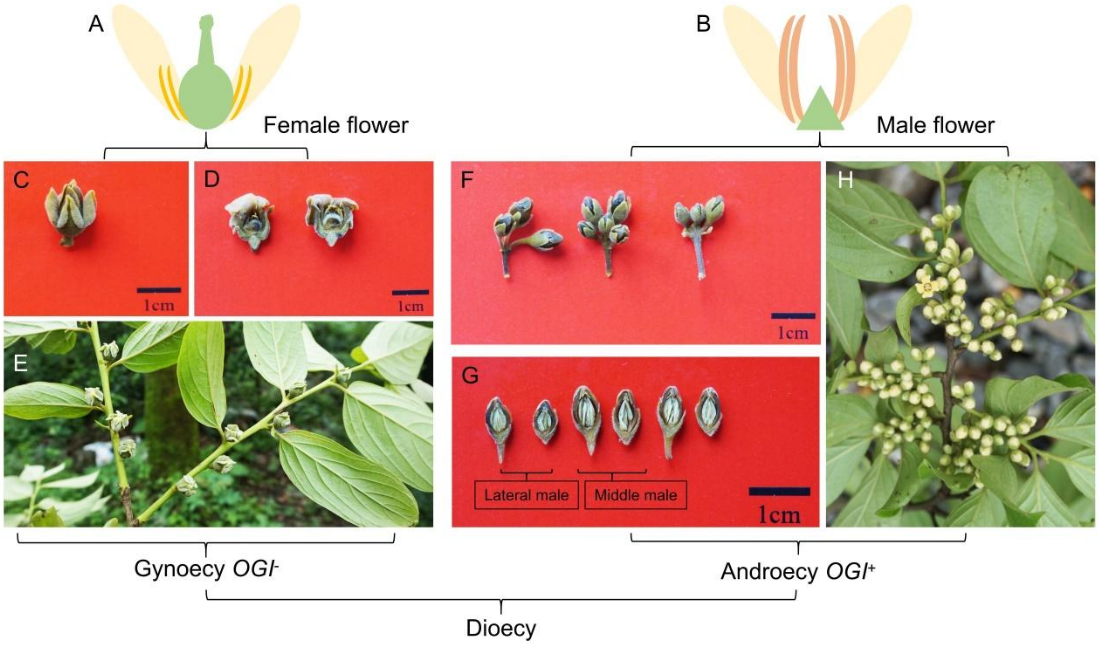


**Fig. S1 Floral buds of dioecious *D. oleifera*.** Illustrations of **(A)** female and **(B)** male *D. oleifera* flowers. **(C)** An intact female floral bud. **(D)** Dissection of a female floral bud showing active pistil and arrested stamen. **(E)** Female shoots bearing solitary female floral buds. **(F)** Intact male floral buds. **(G)** Dissection of male floral buds showing arrested pistils and active stamens. **(H)** Male shoots bearing male floral buds which formed three (or more)-flower cymes.

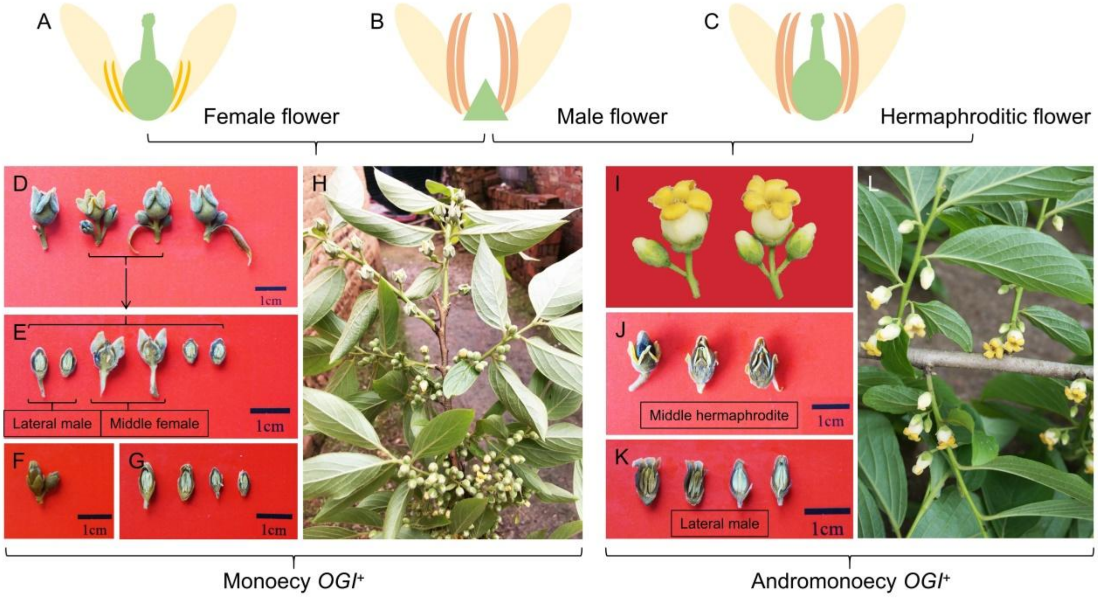


**Fig. S2 Floral buds of monoecious and andromonoecious *D. oleifera*.** Illustrations of **(A)** female, **(B)** male, and **(C)** hermaphroditic *D. oleifera* flowers. **(D)** Female floral bud with two small male floral buds in lateral obtained from a monoecious *D. oleifera.* **(E)** Dissection of a middle female floral bud and two lateral male floral buds. **(F)** Male floral buds which formed a three-flower cyme in monoecious *D. oleifera*. **(G)** dissection of male floral buds; **(H)** Above female shoots and lower male shoots of a monoecious *D. oleifera*. **(I)** Middle hermaphroditic and lateral male floral buds of an andromonoecious *D. oleifera*. **(J)** Dissection of middle hermaphroditic floral buds showing active pistils and stamens. **(K)** Dissection of lateral male floral buds showing arrested pistils and active stamens. **(L)** Flowering shoots bearing hermaphroditic and male floral buds of an andromonoecious *D. oleifera.*

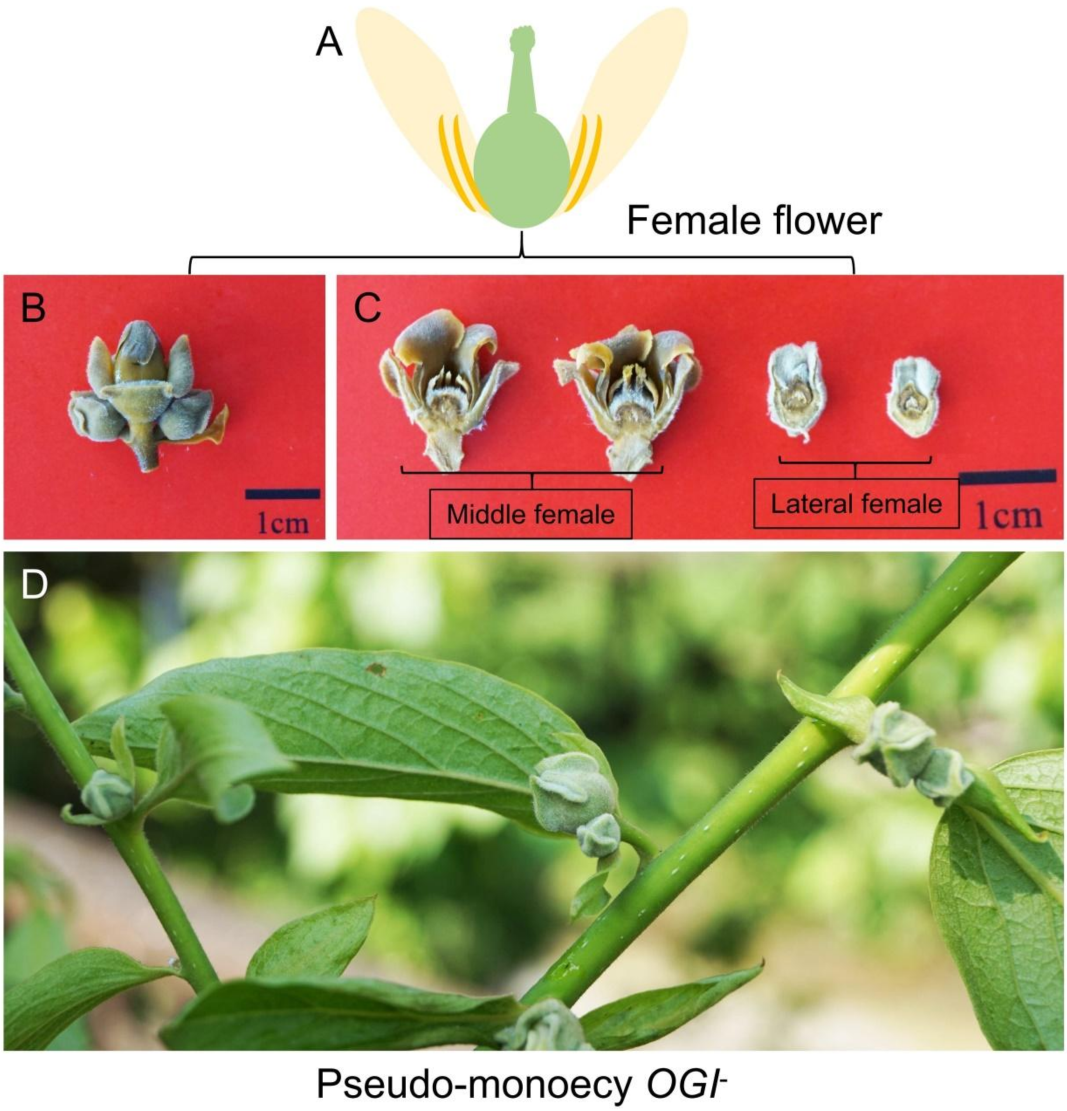


**Fig. S3 Floral buds of pseudo-monoecious *D. oleifera*. (A)** An illustration of a female *D. oleifera* flower. **(B)** Female floral buds which formed a three-flower cyme. **(C)** Dissection of a middle and a lateral female floral bud. **(D)** Flowering shoots bearing female floral buds which formed two or three-flower cymes.

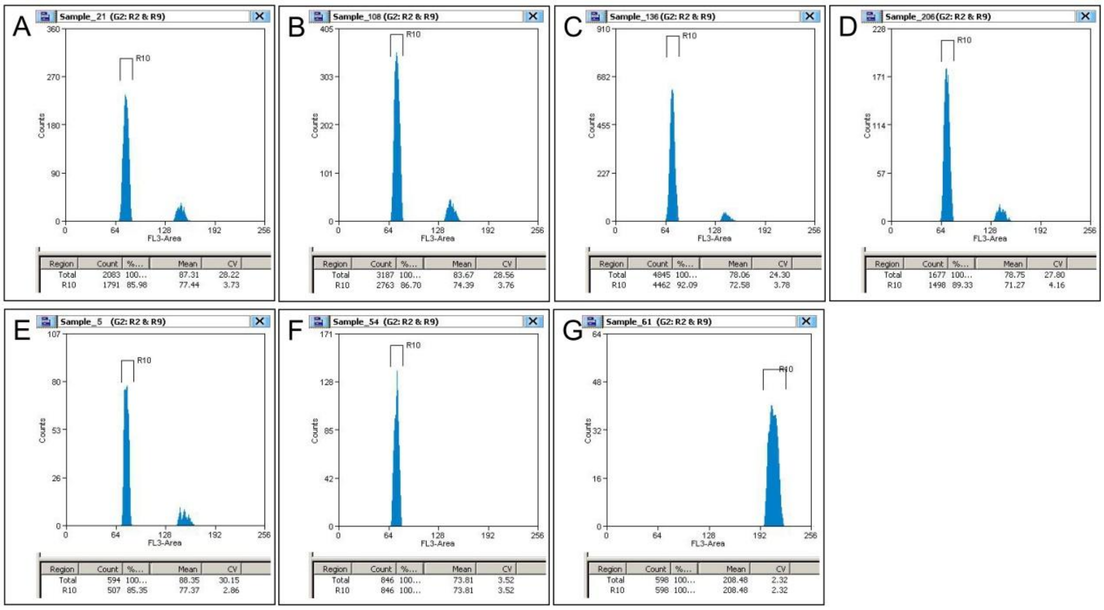


**Fig. S4 Flow-cytometry results of *D. oleifera*, *D. lotus,* and *D. kaki*. (A) - (E)** represent DNA levels of gynoecious, monoecious, androgynomonoecious, andromonoecious, and pseudo-monoecious *D. oleifera* plants, respectively. **(F)** and **(G)** represent DNA levels of a diploid *D. lotus* plant and a hexaploid *D. kaki* plant, respectively. Peak R10 is DNA, and small duplicated DNA peaks were observed in **(A) - (E)**.

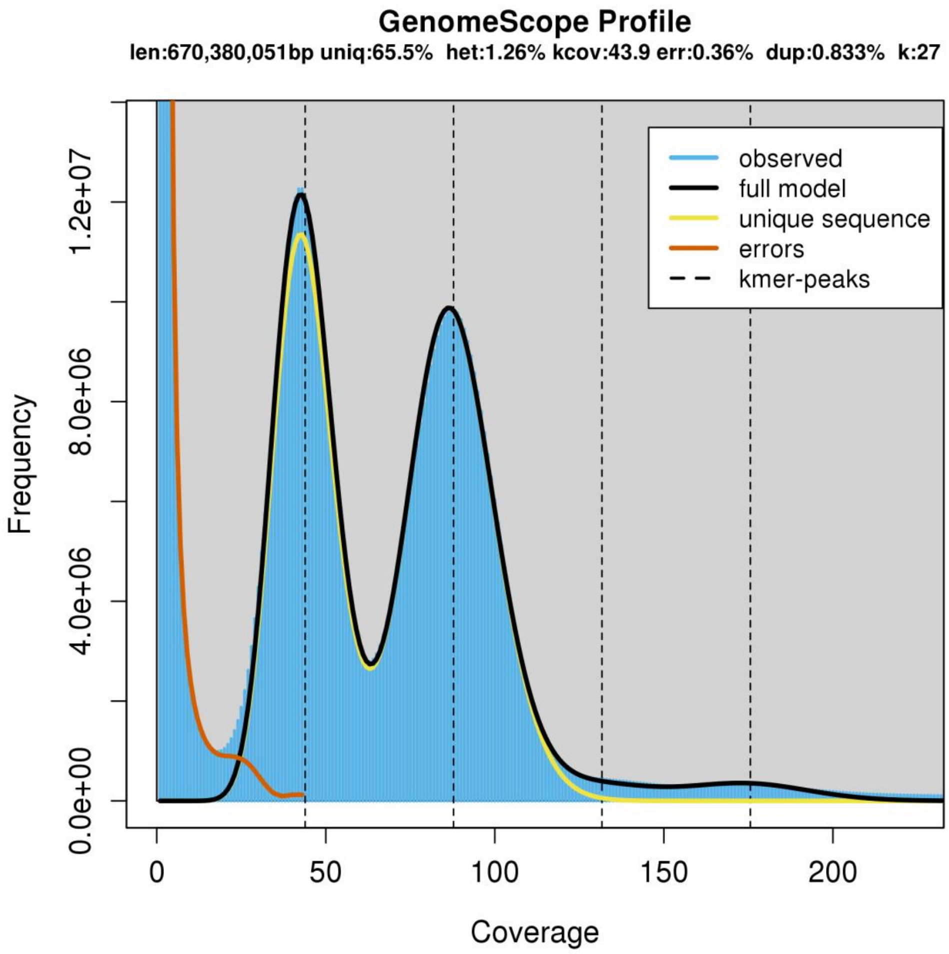


**Fig. S5 GenomeScope (http://qb.cshl.edu/genomescope/) profile of *D. oleifera* genome based on the Illumina genomic sequence data.**

40

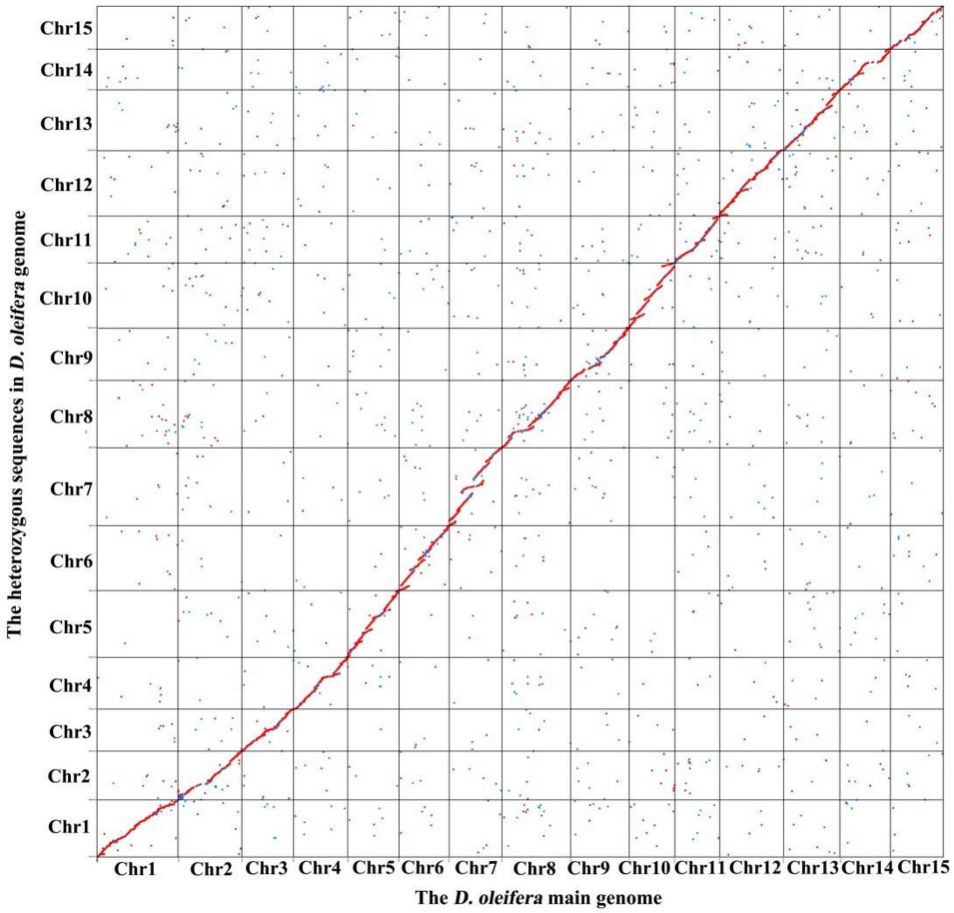


41 **Fig. S6 Synteny analysis between the *D. oleifera* main genome and the heterozygous**
42 **sequences.**

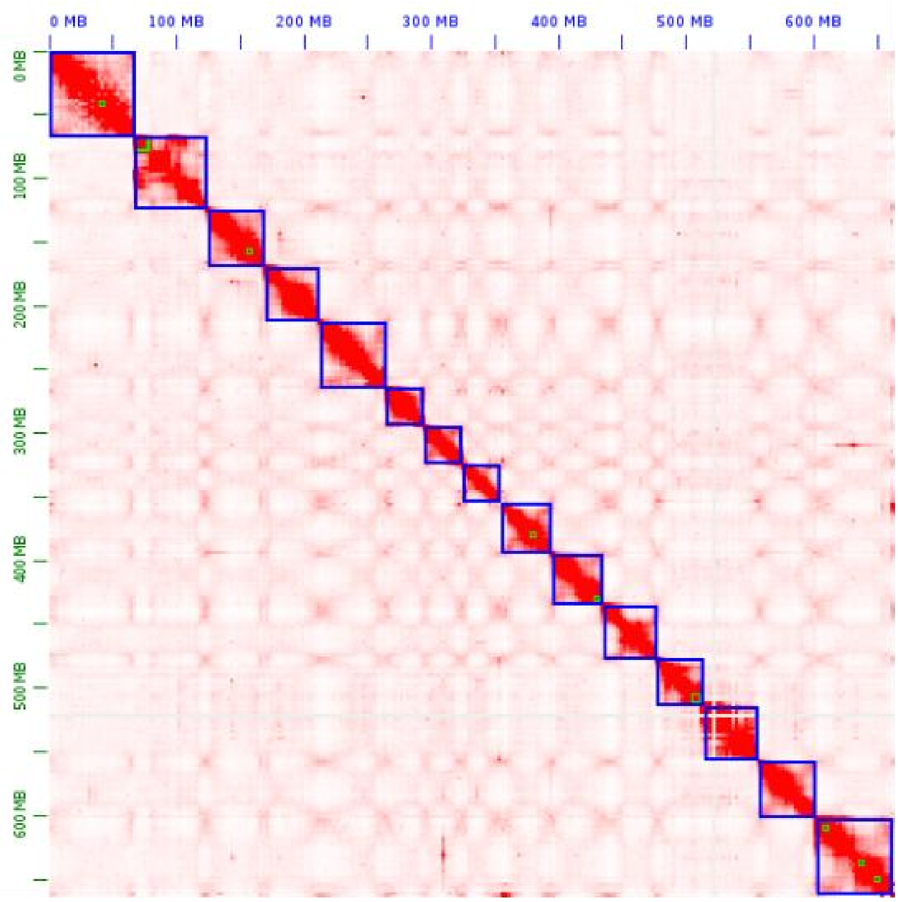


**Fig. S7 Hi-C interaction heat map for *D. oleifera* main genome showing interactions among 15 chromosomes.**

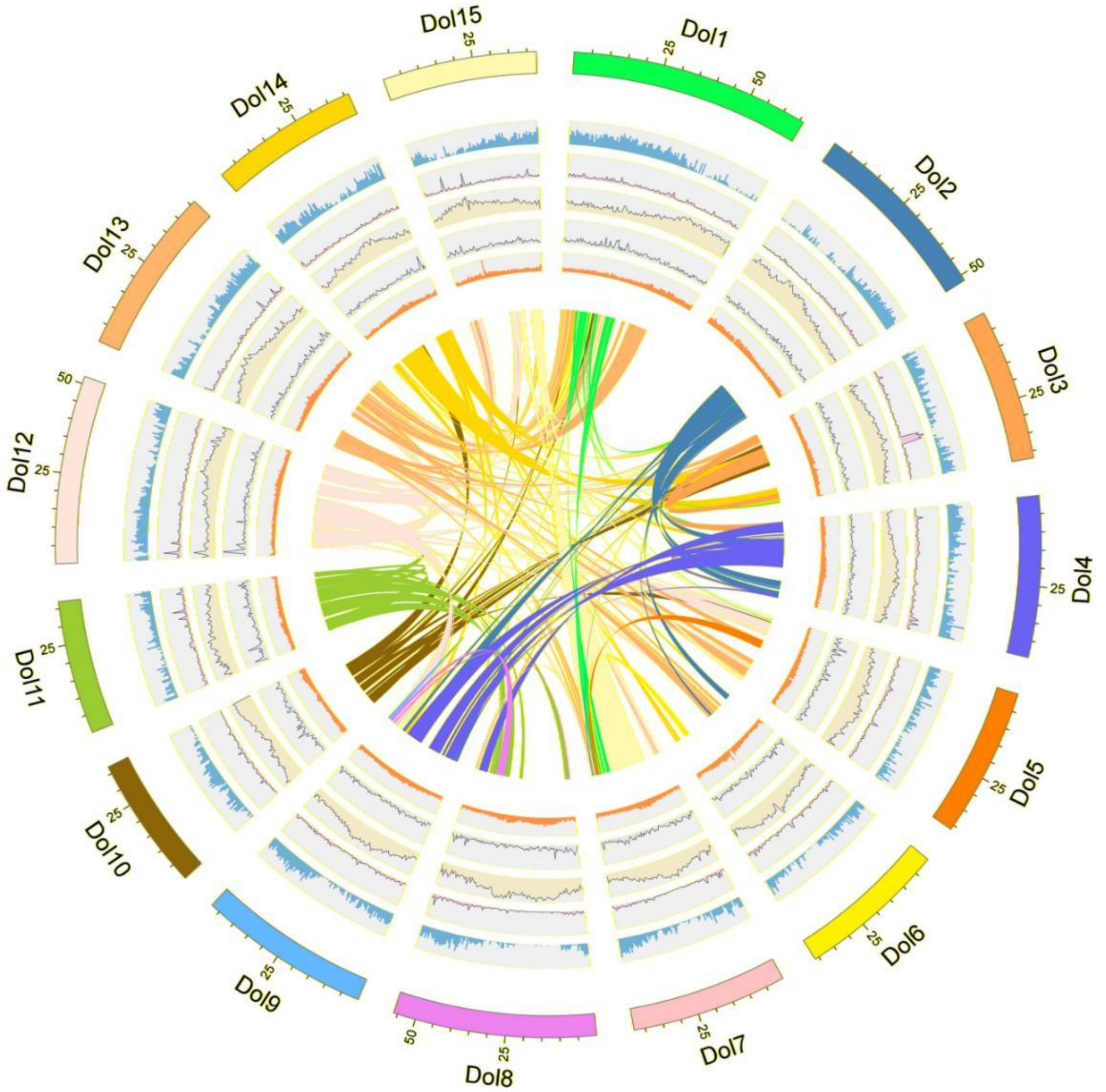


**Fig. S8 *D. oleifera* genome features. Tracks from outside to inside are as follows: the distribution of gene density, LINE retrotransposons density, LTR retrotransposons density, DNA transposons density, GC density, and syntenic blocks.**

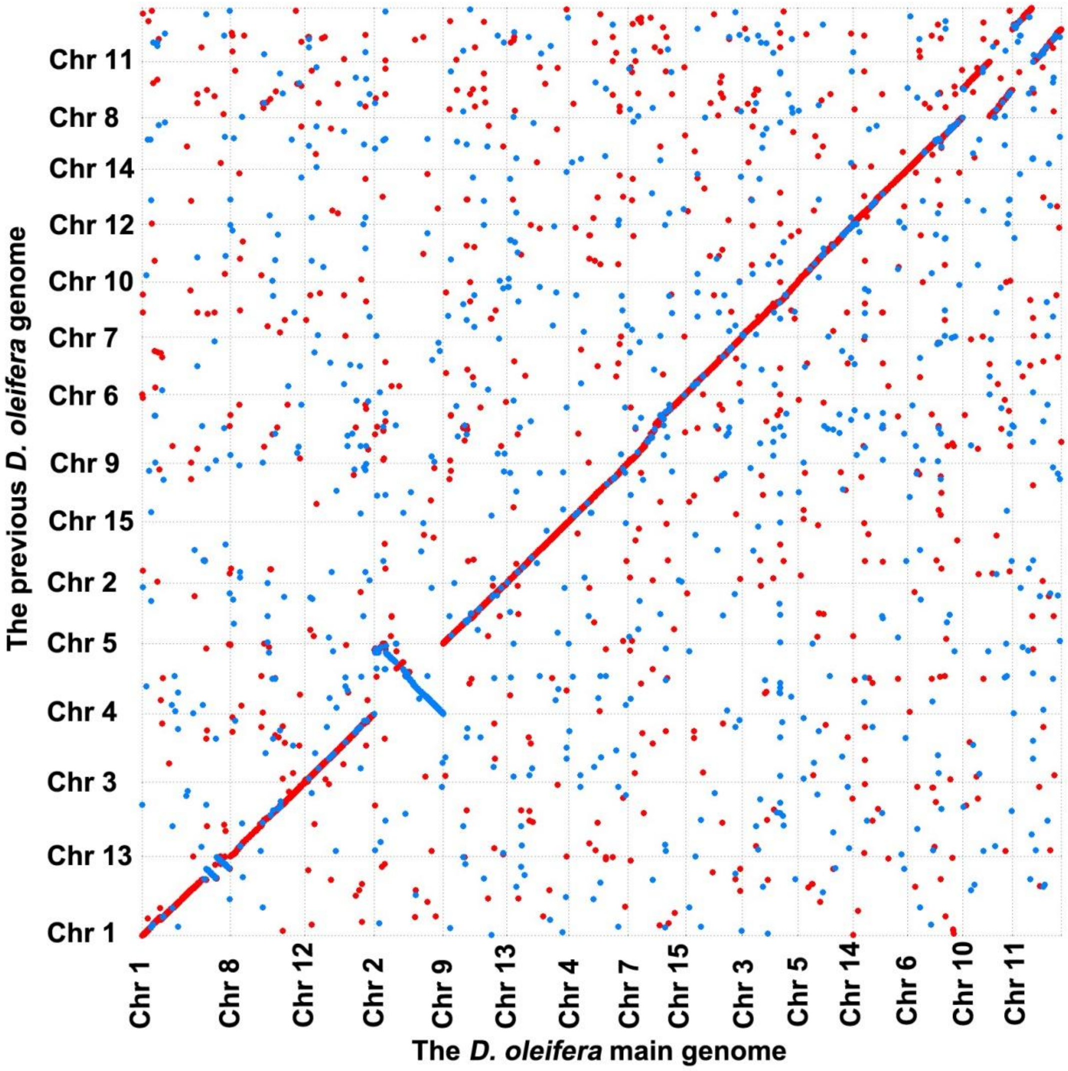


**Fig. S9 Synteny analysis between the current *D. oleifera* main genome and the previous *D. oleifera* genome.**

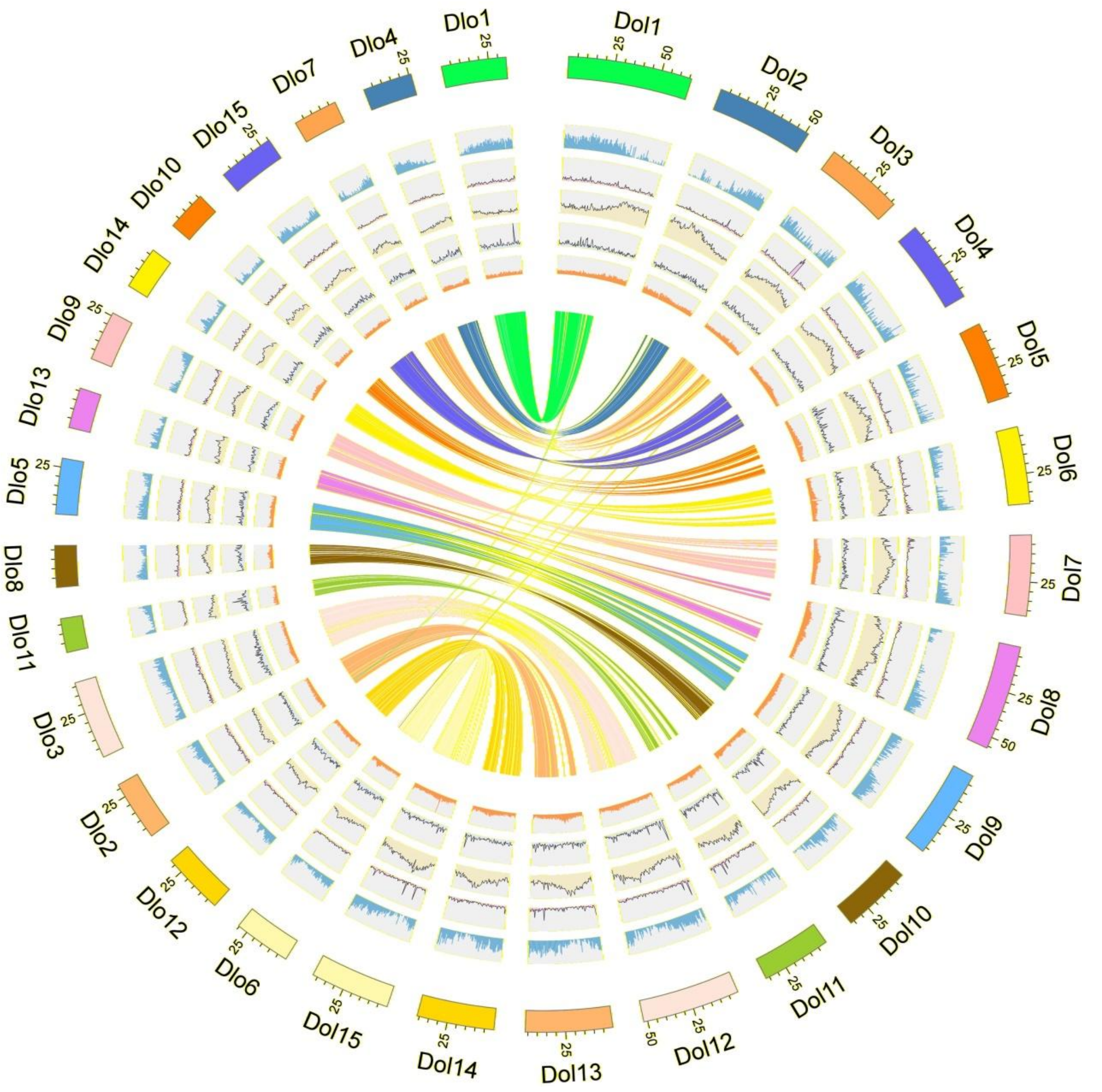


**Fig. S10 Synteny analysis between the current *D. oleifera* main genome and the *D. lotus* genome.** Tracks from outside to inside are as follows: the distribution of gene density, LINE retrotransposons density, LTR retrotransposons density, DNA transposons density, GC density, and syntenic blocks. Dol and Dlo represent *D. oleifera* and *D. lotus*, respectively.

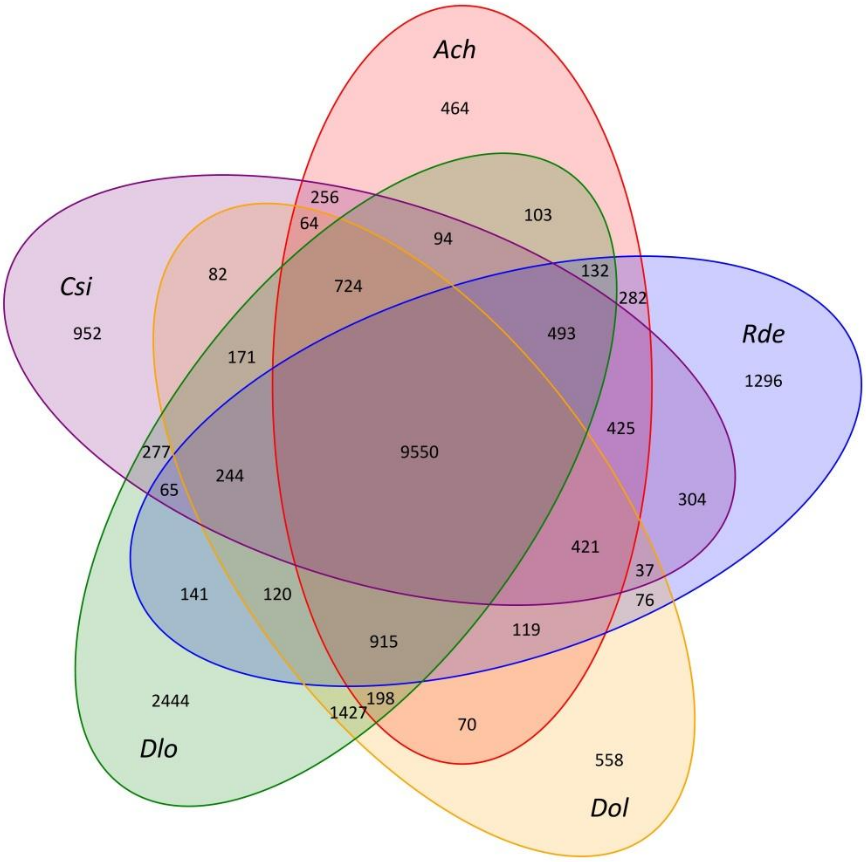


**Fig. S11 Common and unique gene families among 5 species in the Ericales.** *Ach*, *Rde*, *Dol*, *Dlo*, *Csi* represent *Actinidia chinensis*, *Rhododendron delavayi*, *Diospyros oleifera*, *Diospyros lotus*, *Camellia sinensis*, respectively.

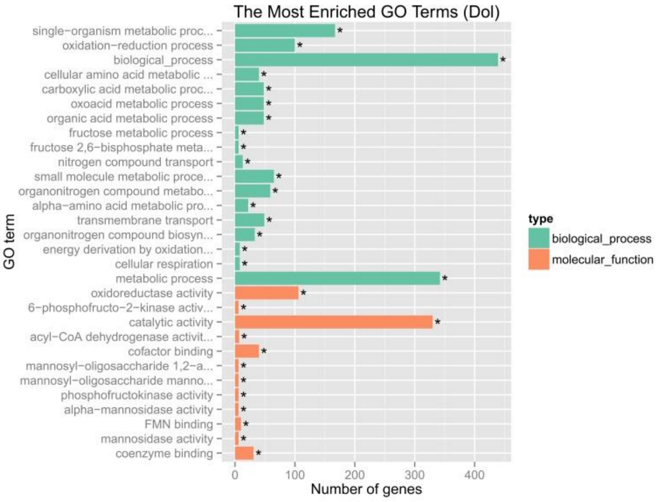


**Fig. S12 GO enrichments of genes unique to *D. oleifera***

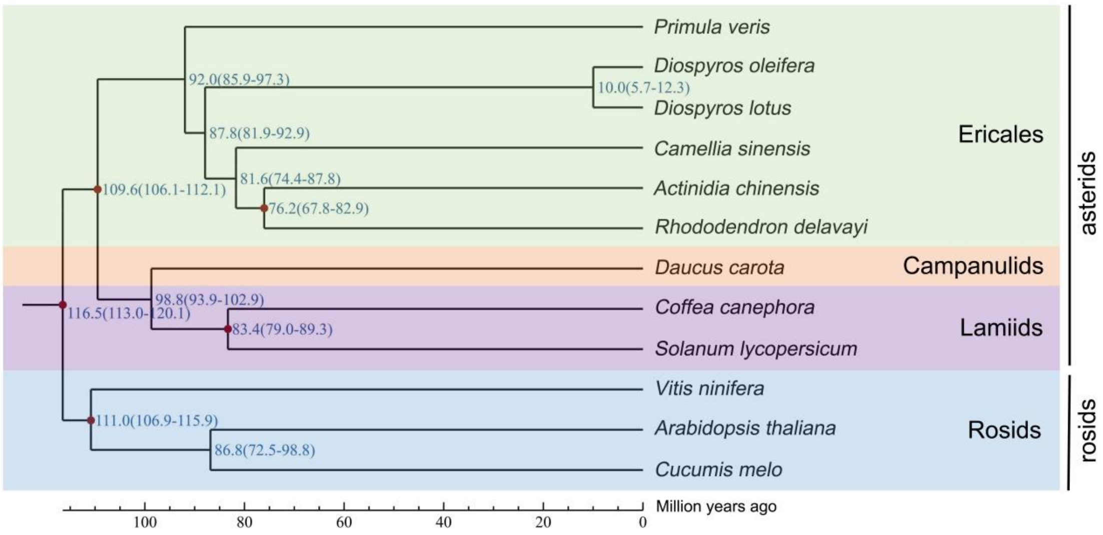


**Fig. S13 Estimation of divergence time.** The numbers on the nodes represent the divergence times from the present (million years ago, Mya).

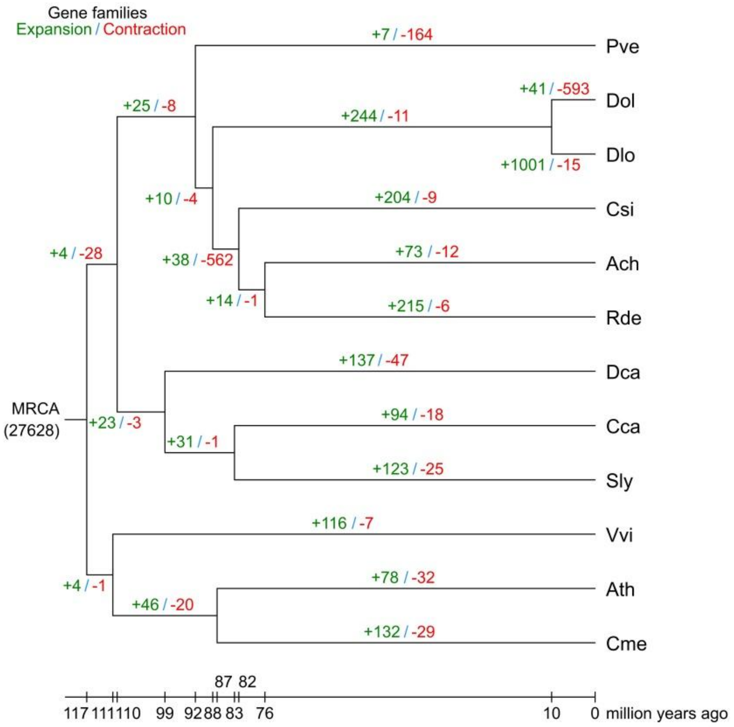


**Fig. S14 Expansion and contraction in gene families.** Pve, Dol, Dlo, Csi, Ach, Rde, Dca, Cca, Sly, Vvi, Ath and Cme represent *Primula veris*, *Diospyros oleifera*, *Diospyros lotus*, *Camellia sinensis*, *Actinidia chinensis*, *Rhododendron delavayi*, *Daucus carota*, *Coffea canephora*, *Solanum lycopersicum*, *Vitis vinifera*, *Arabidopsis thaliana* and *Cucumis melo*, respectively.

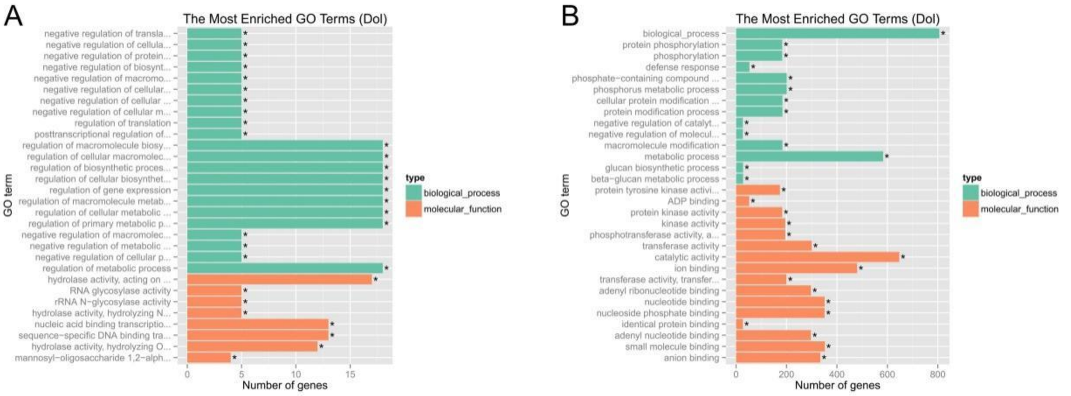


**Fig. S15 GO enrichments of (A) expansion and (B) contraction genes in *D. oleifeara*.**

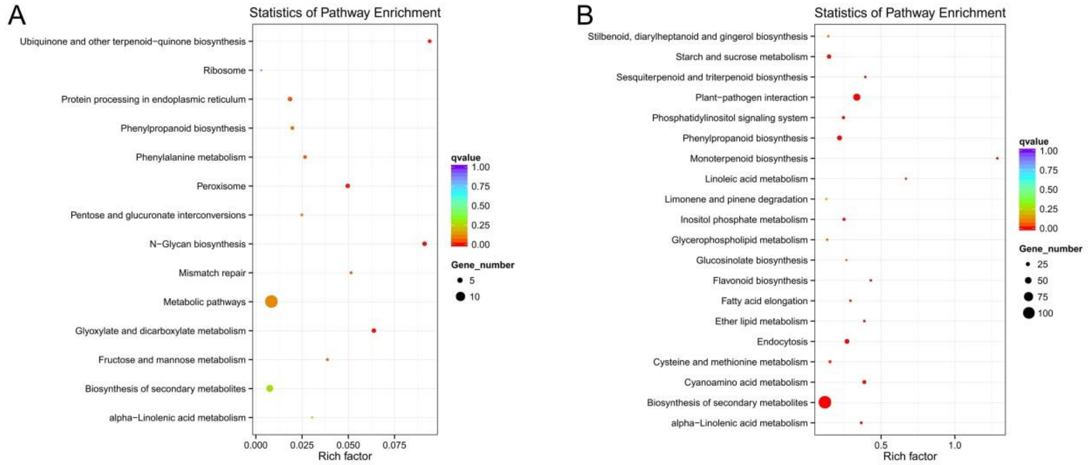


**Fig. S16 KEGG pathway enrichments of (A) expansion and (B) contraction genes in *D. oleifera*.**

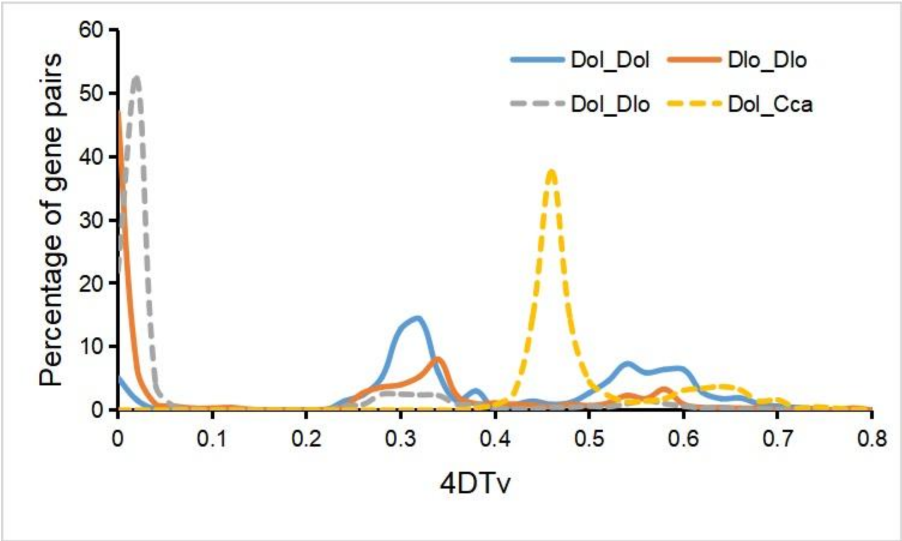


**Fig. S17 Whole-genome duplication analysis of *D. oleifera* genome.** Dol, Dlo and Cca represent *Diospyros oleifera*, *Diospyros lotus*, and *Coffea canephora*.

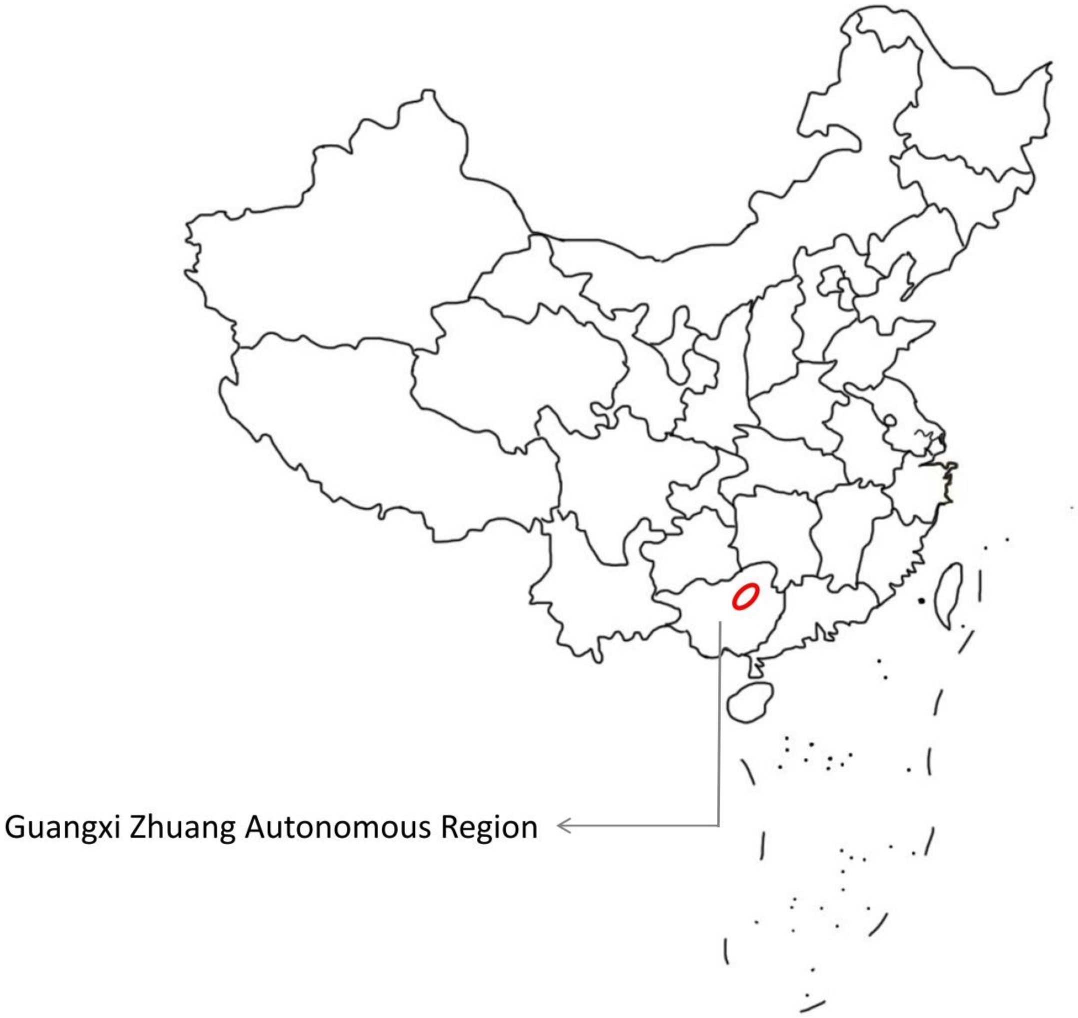


**Fig. S18 Sampling location.** The red circle in the map shows the origin of the natural populations investigated in this study.

```
CLUSTAL format alignment by MAFFT (v7.475)


Monoecy-D.oleif --------nnnnnnnnnnttgttnngtcttttatcaacaaatcacactcaaaatcaaact
Monoecy-D.kaki  nnnnnnnnnnnnnnnanntngttnnnnnttttatcaacaaataacactcaaaatcaaact
                                  * ***     ************** *****************

Monoecy-D.oleif aaatcaaacaaaatcaaaattaatttaaaatttacacaaaccacaaggaaaaattgaatt
Monoecy-D.kaki  ----------aaatcaaaattaatttaaaatttacacaaagcacaaggaaaaattgaatt
                          ****************************** *******************

Monoecy-D.oleif attttgcttttcatgtattaatataataaaattaagctccagtttagttatggaagagat
Monoecy-D.kaki  attttgctt------tattaatataataaaattaagctccagtttagttatggaagagat
                *********      *********************************************

Monoecy-D.oleif ttgttatgatgattaattaaaatgtaatgctccatccttgtggcatatctgaatatcacg
Monoecy-D.kaki  ttgttatgtngattaattaaaatgtaaagctccatccttgtggcatatctgaatatcatg
                ********  ***************** ******************************.*

Monoecy-D.oleif tcaaaatgc---------------------------------------------------
Monoecy-D.kaki  tcaaaaaactttatcacaagtgcccagggatataactaagtggtaaaggagggagttgta
                ****** .*

Monoecy-D.oleif ------------------------------------------------------------
Monoecy-D.kaki  aagatatagtcatactaaaaactgctagcatgatattgatttctagagaaaataatttct


Monoecy-D.oleif ------------------------------------------------------------
Monoecy-D.kaki  tctggggnggtttttagtaaatgttagtggacgcagtctcaaaccctgaccatttaagca


Monoecy-D.oleif ------------------------------------------------------------
Monoecy-D.kaki  tatctctcgatttacctccctcgtgataacccaggagtggattgatcaagggcgcgggtg


Monoecy-D.oleif --------------------------tttatcacataacttttgggaatcaccaaccaac
Monoecy-D.kaki  ttggagcataataataaaaaaaaaaatttatcacataacttttgggaatcaacaaccaac
                                          ************************* ********

Monoecy-D.oleif gatcagaagcattgagatttgtgc--aaaaaaacaagtgatgtgaacgttggcaattccg
Monoecy-D.kaki  gataagaagcattgagatttgtgcaaaaaaaaaacaagtgatgtgaacgttggcaattccg
                *** ********************  **********************************

Monoecy-D.oleif acttctggtgaaatgtcaaattccactg-aaacacttgaatagagataacgnnangataa
Monoecy-D.kaki  acttctggtgaaatgtcaaattccactgaaaacacttgaatagagataacgcaannnnna
                **************************** **********************  *     *

Monoecy-D.oleif gccnna
Monoecy-D.kaki  -----n
```

**Fig. S19 The sequence of 5' untranslated region of *OGI* between monoecious *D. oleifera* (above) and monoecious *D. kaki* cultivar 'Taishuu' (below).**

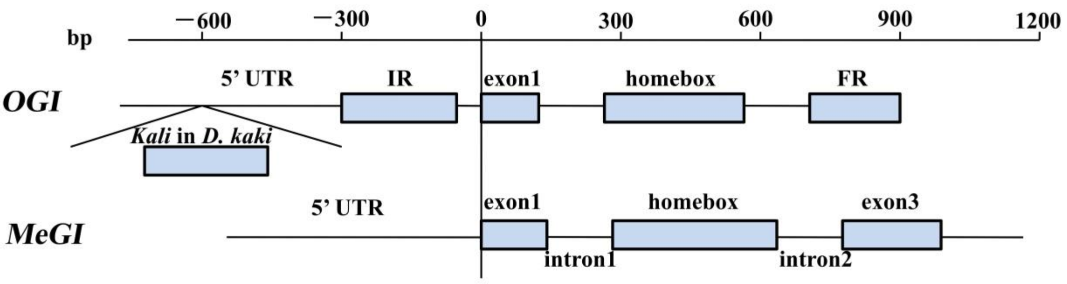


**Fig. S20 Comparison of *OGI* and *MeGI* gene model.** UTR: untranslated regions; IR: inverted repeat; FR: forward repeat.

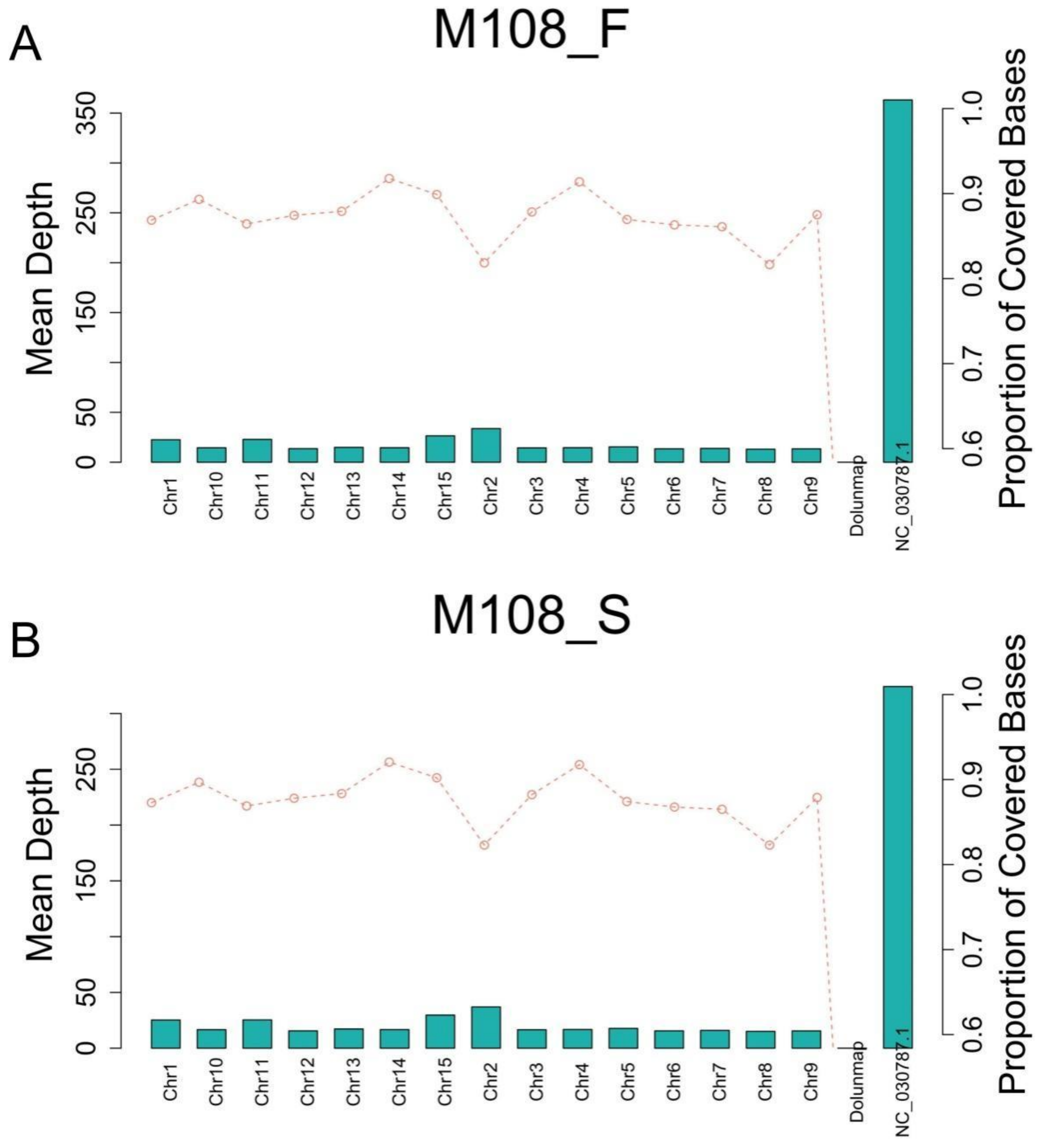


**Fig. S21 Distribution of BS-seq reads on each chromosome in (A) M108_F and (B) M108_S.** The histogram represents the mean coverage depth. The Scatter plot represents the proportion of covered bases. M108_F and M108_S represents female floral buds and stems of immature flowering shoots obtained from the monoecious #108 tree (the same as in Fig. S22). Dolunmap reprents the male-unmapped sequences assembled in this study, and NC_030787.1 reprents the chloroplast gemome of *D. oleifera*, which is consistent in the whole text.

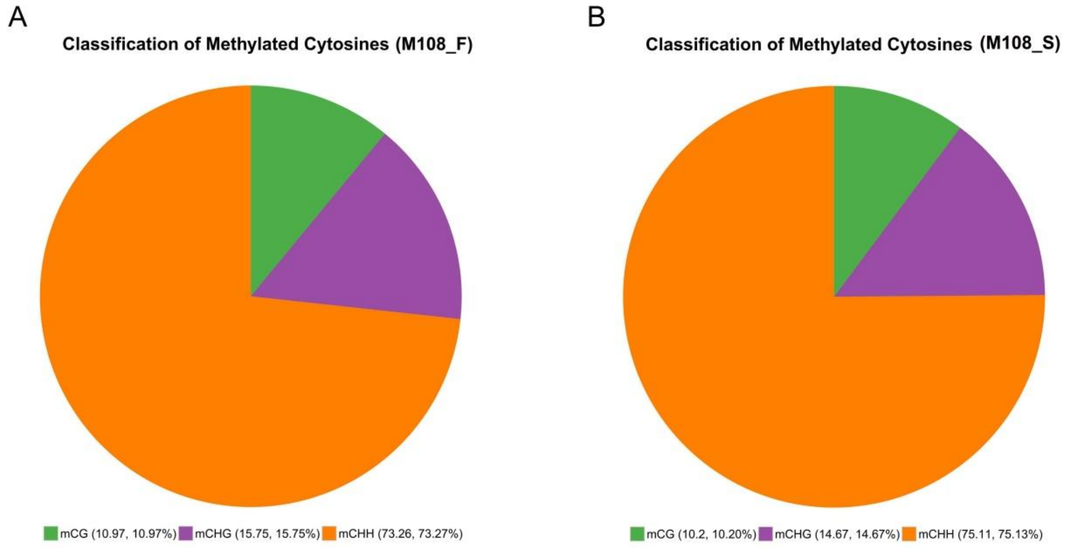


**Fig. S22 Constitution of methylated cytosines in (A) M108_F and (B) M108_S.**

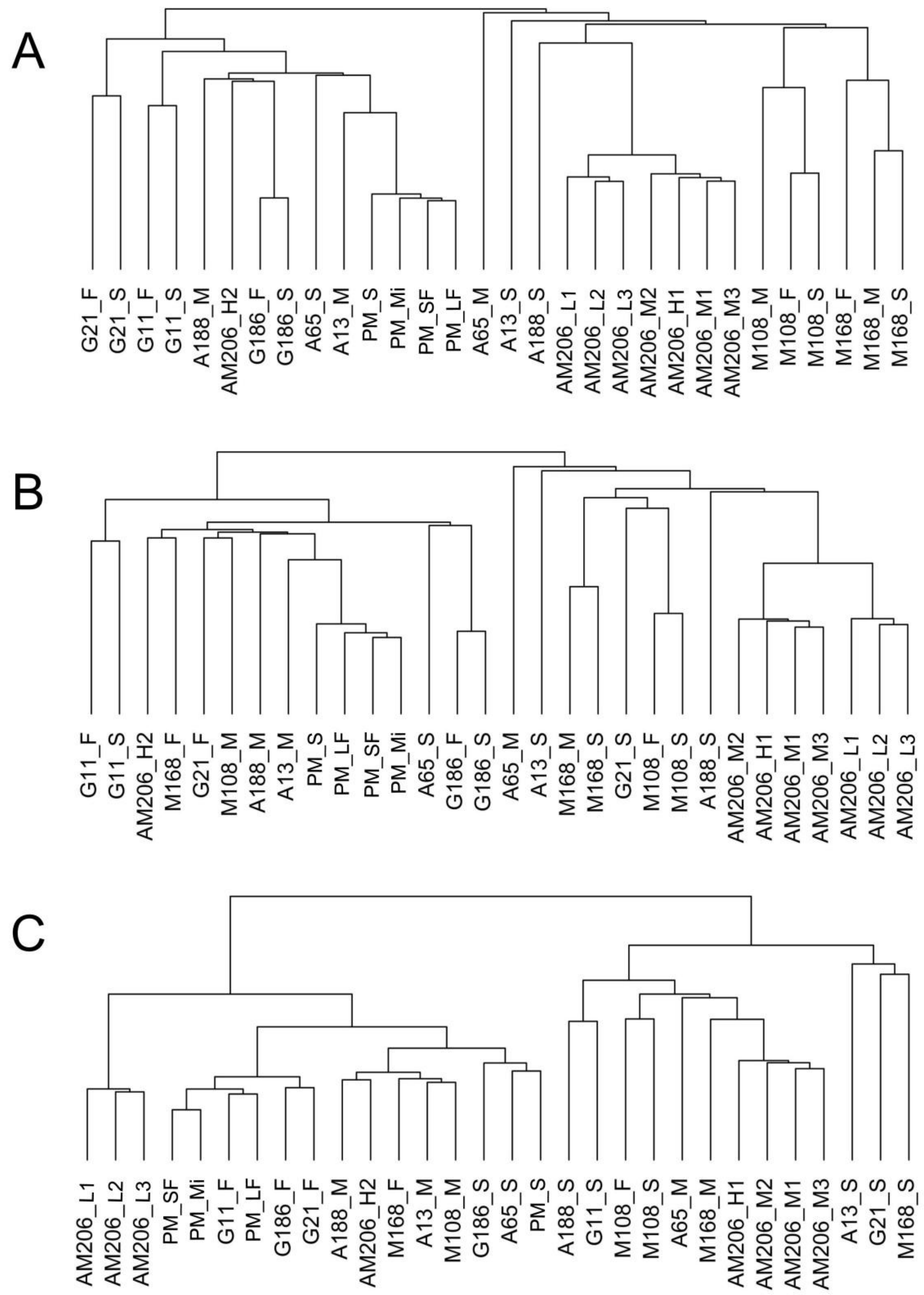


**Fig. S23 Hierarchical clustering based on PCA analysis on the methylation levels of (A) CG, (B) CHG, and (C) CHH subcontexts among all the samples.**

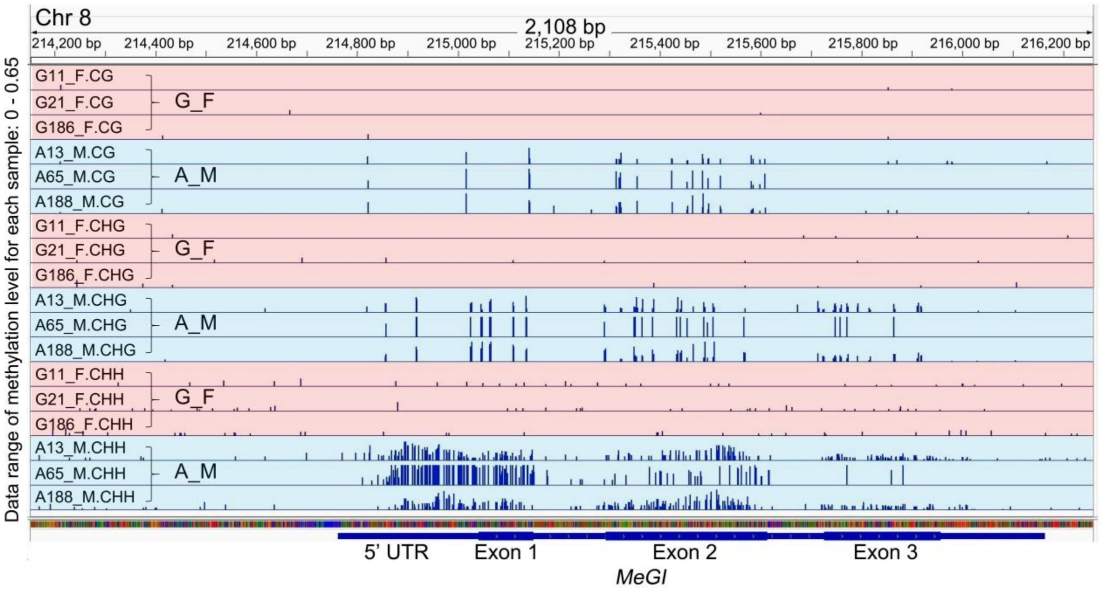


**Fig. S24 Methylation levels of the *MeGI* genomic region in the floral buds obtained from single-sex *D. oleifera*.**

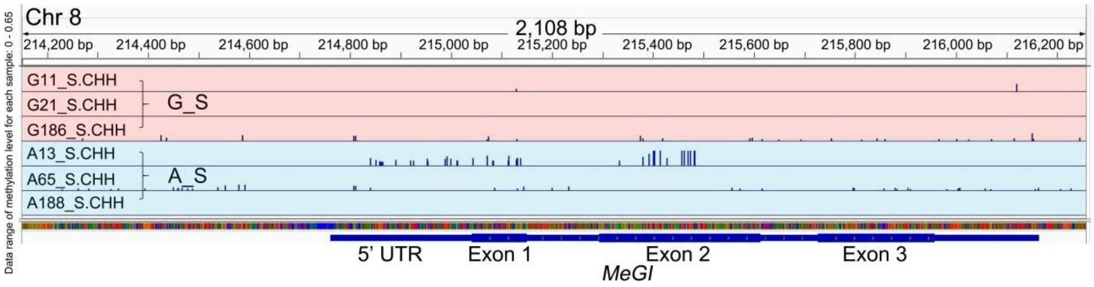


**Fig. S25 Methylation levels of the *MeGI* genomic region in the immature stems of flowering shoots obtained from single-sex *D. oleifera*.**

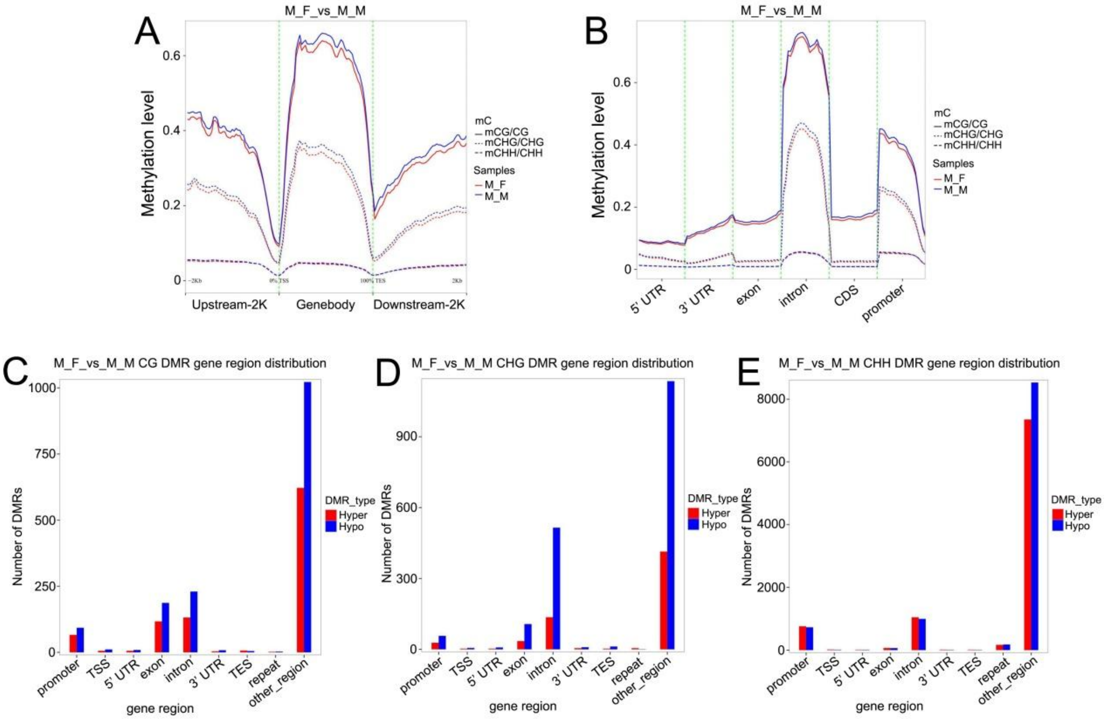


**Fig. S26 Comparison of DNA methylation between female and male flowers in monoecious plants. (A-B)** Comparison of methylation levels of CG, CHG, and CHH subcontexts between M_F and M_M in **(A)** gene body as well as up and downstream regions, and in **(B)** regions with different genomic features. **(C-E)** Number of DMRs in **(C)** CG, **(D)** CHG, and **(E)** CHH subcontexts between M_F and M_M in regions with different genomic features. “DMR” represents differentially methylated regions; “Hyper DMR” and "hypo DMR" represent regions with higher and lower methylation levels in the former sample compared with the latter one in the comparative combination, respectively. This criterion is consistent in the following figures.

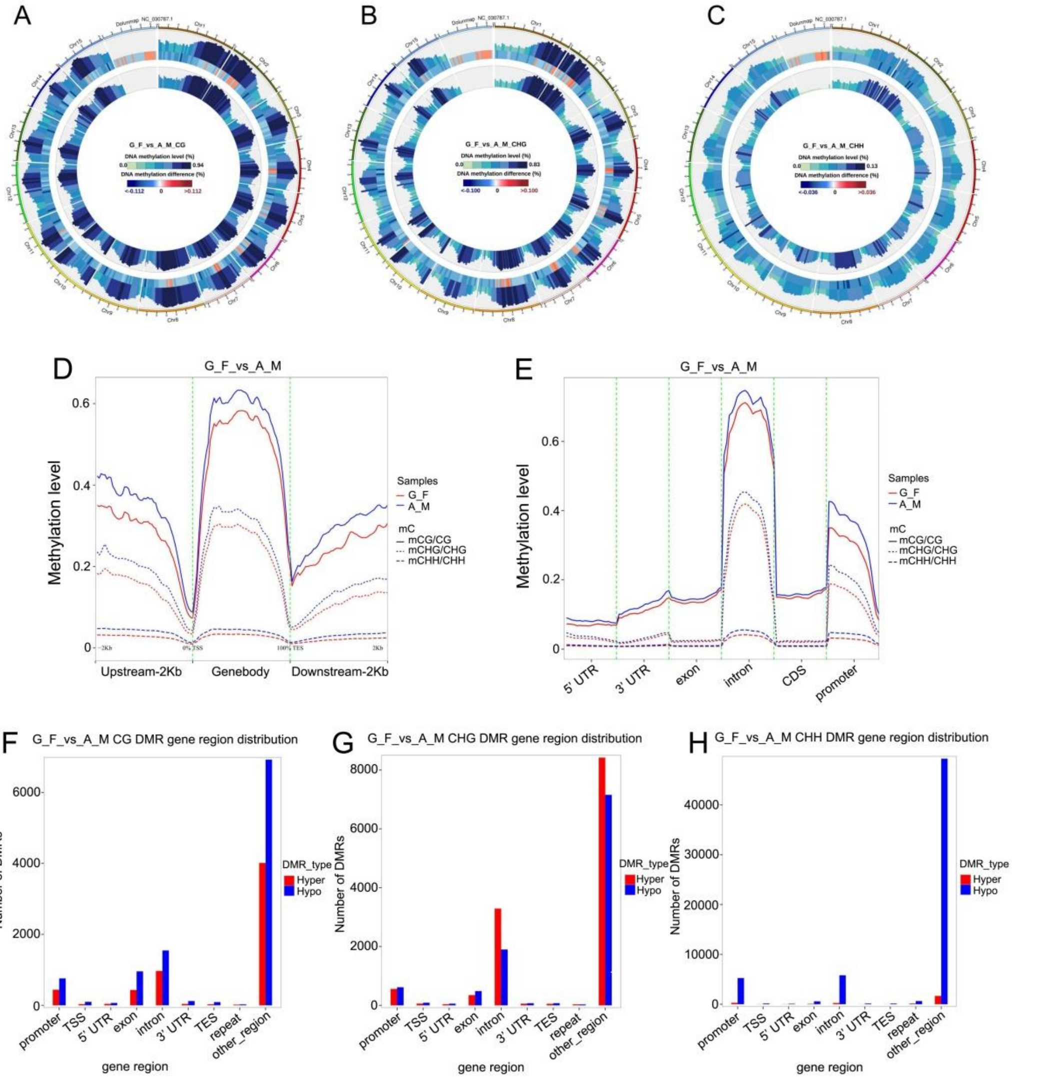


**Fig. S27 Comparison of DNA methylation levels between female and male flowers in single-sex plants. (A-C)** Whole-genome comparison of methylation levels in the **(A)** CG, **(B)** CHG, and **(C)** CHH subcontexts between G_F and A_M. Tracks from outside to inside: methylation level of G_F; different methylation levels between G_F and A_M, where red and blue represent higher and lower methylation levels in G_F than in A_M, respectively; methylation level of A_M. **(D-E)** Comparison of methylation levels of the CG, CHG, and CHH subcontexts between G_F and A_M in **(D)** gene body as well as up and downstream regions, and in **(E)** regions with different genomic features. **(F-H)** Number of DMRs in the **(F)** CG, **(G)** CHG, and **(H)** CHH subcontexts between G_F and A_M in regions with different genomic features.

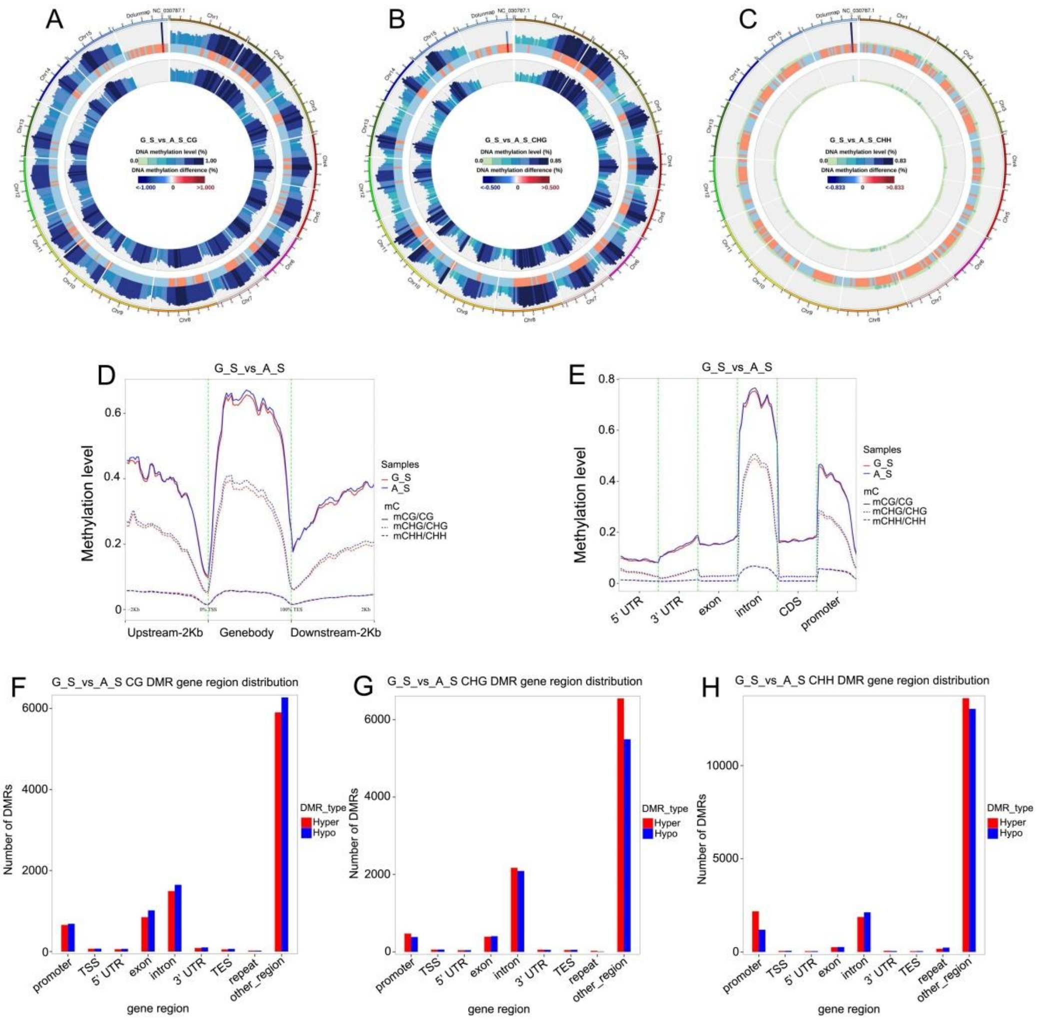


**Fig. S28 Comparison of DNA methylation levels between male and female immature stems in single-sex plants. (A-C)** Whole-genome comparison of methylation levels in the **(A)** CG, **(B)** CHG, and **(C)** CHH subcontexts between G_S and A_S. Tracks from outside to inside: methylation level of G_S; different methylation level between G_S and A_S, where red and blue represent higher and lower methylation levels in G_S than in A_S, respectively; methylation level of A_S. **(D-E)** Comparison of methylation levels of the CG, CHG, and CHH subcontexts between G_S and A_S in **(D)** gene body as well as up and downstream regions, and in **(E)** regions with different genomic features. **(F-H)** Number of DMRs in the **(F)** CG, **(G)** CHG, and **(H)** CHH subcontexts between G_S and A_S in regions with different genomic features.

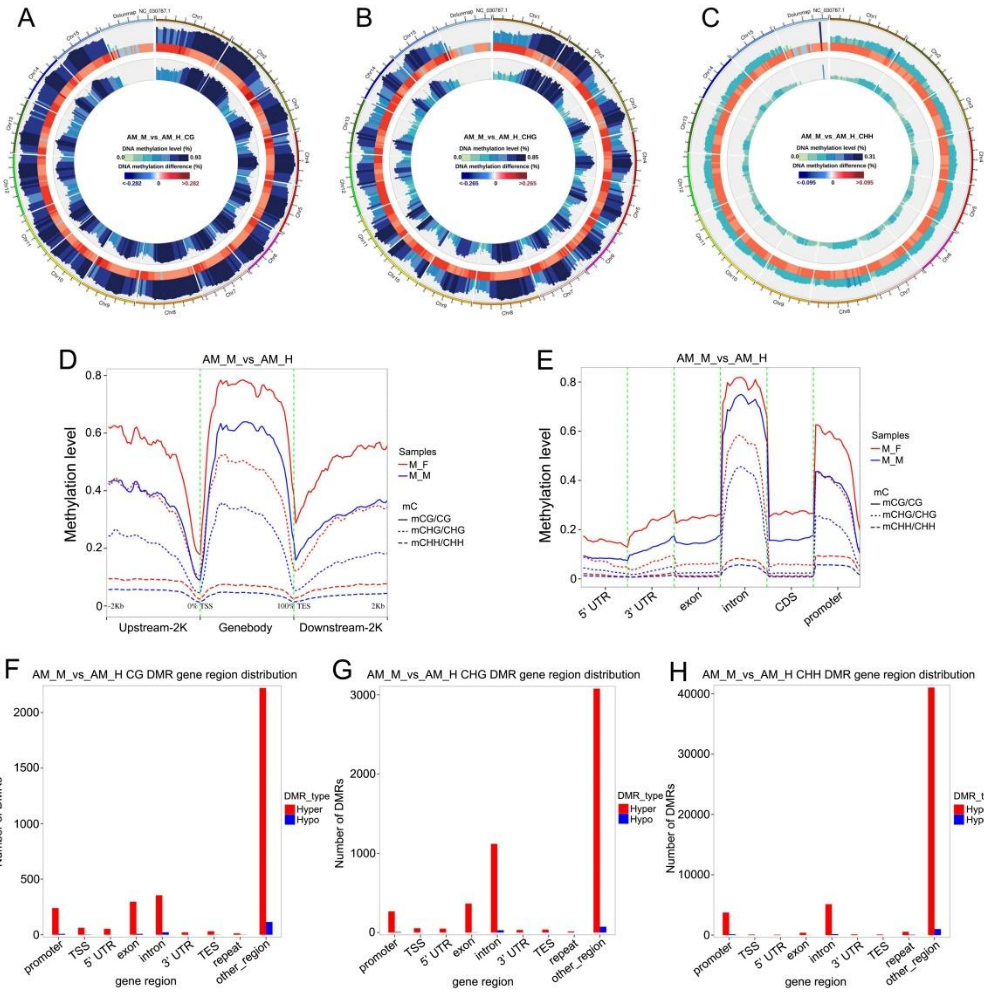


**Fig. S29 Comparison of DNA methylation levels between male and hermaphroditic flowers in andromonoecios plants.** **(A-C)** Whole-genome comparison of methylation levels in the **(A)** CG, **(B)** CHG, and **(C)** CHH subcontexts between AM_M and AM_H. Tracks from outside to inside: methylation level of AM_M; different methylation level between AM_M and AM_H, where red and blue represent higher and lower methylation levels in AM_M than AM_H, respectively; methylation level of AM_H. **(D-E)** Comparison of methylation levels of the CG, CHG, and CHH subcontexts between AM_M and AM_H in **(D)** gene body as well as up and downstream regions, and in **(E)** regions with different genomic features. **(F-H)** Number of DMRs in the **(F)** CG, **(G)** CHG, and **(H)** CHH subcontexts between AM_M and AM_H in regions with different genomic features.

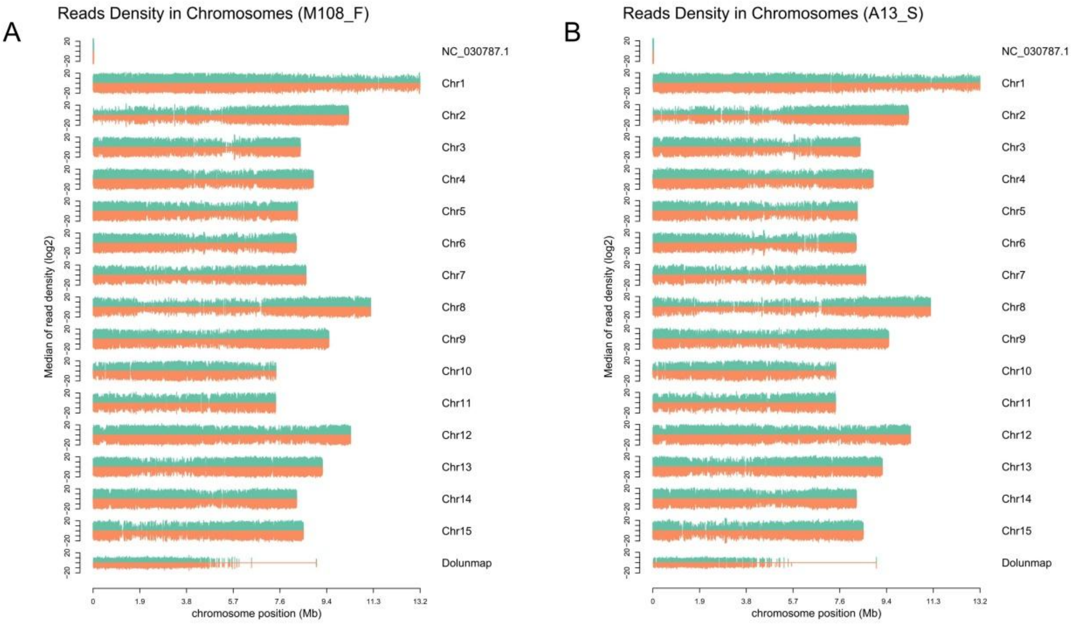


**Fig. S30 Distribution of RNA reads on chromosomes in (A) M108_F and (B) A13_S.** M108_F represents female floral buds obtained from the monoecious #108 tree, and A13_S represents immature stems of flowering shoots obtained from the androecious #13 tree (the same as in Fig. S31).

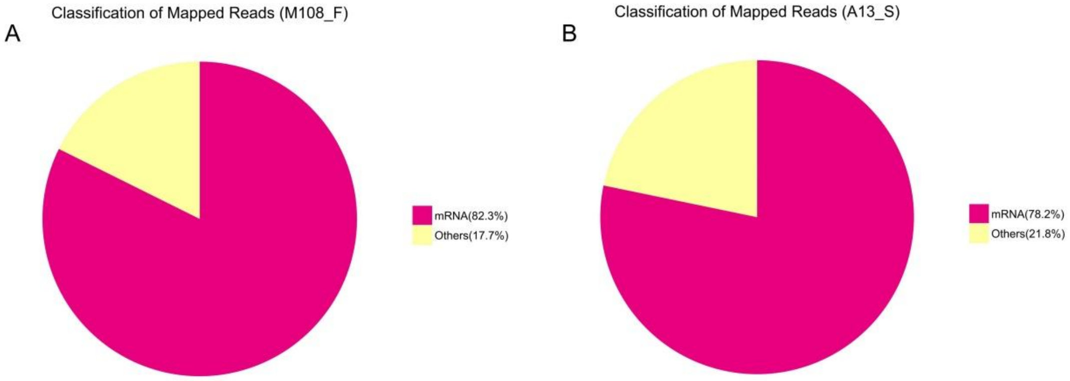


**Fig. S31 Classification of mapped rRNA-depleted RNA-seq reads in (A) M108_F and (B) A13_S.**

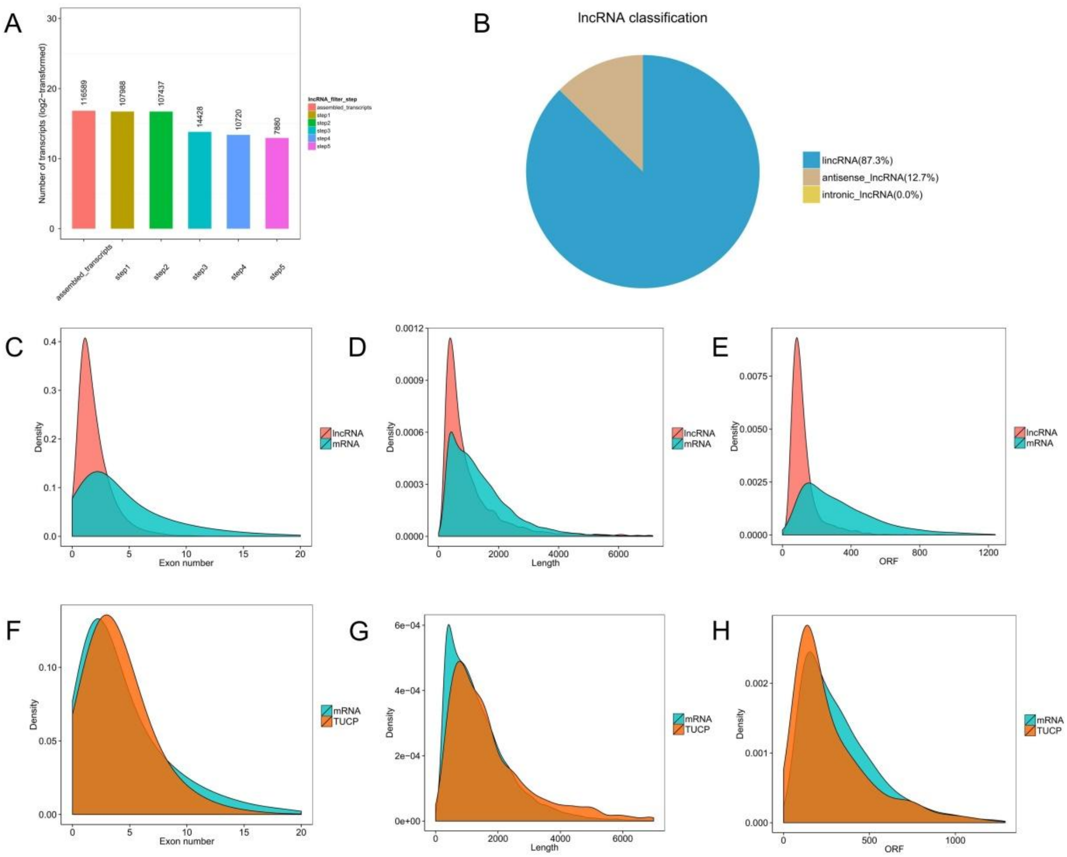


**Fig. S32 Screening of lncRNAs and basic characteristics of lncRNA and TUCP. (A)** Five steps for lncRNAs screening. **(B)** Constitution of lncRNAs. **(C)** Number density of mRNA and lncRNA exons. **(D)** Length density of mRNA and lncRNA. **(E)** ORF length density of mRNA and lncRNA. **(F)** Number density of mRNA and TUCP exons. **(G)** Length density of mRNA and TUCP. **(H)** ORF length density of mRNA and TUCP.

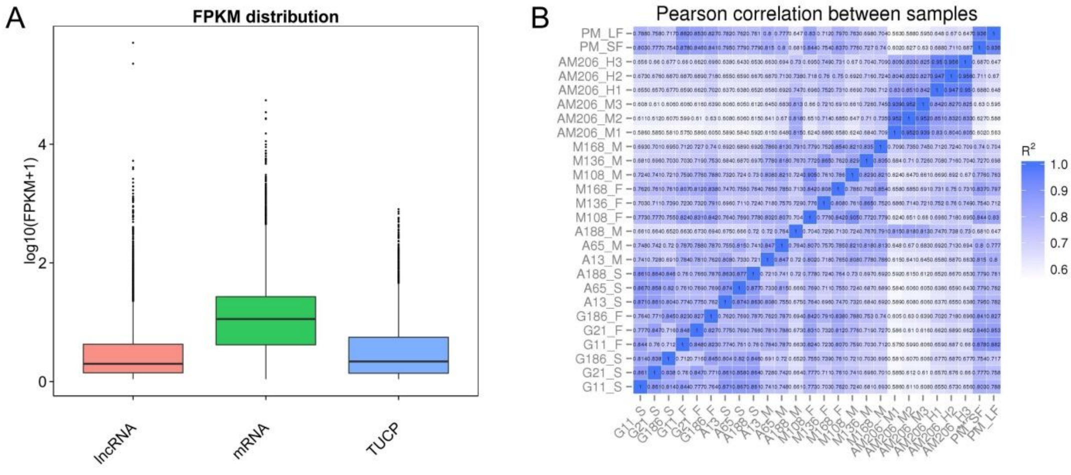


**Fig. S33 (A) FPKM distribution of mRNA, lncRNA, and TUCP and (B) Pearson correlation analysis of expression levels among samples.**

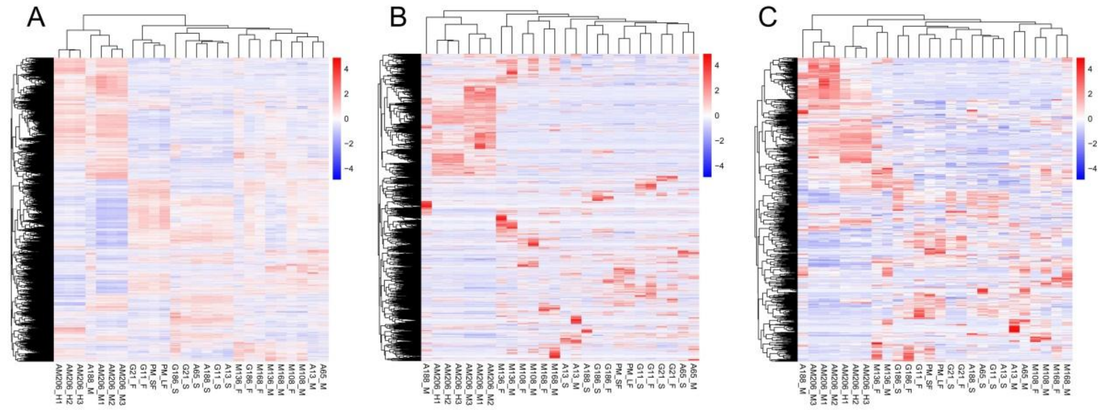


**Fig. S34 Clustering of samples based on the FPKM of all the differentially expressed (A) mRNAs, (B) lncRNAs, and (C) TUCPs.**

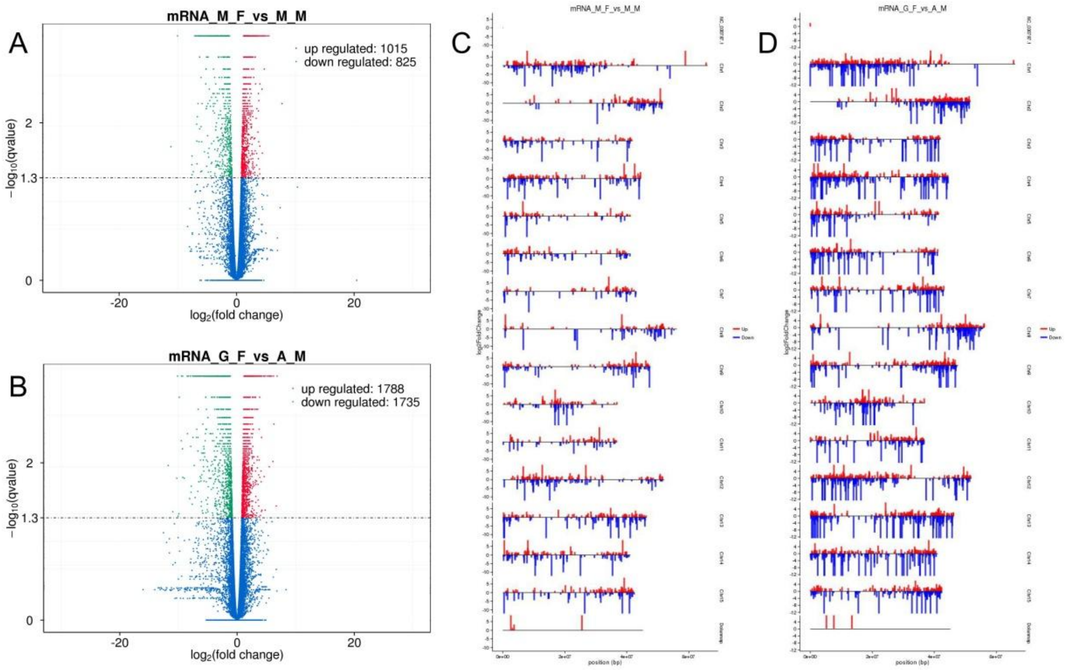


**Fig. S35 The volcano plot of differentially expressed mRNAs (DEGs) in (A) M_F compared with M_M, and (B) G_F compared with A_M. Distribution of DEGs on chromosomes in (C) M_F compared with M_M, and (D) G_F compared with A_M.**

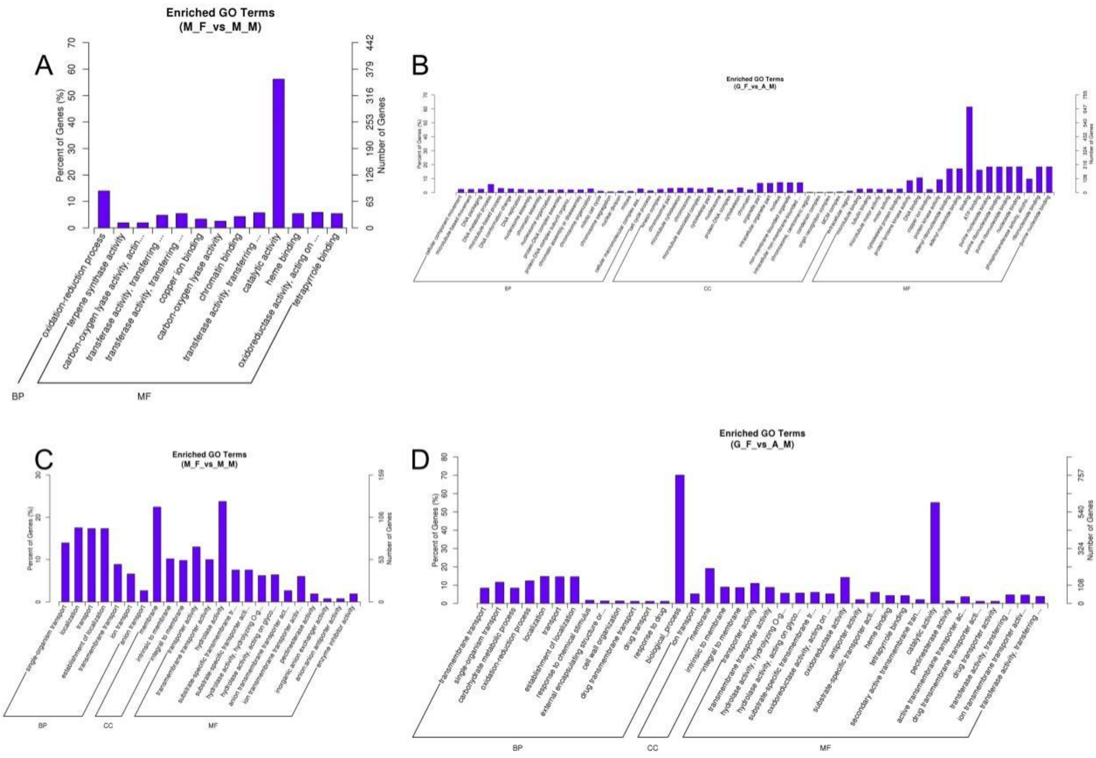


**Fig. S36 Significant GO enrichment of upregulated DEGs in (A) M_F vs. M_M and (B) G_F vs. A_M. Significant GO enrichment of downregulated DEGs in (C) M_F vs. M_M and (D) G_F vs. A_M.** "M_F vs. M_M", "G_F vs. A_M", and "AM_M vs. AM_H" represent M_F compared with M_M, G_F compared with A_M, and AM_M compared with AM_H, respectively, in the whole text.

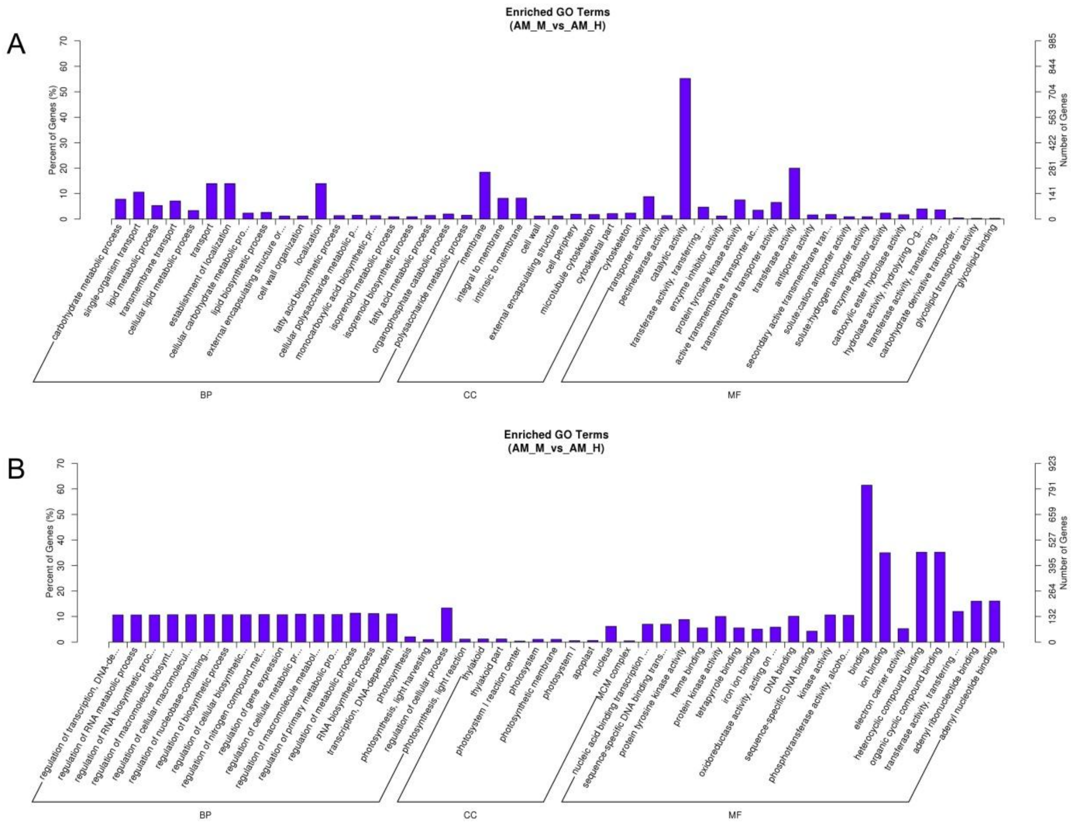


**Fig. S37 GO enrichment of (A) up- and (B) downregulated DEGs in AM_M vs. AM_H.**

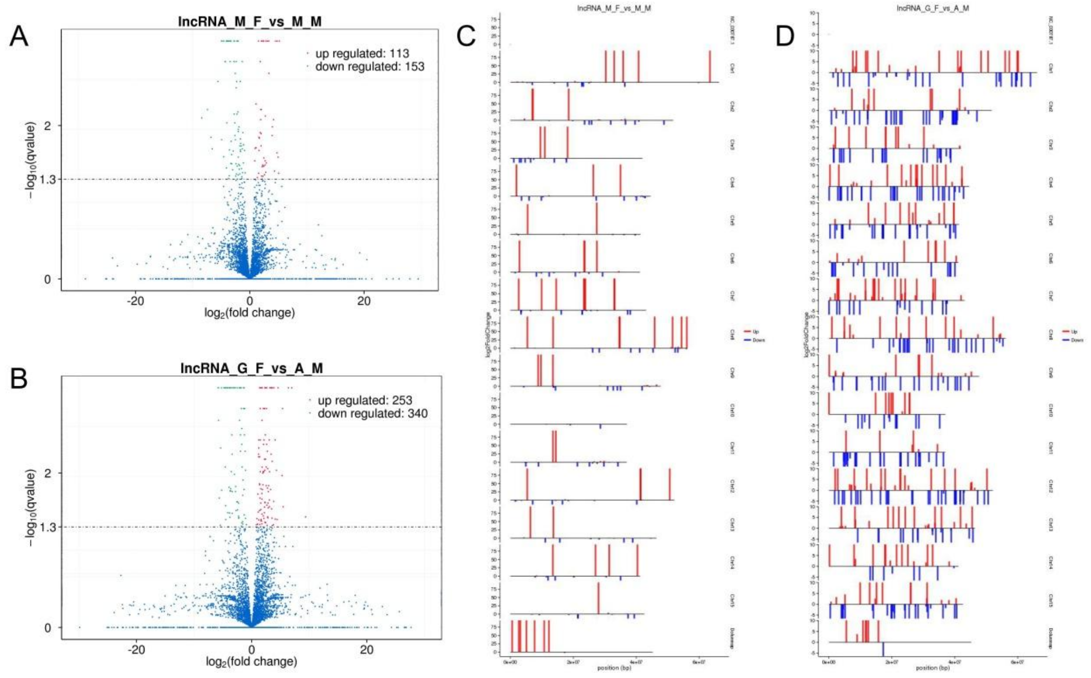


**Fig. S38 The volcano plot of differentially expressed lncRNAs (DELs) in (A) M_F vs. M_M, and in (B) G_F vs. A_M. Distribution of DELs on chromosomes in (C) M_F vs. M_M, and (D) G_F vs. A_M.**

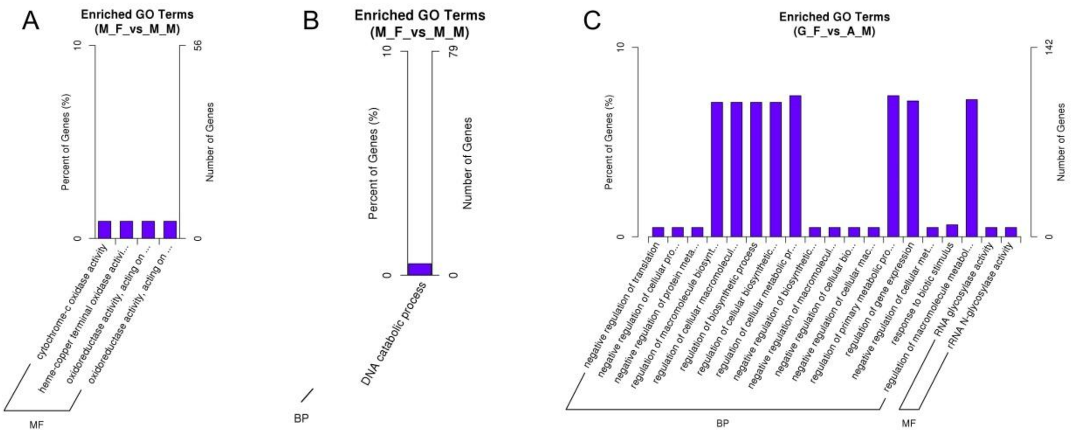


**Supplementary Fig. S39 Significant GO enrichment of mRNA co-located with (A) up- and (B) downregulated DELs in M_F vs. M_M. (C) Significant GO enrichment of mRNA co-located with downregulated DELs in G_F vs. A_M.**

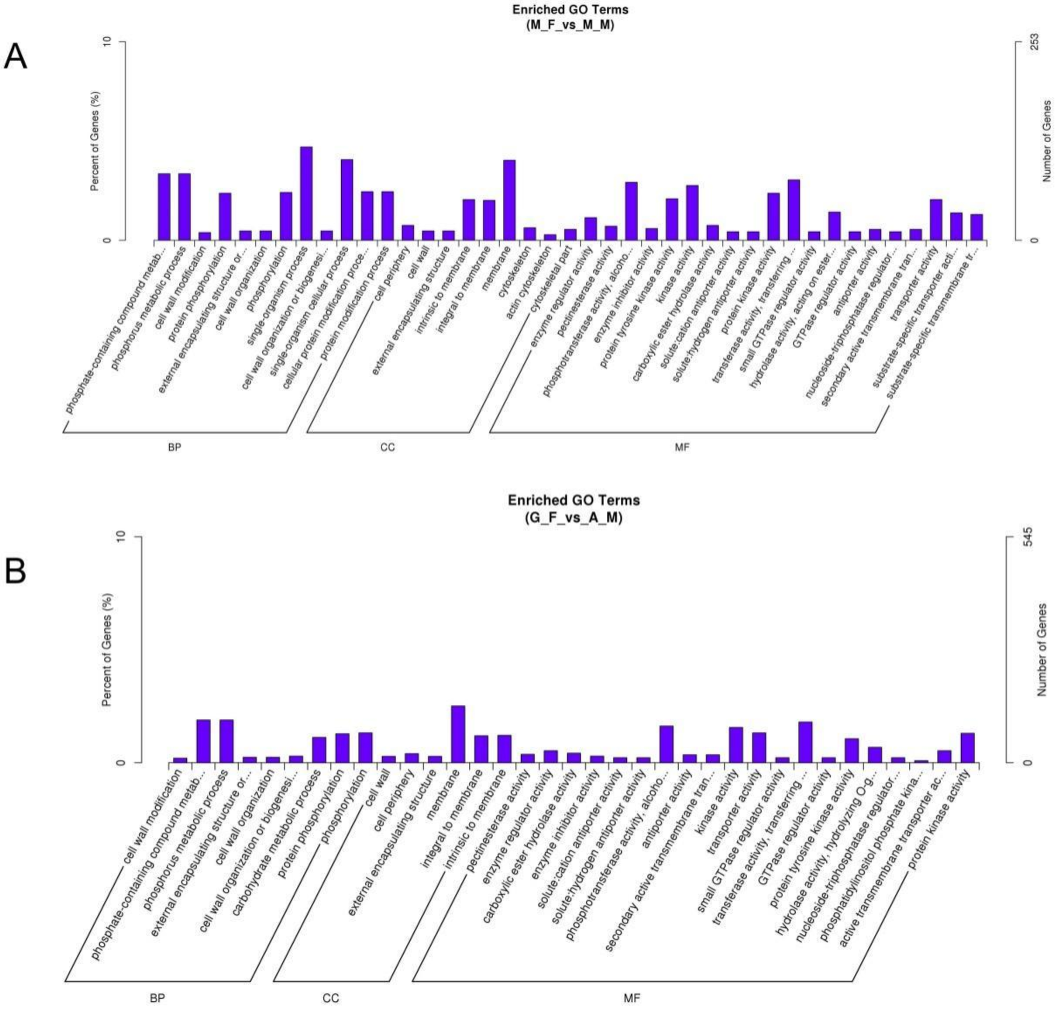


**Fig. S40 Significant GO enrichment of mRNA coexpressed with downregulated DELs in (A) M_F vs. M_M and (B) G_F vs. A_M.**

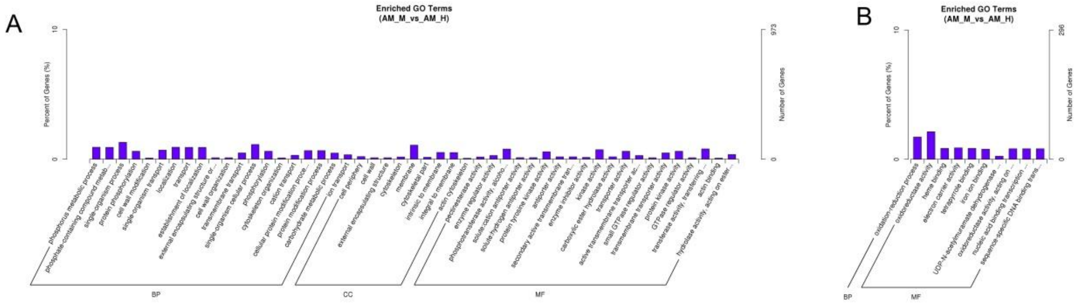


**Fig. S41 Significant GO enrichment of mRNA coexpressed with (A) up- and (B) downregulated DELs in AM_M vs. AM_H.**

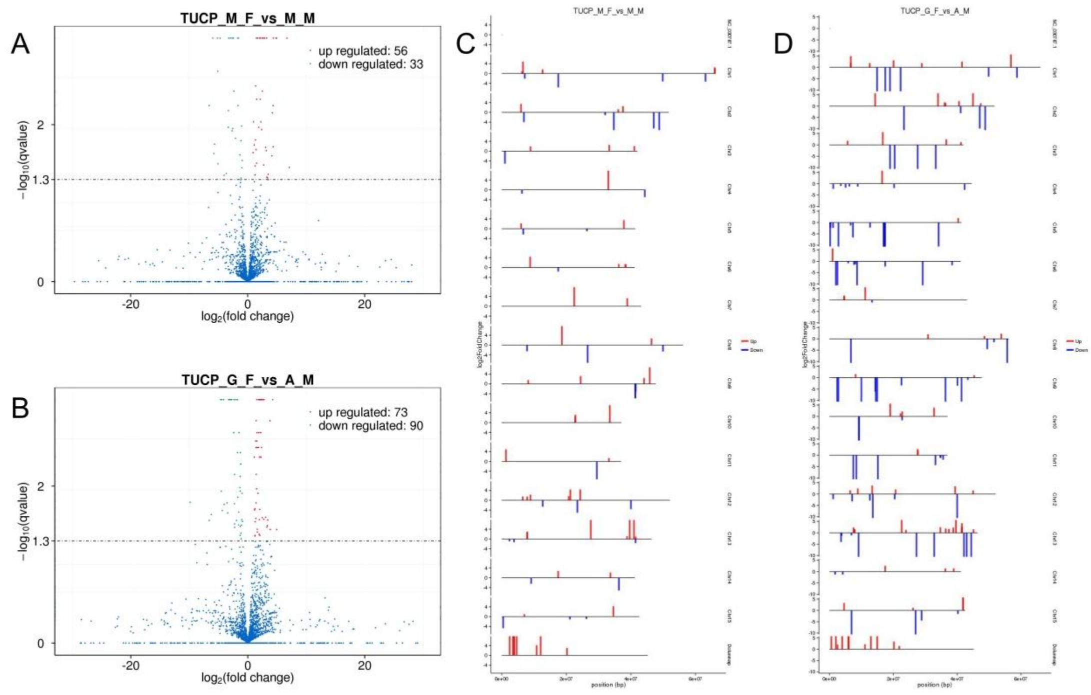


**Fig. S42 The volcano plot of differentially expressed TUCPs (DETs) in (A) M_F vs. M_M, and in (B) G_F vs. A_M. Distribution of DETs on chromosomes in (C) M_F vs. M_M, and in (D) G_F vs. A_M.**

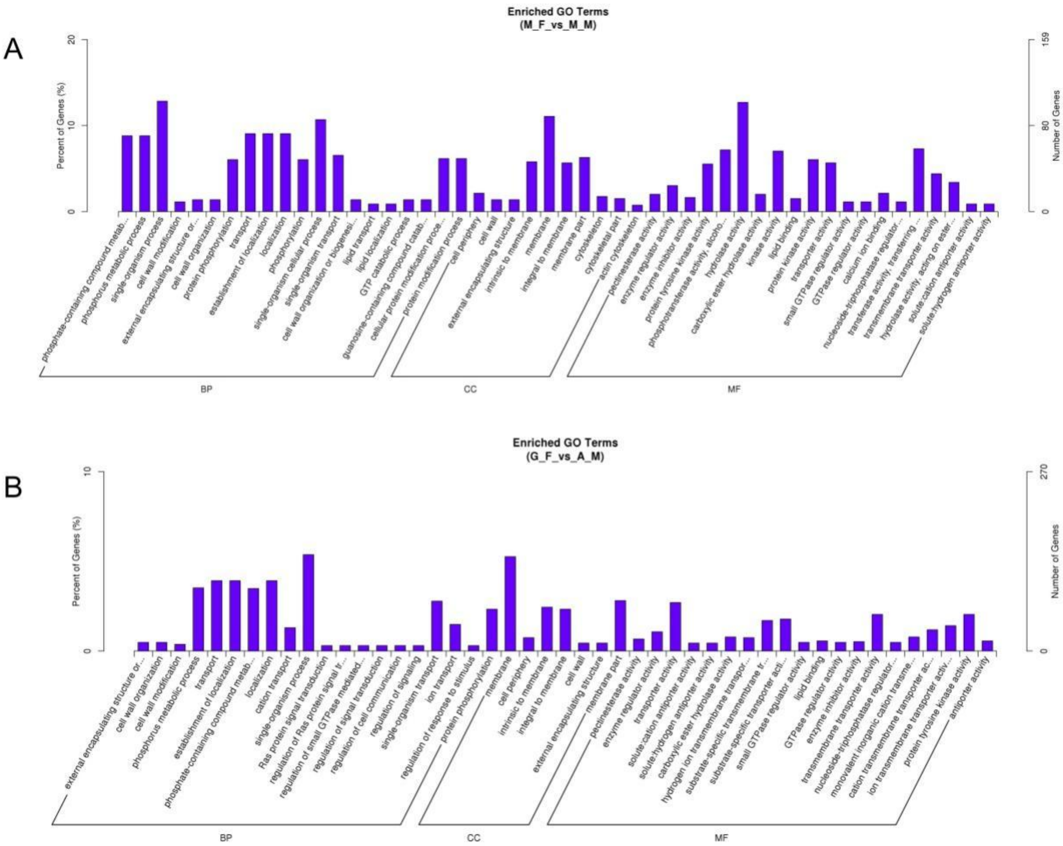


**Fig. S43 Significant GO enrichment of mRNA coexpressed with downregulated DETs in (A) M_F vs. M_M and (B) G_F vs. A_M.**

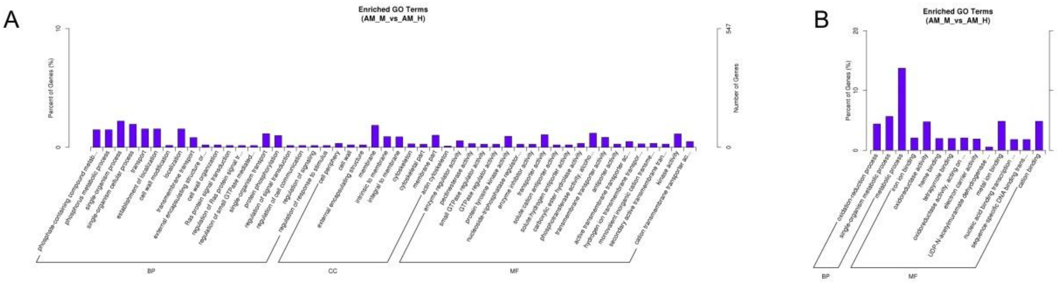


**Fig. S44 Significant GO enrichment of mRNA coexpressed with (A) up- and (B) downregulated DETs in AM_M vs. AM_H.**

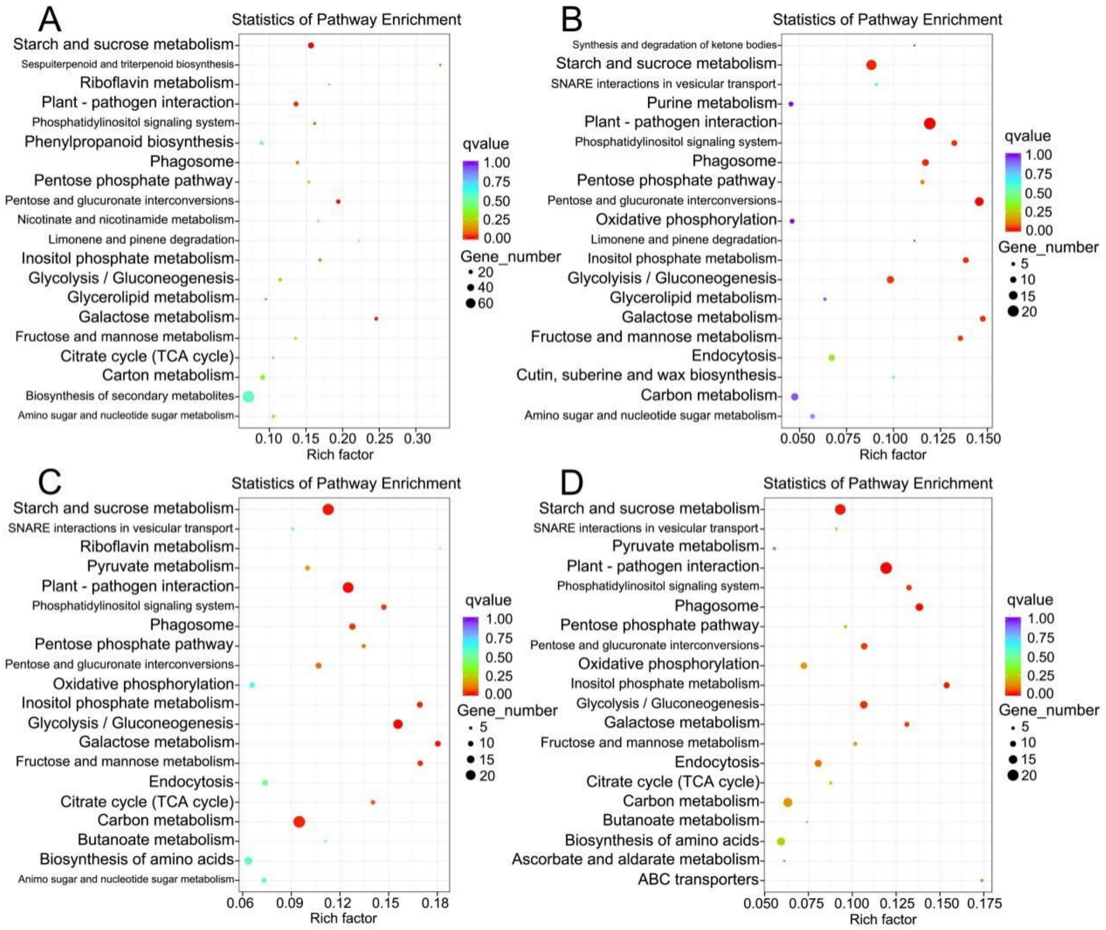


**Fig. S45 KEGG pathway enrichment of mRNA co-expressed with downregulated lncRNAs in (A) G_F vs. A_M, and in (B) M_F vs. M_M. KEGG pathway enrichment of mRNA co-expressed with downregulated (C) DELs and (D) DETs in AM_H vs. AM_M.**

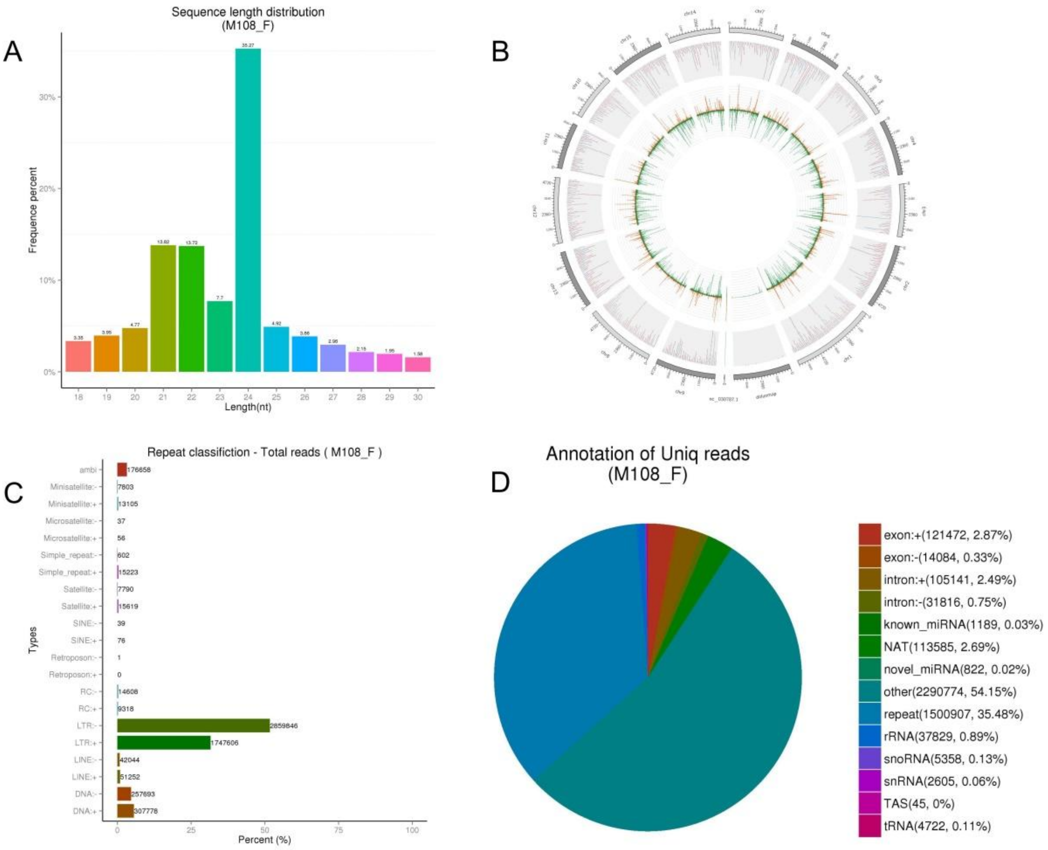


**Fig. S46 (A) Length distribution of small RNAs in M108_F. (B) Distribution of small RNAs reads obtained from M108_F on the genome. (C) Classification of repeat sequence in M108_F. (D) Annotation of unique reads in M108_F. M108_F represents female floral buds obtained from the monoecious #108 tree.** Dolunmap and NC_030787.1 in **(B)** reprent the male-unmapped sequences and the chloroplast gemome of *D. oleifera*, respectively. The "+" and "-"in (C) represent forward and reverse sequence, respectively.

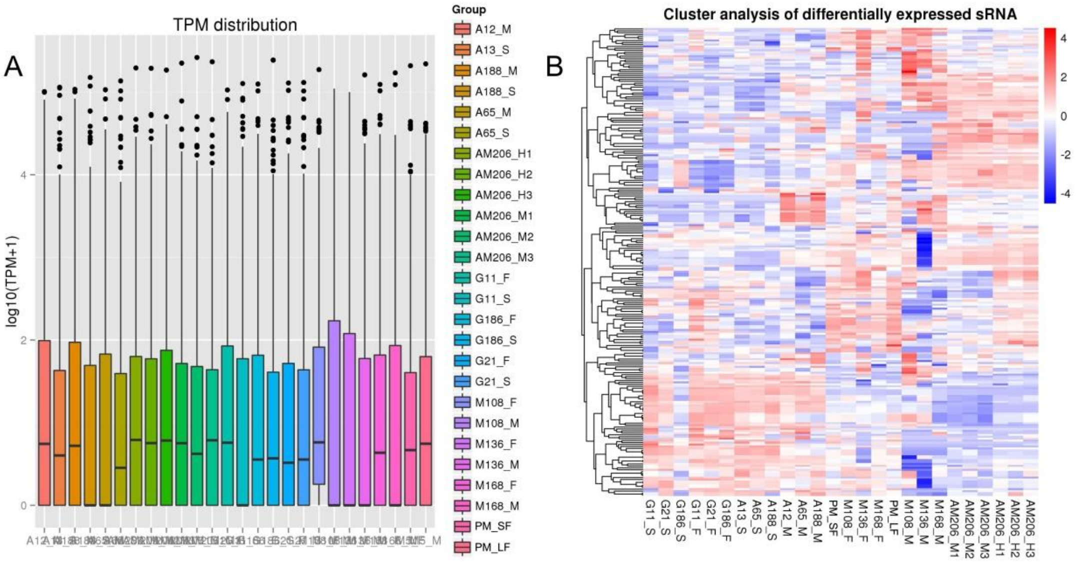


**Fig. S47 (A) Expression levels of small RNAs in all the samples. (B) Clustering of samples based on the TPM of all the differentially expressed miRNAs (DEMs).**

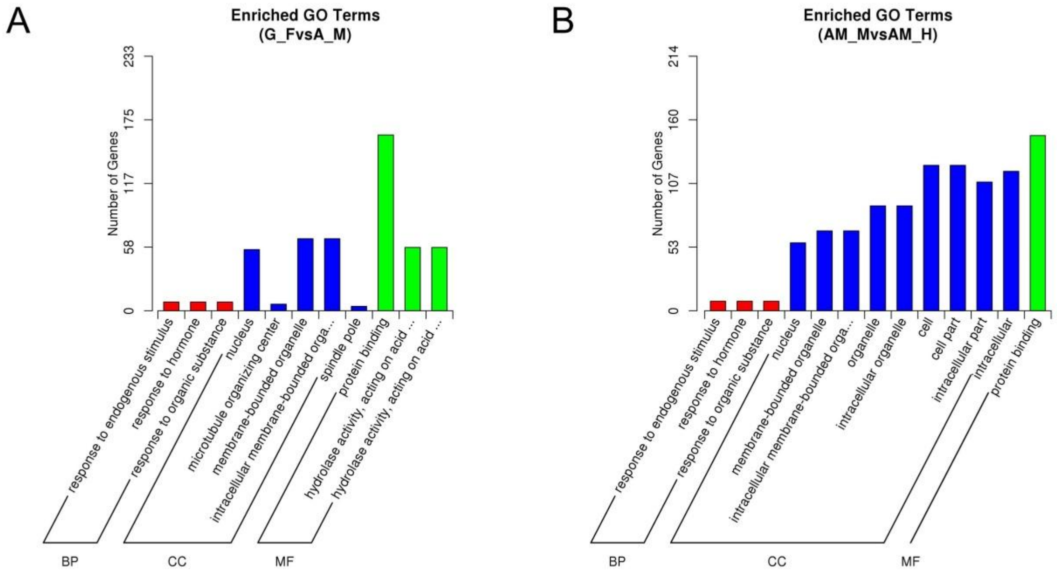


**Fig. S48 Significant GO enrichment of mRNA targeted by DEMs in (A) G_F vs. A_M, and (B) AM_M vs. AM_H.**

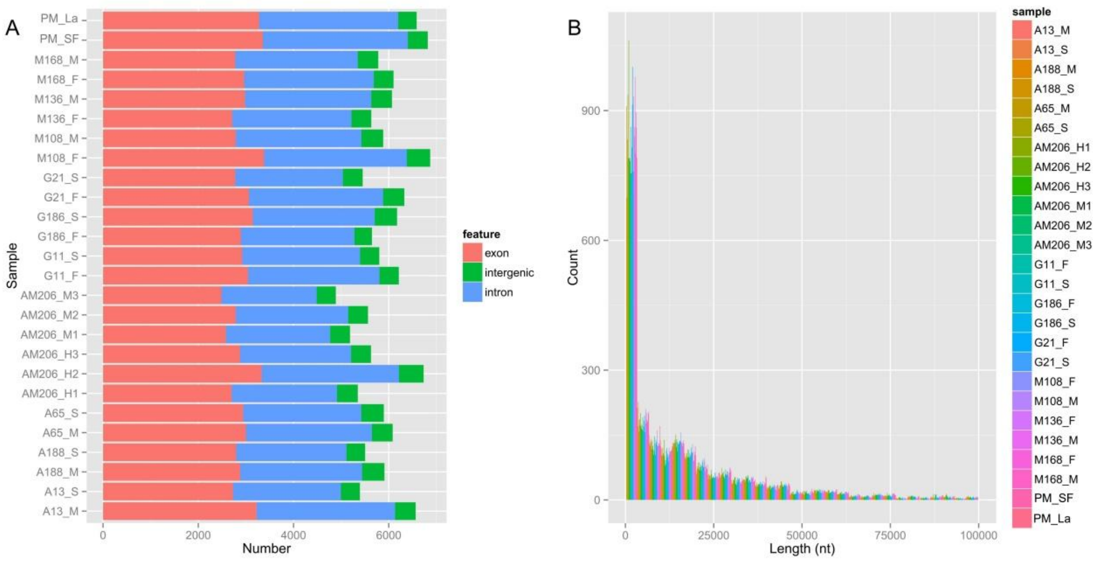


**Fig. S49 (A) Genomic features of the host sequences of circRNAs. (B) Length distribution of circRNAs identified in this study.**

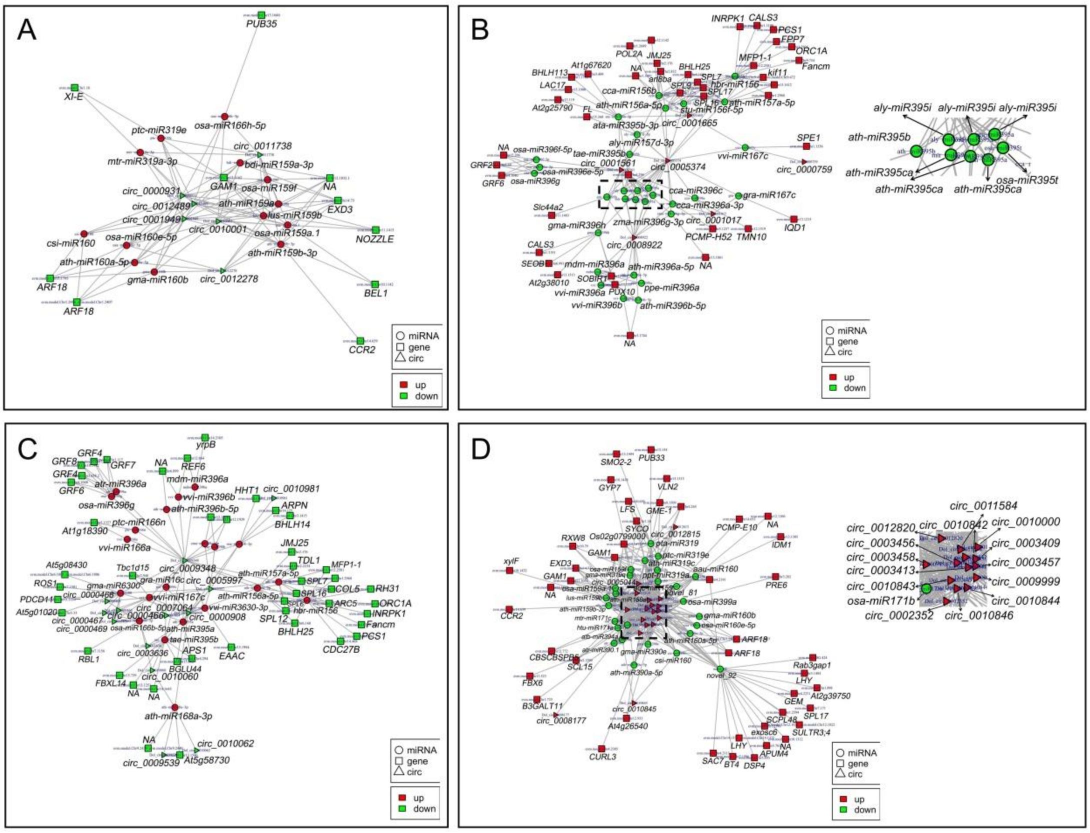


**Fig. S50 Competing RNA (ceRNA) networks constructed based on differentially expressed circRNAs (DECs), DEMs, and their DEGs targets in G_F vs. A_M and AM_M vs. AM_H. (A)** A ceRNA network constructed based on 6 downregulated DECs, 12 upregulated DEMs, and their 11 downregulated DEGs targets in G_F compared with A_M. **(B)** A ceRNA network constructed based on 6 upregulated DECs, 30 downregulated DEMs, and their 46 upregulated DEGs targets in G_F compared with A_M. **(C)** A ceRNA network constructed based on 13 downregulated DECs, 20 upregulated DEMs and their 50 downregulated DEGs targets in AM_M compared with AM_H. **(D)** A ceRNA network was constructed based on 19 upregulated DECs, 25 downregulated DEMs and their 49 upregulated DEGs targets in AM_M compared with AM_H.

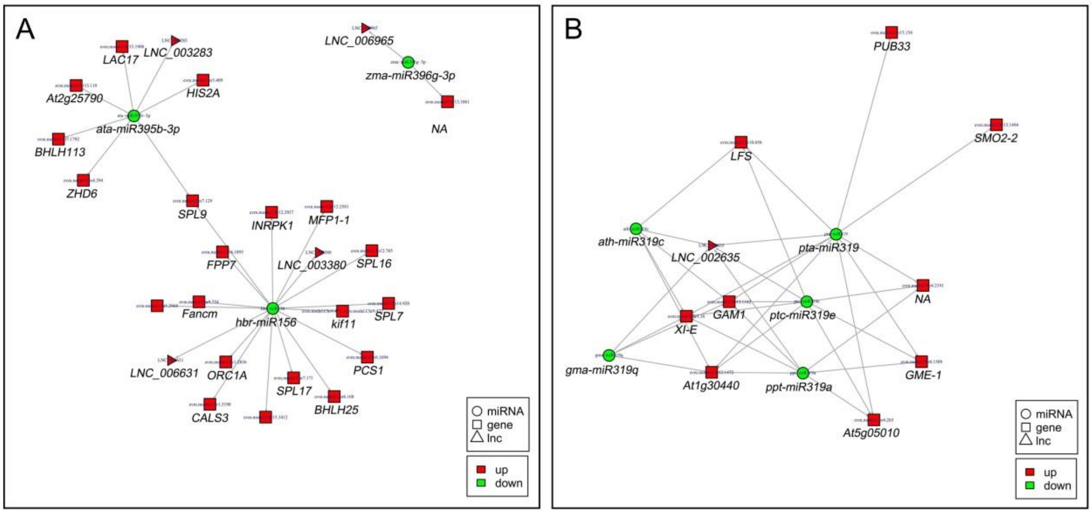


**Fig. S51 ceRNA Networks constructed based on DELs, DEMs, and their DEGs targets in**

**G_F vs. A_M and AM_M vs. AM_H. (A)** A ceRNA network constructed based on 4 upregulated DELs, 3 downregulated DEMs, and their 21 upregulated DEGs targets in G_F compared with A_M. **(B)** A ceRNA network constructed based on an upregulated DEL, 5 downregulated DEMs and their 9 upregulated DEGs targets in AM_M compared with AM_H.

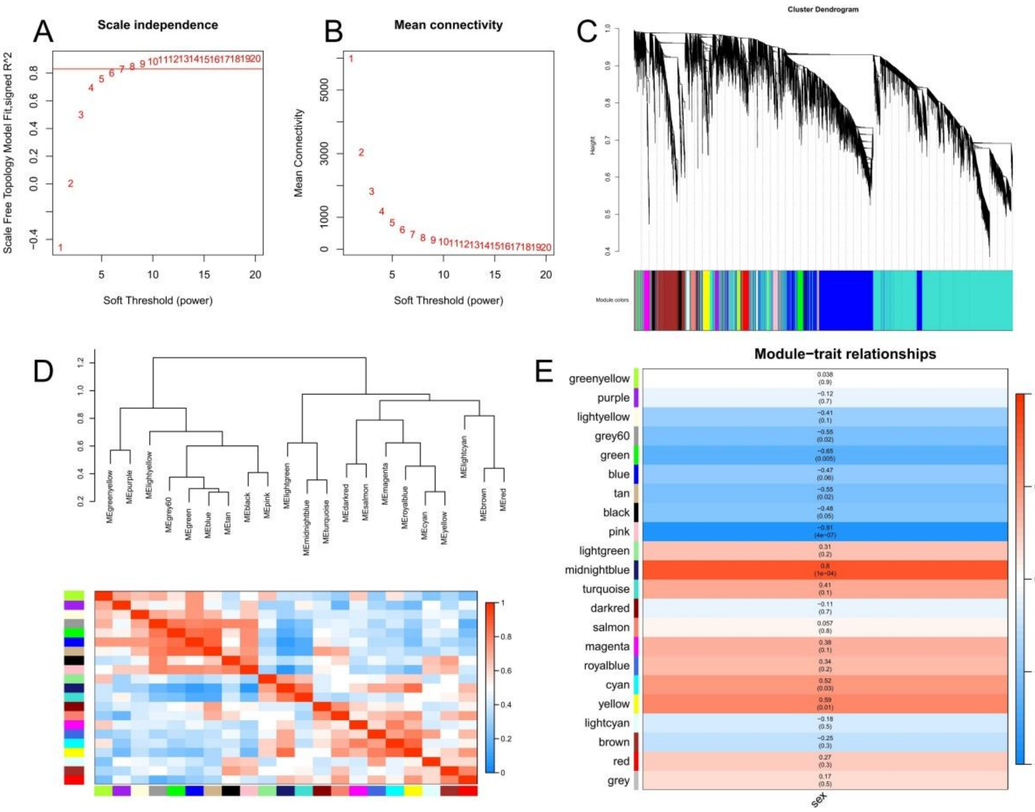


**Fig. S52 Coexpression analysis correlated with sex differentiation between female and male floral buds. (A)** Soft threshold of scale-free topology model fit. **(B)** Soft threshold of mean connectivity. **(C)** Gene cluster dendrogram. **(D)** Eigengene dendrogram. **(E)** Correlation between modules and phenotypes.

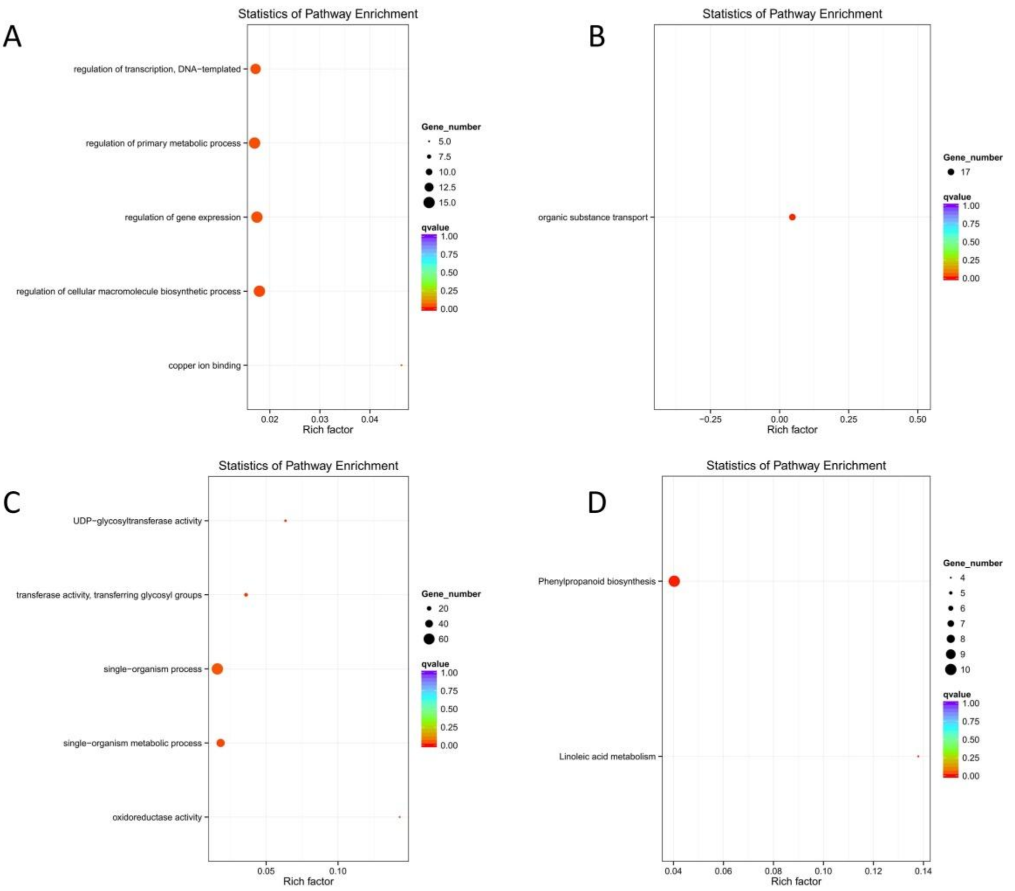


**Fig. S53 Significant GO enrichments in the (A) midnightblue, (B) yellow and (C) pink modules. (D) Significant KEGG pathways in the pink module.**

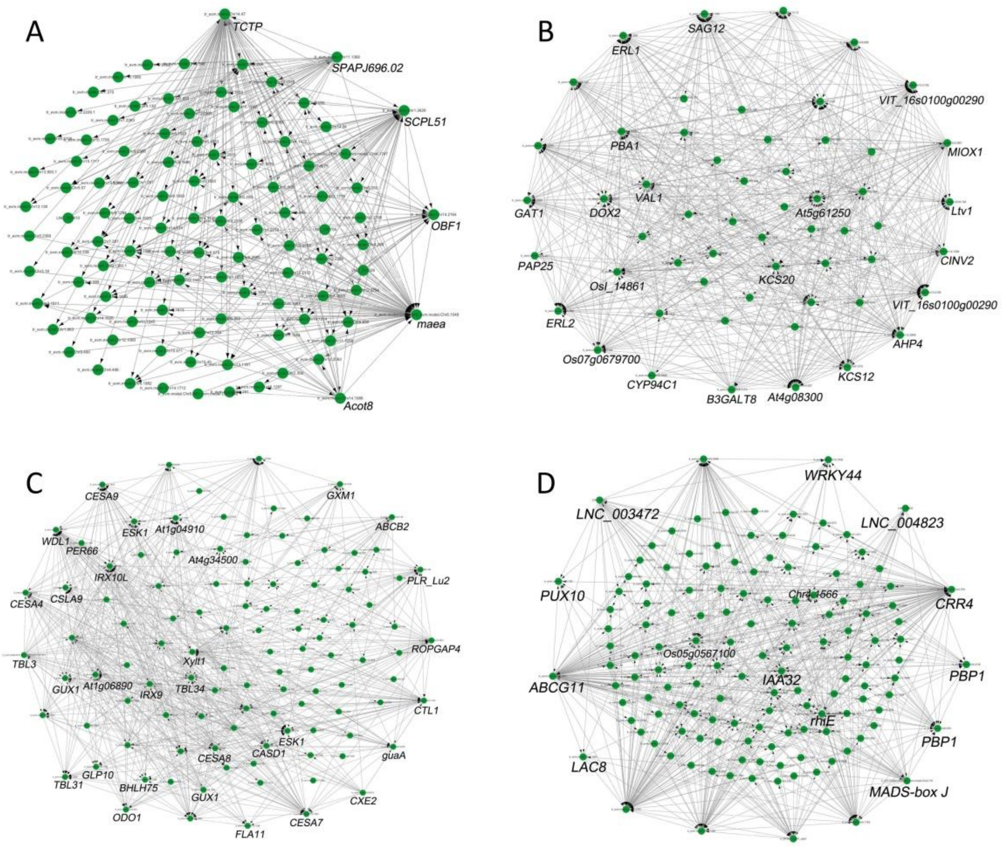


**Fig. S54 Core gene networks in the modules of (A) midnightblue, (B) yellow, (C) pink, and (D) green.**

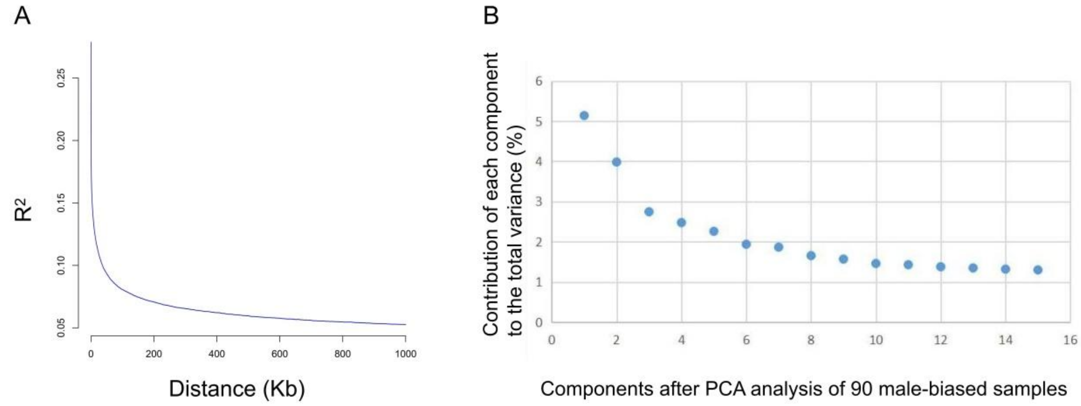


**Fig. S55 (A) LD decay of SNPs obtained from 90 samples before LD pruning. (B) Contribution of each component to the total variance after PCA analysis of 90 genetically male samples.**

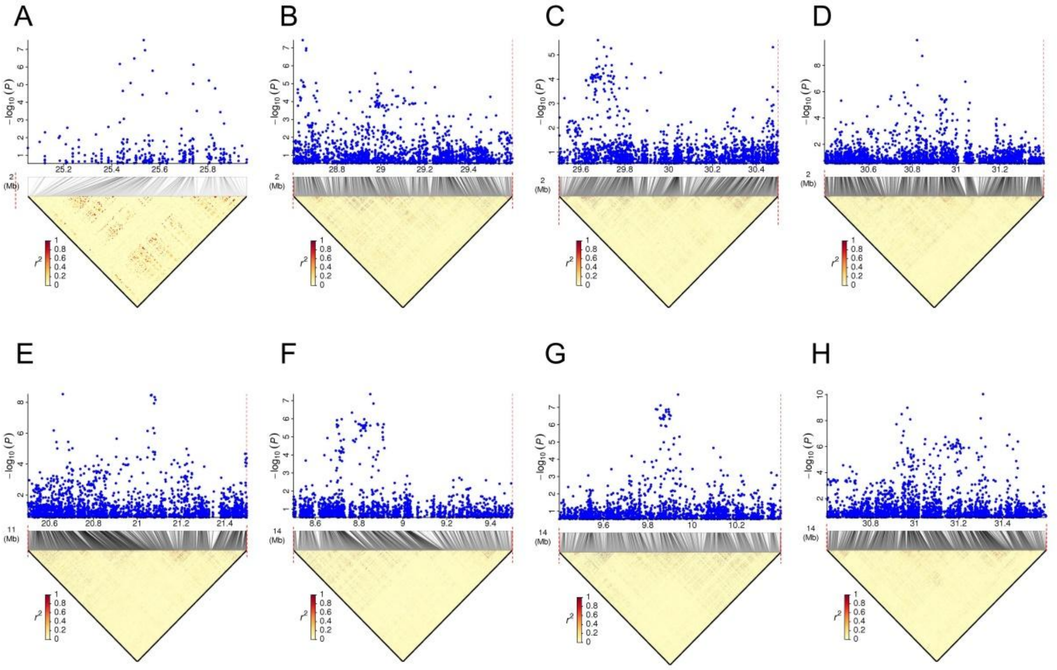


**Fig. S56 Local Manhattan plot (top) and LD heatmap (bottom) within the associated region of proportion of hermaphroditic floral buds. (A)** Chr2: 25.0-26.0. **(B)** Chr2: 28.6-29.6. **(C)** Chr2: 29.5-30.5. **(D)** Chr2: 30.4-31.4. **(E)** Chr11: 20.5-21.5. **(F)** Chr14: 8.5-9.5. **(G)** Chr14: 9.4-10.4. **(H)** Chr14: 30.6-31.6.

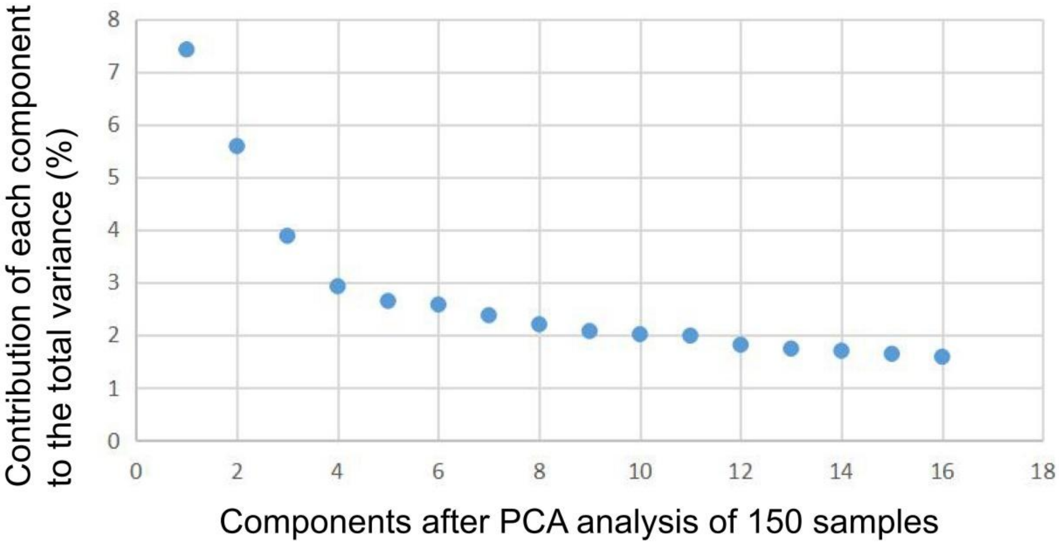


**Fig. S57 Contribution of each component to the total variance after PCA analysis of 150 samples.**

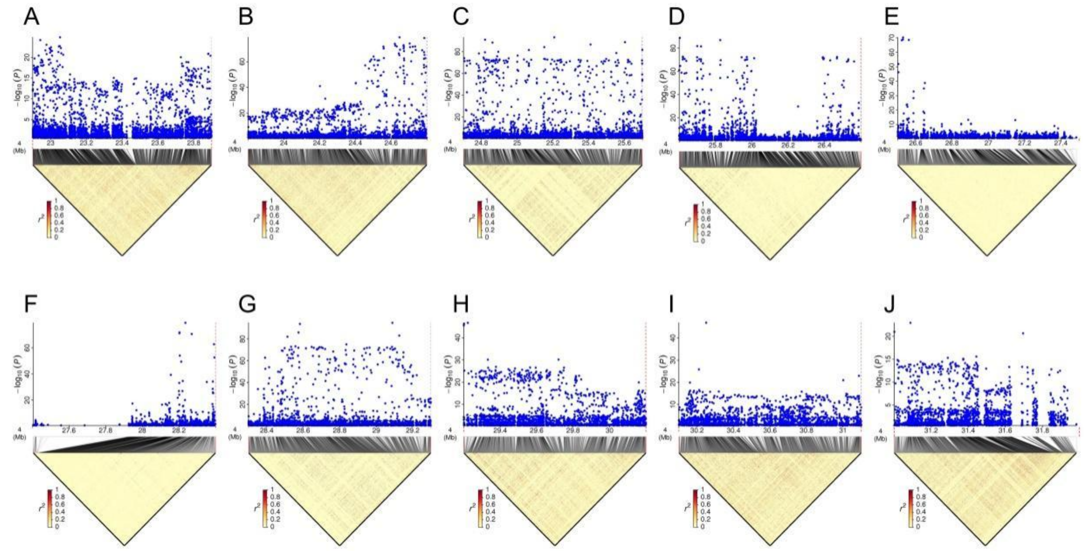


**Fig. S58 Local Manhattan plot (top) and LD heatmap (bottom) in the sex-linked region. (A)** 22.9-23.9 Mb. **(B)** 23.8-24.8 Mb. **(C)** 24.7-25.7 Mb. **(D)** 25.6-26.6 Mb. **(E)** 26.5-27.5 Mb. **(F)** 27.4-28.4 Mb. **(G)** 28.3-29.3. **(H)** 29.2-30.2. **(I)** 30.1-31.1. **(J)** 31.0-32.0.

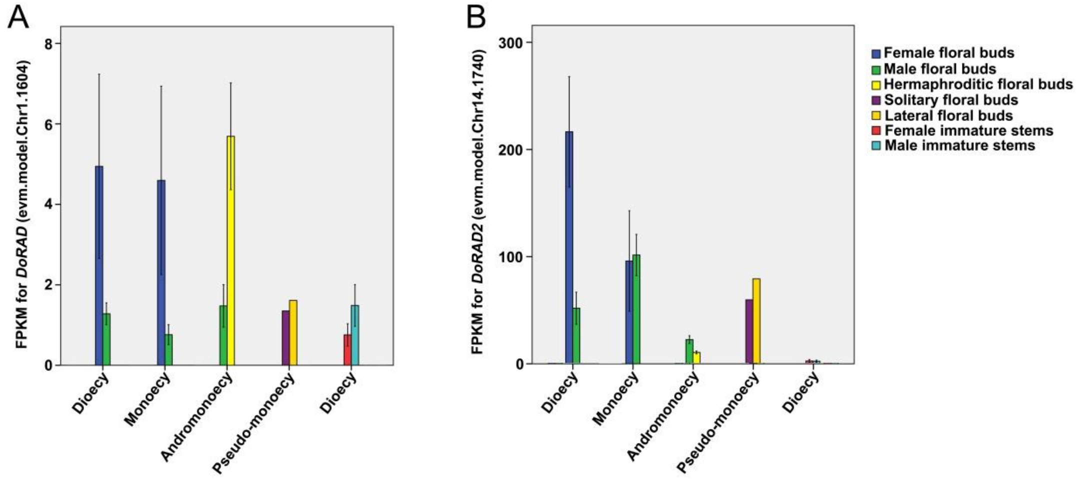


**Fig. S59 FPKM of *DoRAD* and *DoRAD2* in floral buds and stems.** Data are expressed as mean ± standard error (three biological replicates) except for floral buds in pseudo-monoecious trees, for which no biological replicates were available. *DoRAD* and *DoRAD2* represent the genes in *D. oleifeara* homologous to *DkRAD* and *DkRAD2*, respectively. BLAST analysis showed that E-values of DoRAD and DoRAD2 protein against DkRAD and DkRAD2 protein were 3.18E-59 and 3.25E-66, respectively.

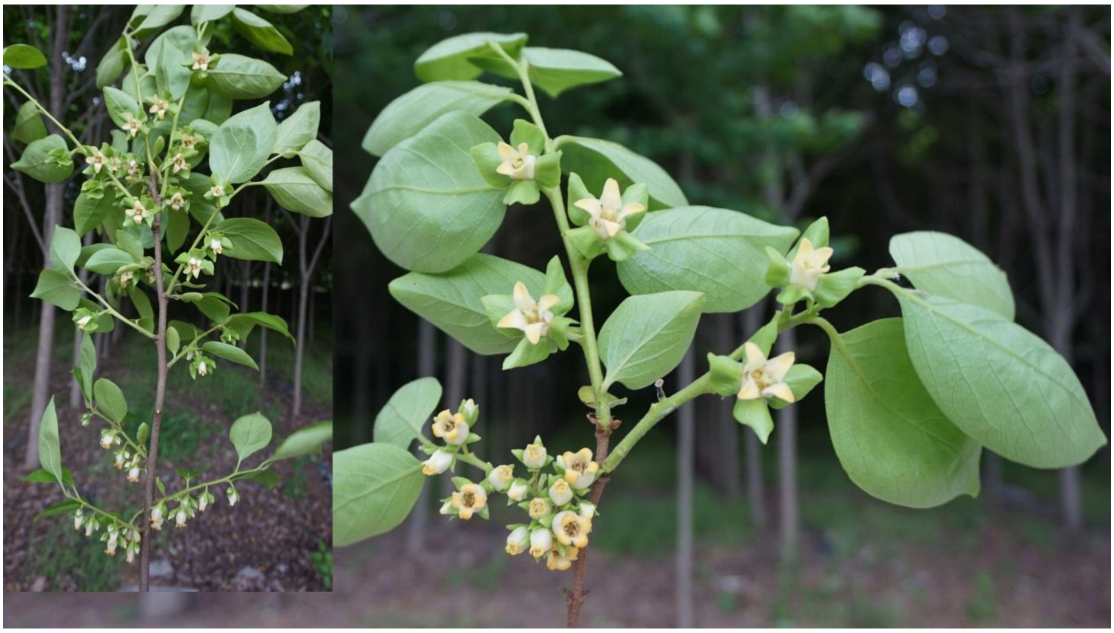

**Fig. S60 Flowering mother branches in the monoecious *D. kaki* cultivar 'Nishimurawase'.** Female shoots are mainly developed from mixed dormant buds that developed on the tips of the fruiting mother branches, while male shoots are mainly developed on the basal parts of the mother branches.
