## Supplementary Tables for "Molecular and genetic mechanisms conferring dissolution of dioecy in *Diospyros oleifera* Cheng"

**Table S1 Contig and scaffold information of *D. oleifera* genome**

| | count | N50 count | N90 count | Min length | N95 length | N90 length | N50 Length | Max length | Total length (Mbp) |
|---|---|---|---|---|---|---|---|---|---|
| contig | 104 | 17 | 49 | 25,519 | 2,198,226 | 3,984,629 | 14,943,463 | 31,016,264 | 690 |
| Scaffold, main genome | 73 | 13 | 31 | 25,519 | 3,843,193 | 7,250,931 | 20,820,941 | 47,738,309 | 700 |
| Scaffold, heterozygous genome | 3735 | 108 | 1761 | 1,469 | 37,312 | 44,522 | 1,111,905 | 13,975,704 | 680 |

**Table S2 Completeness Assessment of the *D. oleifera* main and whole genome by BUSCO**

| **Sample** | **TYPE** | **Main genome** | | **Whole genome** | |
|---|---|---|---|---|---|
| | | **Number** | **Percent (%)** | **Number** | **Percent (%)** |
| *D. oleifera* | Complete BUSCOs (C) | 1241 | 86.2 | 1330 | 92.3 |
| | Complete and single-copy BUSCOs (S) | 1192 | 82.8 | 856 | 59.4 |
| | Complete and duplicated BUSCOs (D) | 49 | 3.4 | 474 | 32.9 |
| | Fragmented BUSCOs (F) | 52 | 3.6 | 39 | 2.7 |
| | Missing BUSCOs (M) | 147 | 10.2 | 71 | 5.0 |
| | Total BUSCO groups searched | 1440 | - | 1440 | - |

**Table S3 Classification of repetitive elements in *D. oleifera* main genome**

| #Total repeative elements | | | |
|---|---|---|---|
| **Program** | | **Repeat size (bp)** | **% of genome** |
| TRF | | 33,164,745 | 4.81 |
| Repeatmasker | | 391,409,187 | 56.73 |
| Proteinmask | | 131,267,592 | 19.03 |
| Total | | 409,994,948 | 59.43 |
| #Transposon elements | | | |
| **Type** | | **Transposon element length (bp)** | **% in genome** |
| DNA | | 36,380,245 | 5.27 |
| LINE | | 13,002,874 | 1.88 |
| SINE | | 92,888 | 0.01 |
| LTR | Total | 338,795,555 | 49.11 |
| | Gypsy | 203,251,006 | 29.46 |
| | Copia | 117,977,800 | 17.10 |
| | Other | 39,723,839 | 5.76 |
| Unknown type | | 25,154,431 | 3.65 |
| Total | | 400,907,095 | 58.11 |

**Table S4 Gene annotation of *D. oleifera* main genome via three methods**

| Method | Gene set | Number | Average transcript length(bp) | Average CDS length(bp) | Average exons per gene | Average exon length(bp) | Average intron length(bp) |
|---|---|---|---|---|---|---|---|
| De novo | Augustus | 29,999 | 5002.98 | 962.17 | 3.99 | 241.03 | 1350.58 |
| | GlimmerHMM | 63,073 | 9790.97 | 517.83 | 2.95 | 175.56 | 4756.36 |
| | SNAP | 29,053 | 11579.55 | 516.75 | 3.19 | 161.8 | 5042.87 |
| | Geneid | 56,788 | 2206.66 | 578.2 | 2.86 | 201.86 | 873.46 |
| | Genscan | 37,328 | 11844.31 | 966.06 | 5.27 | 183.25 | 2546.52 |
| Homolog | *Actinidia chinensis* | 47,559 | 2663.23 | 709.91 | 2.37 | 299.47 | 1425.16 |
| | *Arabidopsis thaliana* | 37,432 | 2942.66 | 803.25 | 2.82 | 284.95 | 1176.17 |
| | *Camellia sinensis* | 13,547 | 5087.21 | 1224.25 | 3.74 | 326.95 | 1407.56 |
| | *Daucus carota* | 15,002 | 4638.45 | 1330.11 | 3.98 | 334.26 | 1110.45 |
| | *Diospyros lotus* | 22,244 | 2101.69 | 782.28 | 2.86 | 273.79 | 710.42 |
| | *Primula veris* | 19,755 | 4574.47 | 907.81 | 3.64 | 249.32 | 1388.28 |
| | *Rhododendron delavayi* | 24,760 | 3418.28 | 1179.25 | 3.27 | 360.93 | 987.54 |
| | *Solanum lycopersicum* | 26,649 | 3532.98 | 1113.05 | 3.27 | 340.37 | 1065.99 |
| RNAseq | PASA | 86,338 | 5460.14 | 805.64 | 3.76 | 214.01 | 1683.66 |
| | Cufflinks | 74,626 | 14618.57 | 2161.92 | 6.27 | 344.66 | 2362.55 |
| | EVM | 32,974 | 6039.66 | 916.44 | 4.13 | 222.09 | 1638.68 |
| | Pasa-update | 32,713 | 6023.58 | 920.54 | 4.1 | 224.73 | 1648.16 |
| | Final set | 26,164 | 6958.94 | 1044.05 | 4.66 | 223.83 | 1614.09 |

**Table S5 Functional annotation of *D. oleifera* main genome**

| Database | Annotated Num | Annotated Percent(%) |
|---|---|---|
| NR | 23,796 | 90.95 |
| Swiss-Prot | 19,301 | 73.77 |
| KEGG | 18,058 | 69.02 |
| InterPro | 20,135 | 76.96 |
| Pfam | 18,463 | 70.57 |
| GO | 13,598 | 51.97 |
| Annotated | 23,854 | 91.17 |
| Unannotated | 2,310 | 8.83 |
| Total | 26,164 | - |

**Table S6 Annotation of non-coding RNAs of *D. oleifera* main genome**

| Type | | Copy(w) | Average length(bp) | Total length(bp) | % of genome |
|---|---|---|---|---|---|
| **miRNA** | | 481 | 115.039501 | 55334 | 0.00802 |
| **tRNA** | | 504 | 74.90277778 | 37751 | 0.005472 |
| **rRNA** | rRNA | 937 | 201.4951974 | 188801 | 0.027365 |
| | 18S | 115 | 797.3391304 | 91694 | 0.01329 |
| | 28S | 166 | 141.7650602 | 23533 | 0.003411 |
| | 5.8S | 56 | 154.4642857 | 8650 | 0.001254 |
| | 5S | 600 | 108.2066667 | 64924 | 0.00941 |
| **snRNA** | snRNA | 736 | 112.9904891 | 83161 | 0.012054 |
| | CD-box | 583 | 104.9948542 | 61212 | 0.008872 |
| | HACA-box | 42 | 137.1428571 | 5760 | 0.000835 |
| | splicing | 110 | 145.6363636 | 16020 | 0.002322 |

**Table S7 Assembly of the male-unmapped sequences**

| Sample_ID | length | | number | |
|---|---|---|---|---|
| | **Contig** | **Scaffold** | **Contig** | **Scaffold** |
| **Total** | 43,558,496 | 43,560,811 | 16,363 | 16,323 |
| **Max** | 35,828 | 35,828 | - | - |
| **Number>=2000** | - | - | 5725 | 5725 |
| **N50** | 3802 | 3802 | 2860 | 2860 |
| **N60** | 2645 | 2647 | 4233 | 4233 |
| **N70** | 1874 | 1874 | 6212 | 6212 |
| **N80** | 1434 | 1434 | 8895 | 8895 |
| **N90** | 1174 | 1174 | 12,272 | 12,272 |

**Table S8 Classification of repetitive elements in the male-unmapped sequences**

| #Total repeative elements | | |
|---|---|---|
| **Program** | **Repeat size (bp)** | **% of genome** |
| TRF | 216,108 | 0.24 |
| Repeatmasker | 355,768 | 0.40 |
| Proteinmask | 109,423 | 0.12 |
| Total | 629,946 | 1.45 |
| #Transposon elements | | |
| **Type** | **Transposon element length (bp)** | **% in genome** |
| DNA | 56,551 | 0.13 |
| LINE | 53,429 | 0.12 |
| SINE | 537 | 0.001 |
| LTR | 150,247 | 0.34 |
| Unknown type | 178,724 | 0.41 |
| Total | 432,479 | 0.99 |

19 **Table S9 Gene annotation of the male-unmapped sequences**

| Gene set | | Number | Average gene length (bp) | Average CDS length (bp) | Average exons per gene | Average exon length (bp) | Average intron length (bp) |
|---|---|---|---|---|---|---|---|
| *Denovo* | Geneid | 24,516 | 1235.67 | 941.96 | 2.16 | 253.26 | 436.15 |
| | Augustus | 34,790 | 877.04 | 742.95 | 1.68 | 198.34 | 443.26 |
| | Genscan | 22,670 | 1567.7 | 1157.3 | 3.43 | 168.61 | 337.01 |
| Homolog | | 4448 | 832.29 | 787.01 | 1.59 | 76.23 | 493.74 |
| Final set | | 2952 | 1238.19 | 1108.19 | 2.10 | 117.93 | 527.13 |

20

**Table S10 Functional annotation of the male-unmapped sequence**

| Database | Annotated Num | Annotated Percent(%) |
|---|---|---|
| **NR** | 2727 | 92.38 |
| **Swiss-Prot** | 2673 | 90.55 |
| **KEGG** | 2584 | 87.53 |
| **InterPro** | 2676 | 90.65 |
| **Pfam** | 2539 | 86.01 |
| **GO** | 2167 | 73.41 |
| **Annotated** | 2729 | 92.45 |
| **Total** | 2952 | - |

**Table S11 Annotation of non-coding RNAs of the male-unmapped sequence**

| Type | | Copy(w) | Average length(bp) | Total length(bp) | % of genome |
|---|---|---|---|---|---|
| **miRNA** | | 4 | 79.5 | 318 | 0.00073 |
| **tRNA** | | 147 | 92.32 | 13,571 | 0.031154 |
| **snRNA** | **snRNA** | 27 | 148.15 | 4000 | 0.009183 |
| | **CD-box** | 8 | 120.38 | 963 | 0.002211 |
| | **HACA-box** | 4 | 177.25 | 709 | 0.001628 |
| | **splicing** | 15 | 155.2 | 2328 | 0.005344 |

**Table S12 Genes used for gene family clustering in each species**

| Species | Latin Name | Number of Genes |
|---|---|---|
| Dol | *Diospyros oleifera* | 29,203 |
| Dlo | *Diospyros lotus* | 51,693 |
| Pve | *Primula veris* | 11,378 |
| Rde | *Rhododendron delavayi* | 32,938 |
| Csi | *Camellia sinensis* | 32,740 |
| Ach | *Actinidia chinensis* | 33,044 |
| Dca | *Daucus carota* | 31,707 |
| Cca | *Coffea canephora* | 25,258 |
| Sly | *Solanum lycopersicum* | 32,837 |
| Ath | *Arabidopsis thaliana* | 26,869 |
| Vvi | *Vitis vinifera* | 28,150 |
| Cme | *Cucumis melo* | 29,732 |

**Table S13 *OGI* positive, binary male expressions and sex expressions for 150 DNA sampled plants**

| Samples | Sex expressions | *OGI* positive | Male expression | 90 plants with male flower production | | |
|---|---|---|---|---|---|---|
| | | | | Proportion of female shoots | Proportion of male shoots | Proportion of hermaphroditic shoots |
| 10 | monoecious | 1 | 1 | 0.2 | 0.8 | 0 |
| 11 | gynoecious | 0 | 0 | / | / | / |
| 12 | gynoecious | 0 | 0 | / | / | / |
| 13 | andromonoecious | 1 | 1 | 0 | 0.8 | 0.2 |
| 14 | gynoecious | 0 | 0 | / | / | / |
| 15 | gynoecious | 0 | 0 | / | / | / |
| 16 | gynoecious | 0 | 0 | / | / | / |
| 17 | gynoecious | 0 | 0 | / | / | / |
| 21 | gynoecious | 0 | 0 | / | / | / |
| 24 | androgynomonoecious | 1 | 1 | 0.1 | 0.8 | 0.1 |
| 25 | gynoecious | 0 | 0 | / | / | / |
| 26 | androgynomonoecious | 1 | 1 | 0.1 | 0.8 | 0.1 |
| 28 | monoecious | 1 | 1 | 0.2 | 0.8 | 0 |
| 29 | gynoecious | 0 | 0 | / | / | / |
| 30 | gynoecious | 0 | 0 | / | / | / |
| 31 | monoecious | 1 | 1 | 0.2 | 0.8 | 0 |
| 34 | andromonoecious | 1 | 1 | 0 | 0.8 | 0.2 |
| 35 | gynoecious | 0 | 0 | / | / | / |
| 36 | gynoecious | 0 | 0 | / | / | / |
| 38 | gynoecious | 1 | 0 | / | / | / |
| 39 | gynoecious | 0 | 0 | / | / | / |
| 40 | monoecious | 1 | 1 | 0.2 | 0.8 | 0 |
| 44 | monoecious | 1 | 1 | 0.2 | 0.8 | 0 |
| 45 | androecious | 1 | 1 | 0 | 1 | 0 |
| 47 | monoecious | 1 | 1 | 0.5 | 0.5 | 0 |
| 49 | gynoecious | 0 | 0 | / | / | / |
| 4 | gynoecious | 0 | 0 | / | / | / |
| 50 | androgynomonoecious | 1 | 1 | 0.1 | 0.8 | 0.1 |

| | | | | | | |
|---|---|---|---|---|---|---|
| 5 | pseudo-monoecious | 0 | 0 | / | / | / |
| 60 | gynoecious | 0 | 0 | / | / | / |
| 6 | pseudo-monoecious | 0 | 0 | / | / | / |
| 75 | gynoecious | 0 | 0 | / | / | / |
| 83 | andromonoecious | 1 | 1 | 0 | 0.9 | 0.1 |
| 87 | monoecious | 1 | 1 | 0.5 | 0.5 | 0 |
| 8 | monoecious | 1 | 1 | 0.2 | 0.8 | 0 |
| 9 | gynoecious | 0 | 0 | / | / | / |
| 100 | gynoecious | 0 | 0 | / | / | / |
| 108 | monoecious | 1 | 1 | 0.2 | 0.8 | 0 |
| 109 | monoecious | 1 | 1 | 0.5 | 0.5 | 0 |
| 111 | monoecious | 1 | 1 | 0.2 | 0.8 | 0 |
| 114 | gynoecious | 0 | 0 | / | / | / |
| 116 | gynoecious | 0 | 0 | / | / | / |
| 118 | monoecious | 1 | 1 | 0.5 | 0.5 | 0 |
| 119 | gynoecious | 0 | 0 | / | / | / |
| 121 | monoecious | 1 | 1 | 0.5 | 0.5 | 0 |
| 122 | monoecious | 1 | 1 | 0.5 | 0.5 | 0 |
| 124 | gynoecious | 0 | 0 | / | / | / |
| 125 | androecious | 1 | 1 | 0 | 1 | 0 |
| 126 | gynoecious | 0 | 0 | / | / | / |
| 127 | androecious | 1 | 1 | 0 | 1 | 0 |
| 133 | androecious | 1 | 1 | 0 | 1 | 0 |
| 136 | androgynomonoecious | 1 | 1 | 0.4 | 0.4 | 0.2 |
| 52 | monoecious | 1 | 1 | 0.2 | 0.8 | 0 |
| 53 | monoecious | 1 | 1 | 0.2 | 0.8 | 0 |
| 56 | androecious | 1 | 1 | 0 | 1 | 0 |
| 57 | androecious | 1 | 1 | 0 | 1 | 0 |
| 58 | gynoecious | 0 | 0 | / | / | / |
| 59 | androecious | 1 | 1 | 0 | 1 | 0 |
| 65 | androecious | 1 | 1 | 0 | 1 | 0 |
| 68 | androecious | 1 | 1 | 0 | 1 | 0 |
| 69 | androecious | 1 | 1 | 0 | 1 | 0 |
| 70 | androecious | 1 | 1 | 0 | 1 | 0 |

| | | | | | | |
|---|---|---|---|---|---|---|
| 73 | gynoecious | 0 | 0 | / | / | / |
| 74 | monoecious | 1 | 1 | 0.2 | 0.8 | 0 |
| 78 | gynoecious | 0 | 0 | / | / | / |
| 90 | monoecious | 1 | 1 | 0.2 | 0.8 | 0 |
| 93 | androecious | 1 | 1 | 0 | 1 | 0 |
| 95 | androecious | 1 | 1 | 0 | 1 | 0 |
| 96 | androecious | 1 | 1 | 0 | 1 | 0 |
| 97 | androecious | 1 | 1 | 0 | 1 | 0 |
| 98 | androecious | 1 | 1 | 0 | 1 | 0 |
| 99 | androecious | 1 | 1 | 0 | 1 | 0 |
| 135 | monoecious | 1 | 1 | 0.5 | 0.5 | 0 |
| 138 | gynoecious | 0 | 0 | / | / | / |
| 141 | gynoecious | 0 | 0 | / | / | / |
| 142 | androecious | 1 | 1 | 0 | 1 | 0 |
| 144 | gynoecious | 0 | 0 | / | / | 0 |
| 145 | androecious | 1 | 1 | 0 | 1 | 0 |
| 146 | androecious | 1 | 1 | 0 | 1 | 0 |
| 149 | androecious | 1 | 1 | 0 | 1 | 0 |
| 155 | androecious | 1 | 1 | 0 | 1 | 0 |
| 157 | androecious | 1 | 1 | 0 | 1 | 0 |
| 158 | gynoecious | 0 | 0 | / | / | / |
| 159 | androecious | 1 | 1 | 0 | 1 | 0 |
| 161 | androecious | 1 | 1 | 0 | 1 | 0 |
| 162 | androecious | 1 | 1 | 0 | 1 | 0 |
| 167 | andromonoecious | 1 | 1 | 0 | 0.8 | 0.2 |
| 168 | androgynomonoecious | 1 | 1 | 0.4 | 0.4 | 0.2 |
| 173 | androgynomonoecious | 1 | 1 | 0.4 | 0.4 | 0.2 |
| 174 | androecious | 1 | 1 | 0 | 1 | 0 |
| 175 | gynoecious | 1 | 0 | / | / | / |
| 176 | gynoecious | 0 | 0 | / | / | / |
| 177 | androecious | 1 | 1 | 0 | 1 | 0 |
| 179 | monoecious | 0 | 1 | 0.7 | 0.3 | 0 |
| 180 | monoecious | 1 | 1 | 0.6 | 0.4 | 0 |
| 185 | androecious | 1 | 1 | 0 | 1 | 0 |

| | | | | | | |
|---|---|---|---|---|---|---|
| 186 | gynoecious | 0 | 0 | / | / | / |
| 188 | androecious | 1 | 1 | 0 | 1 | 0 |
| 192 | gynoecious | 0 | 0 | / | / | / |
| 193 | androecious | 1 | 1 | 0 | 1 | 0 |
| 196 | gynoecious | 0 | 0 | / | / | / |
| 198 | gynoecious | 0 | 0 | / | / | / |
| 200 | gynoecious | 0 | 0 | / | / | / |
| 201 | androecious | 1 | 1 | 0 | 1 | 0 |
| 206 | andromonoecious | 1 | 1 | 0 | 0.66 | 0.33 |
| 208 | androecious | 1 | 1 | 0 | 1 | 0 |
| 209 | gynoecious | 0 | 0 | / | / | / |
| 211 | androgynomonoecious | 1 | 1 | 0.3 | 0.5 | 0.2 |
| 102 | gynoecious | 0 | 0 | / | / | / |
| 106 | gynoecious | 0 | 0 | / | / | / |
| 113 | androecious | 1 | 1 | 0 | 1 | 0 |
| 128 | androecious | 1 | 1 | 0 | 1 | 0 |
| 130 | andromonoecious | 1 | 1 | 0 | 0.7 | 0.3 |
| 139 | androecious | 1 | 1 | 0 | 1 | 0 |
| 147 | gynoecious | 0 | 0 | / | / | / |
| 153 | gynoecious | 0 | 0 | / | / | / |
| 154 | androecious | 1 | 1 | 0 | 1 | 0 |
| 163 | gynoecious | 0 | 0 | / | / | / |
| 164 | androecious | 1 | 1 | 0 | 1 | 0 |
| 169 | androecious | 1 | 1 | 0 | 1 | 0 |
| 170 | androecious | 1 | 1 | 0 | 1 | 0 |
| 182 | monoecious | 1 | 1 | 0.05 | 0.95 | 0 |
| 183 | gynoecious | 0 | 0 | / | / | / |
| 187 | androecious | 1 | 1 | 0 | 1 | 0 |
| 189 | monoecious | 1 | 1 | 0.02 | 0.98 | 0 |
| 18 | gynoecious | 0 | 0 | / | / | / |
| 19 | monoecious | 1 | 1 | 0.1 | 0.9 | 0 |
| 204 | androecious | 1 | 1 | 0 | 1 | 0 |
| 210 | androgynomonoecious | 1 | 1 | 0.1 | 0.8 | 0.1 |
| 41 | monoecious | 1 | 1 | 0.2 | 0.8 | 0 |

| | | | | | | |
|---|---|---|---|---|---|---|
| 62 | gynoecious | 1 | 0 | / | / | / |
| 80 | gynoecious | 1 | 0 | / | / | / |
| 82 | gynoecious | 0 | 0 | / | / | / |
| 86 | gynoecious | 0 | 0 | / | / | / |
| 89 | gynoecious | 0 | 0 | / | / | / |
| 107 | androgynomonoecious | 1 | 1 | 0.1 | 0.75 | 0.15 |
| 110 | gynoecious | 0 | 0 | / | / | / |
| 131 | gynoecious | 0 | 0 | / | / | / |
| 137 | androgynomonoecious | 1 | 1 | 0.1 | 0.8 | 0.1 |
| 150 | androecious | 1 | 1 | 0 | 1 | 0 |
| 151 | gynoecious | 0 | 0 | / | / | / |
| 166 | andromonoecious | 1 | 1 | 0 | 0.7 | 0.3 |
| 184 | androecious | 1 | 1 | 0 | 1 | 0 |
| 205 | gynoecious | 0 | 0 | / | / | / |
| 207 | androecious | 1 | 1 | 0 | 1 | 0 |
| 22 | gynoecious | 0 | 0 | / | / | / |
| 42 | monoecious | 1 | 1 | 0.1 | 0.9 | 0 |
| 64 | gynoecious | 0 | 0 | / | / | / |
| 67 | monoecious | 1 | 1 | 0.05 | 0.95 | 0 |
| 84 | monoecious | 1 | 1 | 0.3 | 0.7 | 0 |

**Table S14 Classification of samples used for methylome detection**

| Sex phenotype of a tree | Groups | Samples within each group |
|---|---|---|
| Gynoecious ($OGI^-$) | G_F (gynoecious_female floral buds) | 11F; 21F; 186F |
| | G_S (gynoecious_stems of immature flowering shoots) | 11S; 21S; 186S |
| Androecious ($OGI^+$) | A_M (androecious_male floral buds) | 13M; 65M; 188M |
| | A_S (androecious_stems of immature flowering shoots) | 13S; 65S; 188S |
| Monoecious ($OGI^+$) | M_F (monoecious_female floral buds) | 108F; 168F |
| | M_M (monoecious_male floral buds) | 108M; 168M |
| | M_S (monoecious_stems of immature flowering shoots) | 108S; 168S |
| Andromonoecious ($OGI^+$) | AM_M (andromonoecious_male floral buds) | 206M with three biological replicates |
| | AM_H (andromonoecious_hermaphroditic floral buds) | 206H with two biological replicates |
| | AM_L (andromonoecy_leaf) | 206L with three biological replicates |

| | | |
|---|---|---|
| Pseudo-monoecious (*$OGI^{-}$*) | PM_SF (pseudo-monoecy_solitary female floral buds) | PM_SF |
| | PM_MF (pseudo-monoecy_middle floral buds obtained from the three flower cymes) | PM_MF |
| | PM_LF (pseudo-monoecy_lateral floral buds obtained from the three flower cymes) | PM_LF |
| | PM_S (pseudo-monoecy_stems of immature flowering shoots) | PM_S |

**Table S15 Classification of samples used for whole transcriptome analysis**

| Sex phenotype of a tree | Groups | Samples within each group |
|---|---|---|
| Gynoecious (*$OGI^{-}$*) | G_F (gynoecy_female floral buds) | 11F; 21F; 186F |
| | G_S (gynoecy_stems of immature flowering shoots) | 11S; 21S; 186S |
| Androecious (*$OGI^{+}$*) | A_M (androecy_male floral buds) | 13M; 65M; 188M |
| | A_S (androecy_stems of immature flowering shoots) | 13S; 65S; 188S |
| Monoecious (*$OGI^{+}$*) | M_F (monoecy_female floral buds) | 108F; 136F; 168F |
| | M_M (monoecy_male floral buds) | 108M; 136M; 168M |
| Andromonoecious (*$OGI^{+}$*) | AM_M (andromonoecy_male floral buds) | 206M with three biological replicates |
| | AM_H (andromonoecy_hermaphroditic floral buds) | 206H with three biological replicates |
| Pseudo-monoecious (*$OGI^{-}$*) | PM_SF (pseudo-monoecy_solitary female floral buds) | PM_SF |
| | PM_LF (pseudo monoecy_lateral floral buds obtained from the three flower cymes) | PM_LF |

**Table S16 Statistics of bisulphite sequencing**

| Sample_name | Raw_reads | Raw_base (G) | Clean_reads | Clean_base (G) | Clean_ratio (%) | Q20 (%) | Q30 (%) | GC (%) |
|---|---|---|---|---|---|---|---|---|
| **A13_M** | 92148953 | 27.64 | 89826370 | 24.59 | 88.97 | 96.44 | 89.64 | 22.29 |
| **A13_S** | 93644119 | 28.09 | 92346348 | 25.43 | 90.53 | 97.92 | 93.15 | 27.02 |
| **A188_M** | 103114616 | 30.93 | 100239487 | 27.34 | 88.39 | 97.27 | 91.29 | 22.44 |
| **A188_S** | 90120186 | 27.04 | 88982849 | 24.52 | 90.68 | 97.91 | 93.07 | 23.15 |
| **A65_M** | 96373495 | 28.91 | 95224820 | 26.27 | 90.87 | 97.99 | 93.27 | 24.42 |
| **A65_S** | 112821125 | 33.85 | 109863235 | 30.01 | 88.66 | 97.4 | 91.59 | 22.26 |
| **AM206_H1** | 93454597 | 28.04 | 92224569 | 25.4 | 90.58 | 97.87 | 92.98 | 23.87 |
| **AM206_H2** | 112045237 | 33.61 | 109154015 | 29.8 | 88.66 | 97.38 | 91.53 | 22.15 |
| **AM206_M1** | 112595597 | 33.78 | 110996807 | 30.55 | 90.44 | 97.71 | 92.58 | 23.57 |
| **AM206_M2** | 95875729 | 28.76 | 94681568 | 26.09 | 90.72 | 97.93 | 93.11 | 23.73 |
| **AM206_M3** | 91378618 | 27.41 | 90254213 | 24.89 | 90.81 | 97.99 | 93.26 | 24.05 |
| **AM206_S1** | 107696962 | 32.31 | 106250592 | 29.26 | 90.56 | 97.77 | 92.7 | 22.49 |
| **AM206_S2** | 120835381 | 36.25 | 119325363 | 32.89 | 90.73 | 97.93 | 93.09 | 23.15 |
| **AM206_S3** | 95859229 | 28.76 | 94569883 | 26.05 | 90.58 | 97.8 | 92.77 | 23.14 |
| **G11_F** | 94881996 | 28.46 | 92165053 | 25.13 | 88.3 | 97.22 | 91.18 | 22.14 |
| **G11_S** | 101344475 | 30.4 | 100018248 | 27.54 | 90.59 | 97.82 | 92.85 | 23.42 |
| **G186_F** | 93125674 | 27.94 | 90694141 | 24.81 | 88.8 | 96.31 | 89.39 | 22.27 |
| **G186_S** | 90352254 | 27.11 | 88311166 | 24.18 | 89.19 | 96.56 | 89.88 | 22.45 |
| **G21_F** | 109002759 | 32.7 | 105846010 | 28.85 | 88.23 | 97.17 | 91.06 | 22.52 |
| **G21_S** | 96279608 | 28.88 | 95116684 | 26.2 | 90.72 | 97.95 | 93.18 | 24.03 |
| **M108_F** | 98003581 | 29.4 | 96719011 | 26.64 | 90.61 | 97.8 | 92.78 | 23.4 |
| **M108_M** | 90732540 | 27.22 | 87861494 | 23.92 | 87.88 | 97.12 | 90.99 | 22.48 |
| **M108_S** | 113236451 | 33.97 | 111759050 | 30.79 | 90.64 | 97.84 | 92.89 | 24.47 |
| **M168_F** | 93130691 | 27.94 | 91054869 | 24.97 | 89.37 | 96.73 | 90.2 | 22.37 |
| **M168_M** | 93114357 | 27.93 | 91838635 | 25.29 | 90.55 | 97.74 | 92.65 | 24.21 |
| **M168_S** | 95166720 | 28.55 | 93831524 | 25.83 | 90.47 | 97.84 | 92.94 | 25.44 |
| **M5_F** | 116876643 | 35.06 | 113657234 | 31 | 88.42 | 97.29 | 91.33 | 22.3 |
| **M5_H** | 95238839 | 28.57 | 92837301 | 25.42 | 88.97 | 96.41 | 89.57 | 22.27 |
| **M5_M** | 104829097 | 31.45 | 102199093 | 27.88 | 88.65 | 97.26 | 91.23 | 22.34 |
| **M5_S** | 88724875 | 26.62 | 86324616 | 23.6 | 88.66 | 96.23 | 89.22 | 22.2 |
| **Average** | 99,733,480.13 | 29.92 | 97,805,808.27 | 26.84 | 89.71 | 97.42 | 91.85 | 23.20 |
| **Total** | 2,992,004,404 | 897.58 | 2,934,174,248 | 805.14 | / | / | / | / |

**Table S17 Bisulphite conversion rate**

| Sample | C_Methylation (%) | CpG_methylation (%) | CHG_methylation (%) | CHH_methylation (%) |
|---|---|---|---|---|
| **A13_M** | 99.4911 | 99.5258 | 99.5052 | 99.4643 |
| **A13_S** | 99.9121 | 99.8977 | 99.9115 | 99.9216 |
| **A188_M** | 99.4851 | 99.5191 | 99.4992 | 99.4584 |
| **A188_S** | 99.8587 | 99.8324 | 99.8598 | 99.8711 |
| **A65_M** | 99.8785 | 99.8589 | 99.8719 | 99.8907 |
| **A65_S** | 99.504 | 99.5377 | 99.5184 | 99.4781 |
| **AM206_H1** | 99.8583 | 99.8425 | 99.8499 | 99.8704 |
| **AM206_H2** | 99.6016 | 99.6335 | 99.618 | 99.5748 |
| **AM206_M1** | 99.8557 | 99.8362 | 99.8475 | 99.869 |
| **AM206_M2** | 99.851 | 99.8305 | 99.8496 | 99.8621 |
| **AM206_M3** | 99.8538 | 99.8373 | 99.8482 | 99.8647 |
| **AM206_S1** | 99.8395 | 99.8231 | 99.8355 | 99.85 |
| **AM206_S2** | 99.8533 | 99.8381 | 99.8433 | 99.8663 |
| **AM206_S3** | 99.8492 | 99.8311 | 99.8441 | 99.8612 |
| **G11_F** | 99.5982 | 99.63 | 99.6101 | 99.5743 |
| **G11_S** | 99.8575 | 99.8354 | 99.8522 | 99.8715 |
| **G186_F** | 99.5073 | 99.5363 | 99.5214 | 99.4836 |
| **G186_S** | 99.5257 | 99.5601 | 99.5387 | 99.4999 |
| **G21_F** | 99.5929 | 99.6216 | 99.6076 | 99.5687 |
| **G21_S** | 99.8477 | 99.8259 | 99.8402 | 99.8628 |
| **M108_F** | 99.8613 | 99.8407 | 99.8578 | 99.8734 |
| **M108_M** | 99.5553 | 99.5848 | 99.5687 | 99.5315 |
| **M108_S** | 99.897 | 99.873 | 99.8906 | 99.9129 |
| **M168_F** | 99.5376 | 99.5687 | 99.5514 | 99.5128 |
| **M168_M** | 99.862 | 99.8439 | 99.8569 | 99.873 |
| **M168_S** | 99.8646 | 99.8425 | 99.8589 | 99.8795 |
| **M5_F** | 99.5287 | 99.5595 | 99.5446 | 99.5027 |
| **M5_H** | 99.5495 | 99.5821 | 99.563 | 99.5247 |
| **M5_M** | 99.5479 | 99.5774 | 99.5647 | 99.5219 |
| **M5_S** | 99.5383 | 99.5682 | 99.5532 | 99.5148 |
| **Average** | 99.71 | 99.72 | 99.72 | 99.71 |

**Table S18 Mapping rate of bisulphite sequencing reads to the genome**

| Samples | Total reads | Mapped reads | Mapping rate(%) | Duplication rate(%) |
|---|---|---|---|---|
| **A13_M** | 89,826,370 | 57,300,241 | 63.79 | 14.39 |
| **A13_S** | 92,346,348 | 53,256,138 | 57.67 | 36.56 |
| **A188_M** | 100,239,487 | 65,356,145 | 65.2 | 16.82 |
| **A188_S** | 88,982,849 | 54,849,028 | 61.64 | 27.61 |
| **A65_M** | 95,224,820 | 58,696,579 | 61.64 | 29.93 |
| **A65_S** | 109,863,235 | 68,785,371 | 62.61 | 17.69 |
| **AM206_H1** | 92,224,569 | 55,980,313 | 60.7 | 23.5 |
| **AM206_H2** | 109,154,015 | 71,495,879 | 65.5 | 16.82 |
| **AM206_M1** | 110,996,807 | 68,673,724 | 61.87 | 23.31 |
| **AM206_M2** | 94,681,568 | 58,077,673 | 61.34 | 24.37 |
| **AM206_M3** | 90,254,213 | 55,948,586 | 61.99 | 22.08 |
| **AM206_S1** | 106,250,592 | 68,287,255 | 64.27 | 24.02 |
| **AM206_S2** | 119,325,363 | 76,618,815 | 64.21 | 25.03 |
| **AM206_S3** | 94,569,883 | 60,600,381 | 64.08 | 22.77 |
| **G11_F** | 92,165,053 | 61,234,461 | 66.44 | 17.89 |
| **G11_S** | 100,018,248 | 60,611,058 | 60.6 | 30.87 |
| **G186_F** | 90,694,141 | 59,649,536 | 65.77 | 14.27 |
| **G186_S** | 88,311,166 | 56,890,053 | 64.42 | 13.68 |
| **G21_F** | 105,846,010 | 67,847,292 | 64.1 | 16.14 |
| **G21_S** | 95,116,684 | 56,299,565 | 59.19 | 25.84 |

| | | | | |
|---|---|---|---|---|
| **M108_F** | 96,719,011 | 58,766,471 | 60.76 | 24.83 |
| **M108_M** | 87,861,494 | 57,057,254 | 64.94 | 15.98 |
| **M108_S** | 111,759,050 | 68,508,297 | 61.3 | 26.91 |
| **M168_F** | 91,054,869 | 59,677,361 | 65.54 | 14.58 |
| **M168_M** | 91,838,635 | 57,940,994 | 63.09 | 24.02 |
| **M168_S** | 93,831,524 | 57,828,368 | 61.63 | 31.99 |
| **M5_F** | 113,657,234 | 73,945,396 | 65.06 | 18.92 |
| **M5_H** | 92,837,301 | 58,747,444 | 63.28 | 14.41 |
| **M5_M** | 102,199,093 | 66,899,526 | 65.46 | 16.54 |
| **M5_S** | 86,324,616 | 53,970,149 | 62.52 | 13.69 |
| **Average** | 97,805,808.27 | 61,659,978.43 | 63.02 | 21.52 |

**Table S19 Coverage depth of bisulphite sequencing reads to the genome**

| **Samples** | **sites_num** | **sites_covg Mean** | **sites_numC ovg1 (%)** | **sites_numC ovg5 (%)** | **sites_numC ovg10 (%)** |
|---|---|---|---|---|---|
| **A13_M** | 619,226,585 | 18.35 | 84.22 | 75.07 | 66.67 |
| **A13_S** | 560,678,937 | 12.68 | 76.25 | 49.06 | 25.41 |
| **A188_M** | 628,793,900 | 20.26 | 85.52 | 77.21 | 69.89 |
| **A188_S** | 588,516,665 | 14.92 | 80.04 | 55.19 | 32.4 |
| **A65_M** | 553,905,231 | 15.47 | 75.33 | 48.01 | 31.38 |
| **A65_S** | 619,141,249 | 21.13 | 84.2 | 75.5 | 68.59 |
| **AM206_H1** | 591,566,693 | 16.1 | 80.45 | 58.44 | 38.22 |
| **AM206_H2** | 626,304,725 | 22.18 | 85.18 | 77.84 | 71.39 |
| **AM206_M1** | 603,727,198 | 19.79 | 82.11 | 64.01 | 45.72 |
| **AM206_M2** | 591,624,871 | 16.51 | 80.46 | 56.79 | 35.55 |
| **AM206_M3** | 593,837,658 | 16.4 | 80.76 | 60.01 | 40.22 |
| **AM206_S1** | 605,902,806 | 19.48 | 82.4 | 67.36 | 45.36 |
| **AM206_S2** | 608,301,912 | 21.58 | 82.73 | 69 | 49.33 |
| **AM206_S3** | 600,758,631 | 17.58 | 81.7 | 63.54 | 41.18 |
| **G11_F** | 623,422,972 | 18.73 | 84.79 | 76.18 | 68.14 |
| **G11_S** | 595,848,844 | 15.73 | 81.04 | 62.61 | 36.89 |
| **G186_F** | 628,380,199 | 19.12 | 85.46 | 76.89 | 68.95 |
| **G186_S** | 625,896,923 | 18.37 | 85.12 | 76.45 | 67.9 |
| **G21_F** | 618,159,710 | 21.19 | 84.07 | 75.67 | 68.68 |
| **G21_S** | 586,741,318 | 15.69 | 79.8 | 60.7 | 35.88 |
| **M108_F** | 601,017,566 | 16.6 | 81.74 | 62.08 | 37.47 |
| **M108_M** | 626,443,576 | 17.84 | 85.2 | 76.76 | 67.75 |
| **M108_S** | 603,915,115 | 18.81 | 82.13 | 65.49 | 42.6 |
| **M168_F** | 625,528,178 | 19.09 | 85.07 | 76.73 | 68.74 |
| **M168_M** | 597,317,872 | 16.55 | 81.24 | 60.8 | 41.75 |
| **M168_S** | 588,141,213 | 14.76 | 79.99 | 55.87 | 29.94 |
| **M5_F** | 629,128,853 | 22.34 | 85.56 | 77.26 | 70.59 |
| **M5_H** | 624,064,207 | 18.81 | 84.87 | 75.83 | 67.36 |
| **M5_M** | 625,132,820 | 20.8 | 85.02 | 76.79 | 69.53 |
| **M5_S** | 620,542,040 | 17.41 | 84.4 | 75.17 | 65.66 |
| **Average** | 607,065,615.6 | 18.14 | 82.56 | 67.61 | 52.30 |

**Note: "sites_num" represents the number of loci detected on the genome; "sites_covgMean" represents average coverage depth of all loci on the genome; "sites_numCovg1 (%)", "sites_numCovg5 (%)", and "sites_numCovg10 (%)" represent the proportion of bases with greater than or equal to 1×, 5×, and 10× coverage depth to the total genome length, respectively.**

**Table S20 Percentage of methylated cytocines**

| **Samples** | **mC percent (%)** | **mCpG percent (%)** | **mCHG percent (%)** | **mCHH percent (%)** |
|---|---|---|---|---|
| **A13_M** | 2.48 | 4.07 | 5.19 | 1.74 |

| | | | | |
|---|---|---|---|---|
| **A13_S** | 1.41 | 1.28 | 1.43 | 1.43 |
| **A188_M** | 3.08 | 4.63 | 5.8 | 2.35 |
| **A188_S** | 1.37 | 1.44 | 1.79 | 1.29 |
| **A65_M** | 1.72 | 1.52 | 2.49 | 1.62 |
| **A65_S** | 3.36 | 4.2 | 5.77 | 2.8 |
| **AM206_H1** | 1.77 | 1.64 | 2.18 | 1.72 |
| **AM206_H2** | 3.26 | 4.54 | 6.09 | 2.55 |
| **AM206_M1** | 2.37 | 2.11 | 3.39 | 2.23 |
| **AM206_M2** | 1.85 | 1.63 | 2.2 | 1.83 |
| **AM206_M3** | 1.84 | 1.71 | 2.3 | 1.78 |
| **AM206_S1** | 1.59 | 2 | 2.64 | 1.34 |
| **AM206_S2** | 1.85 | 2.2 | 3.28 | 1.54 |
| **AM206_S3** | 1.52 | 1.8 | 2.57 | 1.29 |
| **G11_F** | 2.42 | 3.67 | 4.8 | 1.79 |
| **G11_S** | 1.45 | 1.39 | 1.58 | 1.43 |
| **G186_F** | 2.66 | 4.32 | 5.42 | 1.91 |
| **G186_S** | 3.06 | 4.1 | 5.12 | 2.53 |
| **G21_F** | 2.97 | 4.2 | 6.04 | 2.23 |
| **G21_S** | 1.71 | 1.43 | 1.75 | 1.75 |
| **M108_F** | 1.79 | 1.62 | 2.14 | 1.76 |
| **M108_M** | 2.63 | 4 | 4.91 | 2 |
| **M108_S** | 2.22 | 1.86 | 2.46 | 2.23 |
| **M168_F** | 2.6 | 3.95 | 5.09 | 1.93 |
| **M168_M** | 1.92 | 1.83 | 2.55 | 1.82 |
| **M168_S** | 1.66 | 1.35 | 1.62 | 1.72 |
| **M5_F** | 2.99 | 4.77 | 6.4 | 2.1 |
| **M5_H** | 2.46 | 4.08 | 4.89 | 1.77 |
| **M5_M** | 2.65 | 4.55 | 5.95 | 1.76 |
| **M5_S** | 2.59 | 3.72 | 4.43 | 2.08 |
| **Average** | 2.24 | 2.85 | 3.74 | 1.88 |

**Note: "mC" percent (%) represents the proportion of methylated cytosines to total cytosines on the whole genome. "mCpG percent (%)", "mCHG percent (%)", and "mCHH percent (%)" represent the proportion of methylated cytosines to total cytosines on the whole genome in CG, CHG and CHH subcontexts, respectively.**

**Table S21 Average coverage depth and methylation level of all the cytosines on the genome**

| Samples | C_covgMean | C (Mb) | CG (Mb) | CHG (Mb) | CHH (Mb) | meanC (%) | MeanCG (%) | MeanCHG (%) | MeanCHH (%) |
|---|---|---|---|---|---|---|---|---|---|
| **A13_M** | 7.5 | 2106.2 | 237.7 | 274.3 | 1594.2 | 17.98 | 68.04 | 44.93 | 5.87 |
| **A13_S** | 3.9 | 1092.3 | 231.8 | 180.3 | 680.2 | 38.96 | 85.92 | 72.17 | 14.16 |
| **A188_M** | 8.3 | 2329.1 | 263.4 | 302.5 | 1763.2 | 18.61 | 68.55 | 44.76 | 6.66 |
| **A188_S** | 4.7 | 1327.8 | 229.7 | 189.8 | 908.3 | 29.81 | 82.99 | 65.2 | 8.97 |
| **A65_M** | 5 | 1386.8 | 254.9 | 206.7 | 925.2 | 33.31 | 84.88 | 69.41 | 11.04 |
| **A65_S** | 8.6 | 2394.9 | 271.6 | 308 | 1815.4 | 19.28 | 68.56 | 46.58 | 7.27 |
| **AM206_H1** | 5.1 | 1414.2 | 241.7 | 202.2 | 970.2 | 30.84 | 84.1 | 67.01 | 10.04 |
| **AM206_H2** | 9 | 2512.1 | 273.3 | 324.3 | 1914.6 | 17.52 | 65.99 | 43.1 | 6.27 |
| **AM206_M1** | 6.5 | 1820.2 | 295.3 | 252.8 | 1272.1 | 29.71 | 83.61 | 65.74 | 10.03 |
| **AM206_M2** | 5.3 | 1475.1 | 258 | 211 | 1006.1 | 31.68 | 84.54 | 66.94 | 10.74 |
| **AM206_M3** | 5.1 | 1436.9 | 240.9 | 204.2 | 991.8 | 30.81 | 84.08 | 66.43 | 10.54 |
| **AM206_S1** | 6.2 | 1722.7 | 273.4 | 238 | 1211.3 | 25.19 | 80.59 | 58.57 | 6.12 |
| **AM206_S2** | 6.7 | 1887.1 | 303.5 | 265.6 | 1318 | 25.94 | 81.26 | 59.87 | 6.36 |
| **AM206_S3** | 5.6 | 1560.6 | 253.9 | 220.1 | 1086.6 | 26.17 | 81.55 | 60.25 | 6.32 |
| **G11_F** | 7.8 | 2190 | 246.9 | 282.3 | 1660.9 | 17.17 | 68.47 | 45.12 | 4.79 |
| **G11_S** | 5.1 | 1427.9 | 254.6 | 207.2 | 966.1 | 30.37 | 83.21 | 64.9 | 9.04 |
| **G186_F** | 8 | 2228.9 | 242.7 | 287.4 | 1698.8 | 17.19 | 66.88 | 44.89 | 5.41 |
| **G186_S** | 7.6 | 2122.9 | 234 | 273.3 | 1615.6 | 19.09 | 67.68 | 46.11 | 7.49 |
| **G21_F** | 8.9 | 2488 | 279.5 | 324.1 | 1884.3 | 17.35 | 66.79 | 44.3 | 5.38 |
| **G21_S** | 5.1 | 1412.9 | 254.4 | 206.7 | 951.9 | 32.93 | 83.83 | 67.39 | 11.85 |
| **M108_F** | 5.2 | 1467 | 248.8 | 208.4 | 1009.7 | 30.06 | 83.5 | 65.32 | 9.61 |
| **M108_M** | 7.5 | 2095.5 | 233.6 | 271.5 | 1590.3 | 18.08 | 68.35 | 45.39 | 6.03 |
| **M108_S** | 5.9 | 1653.4 | 284.4 | 240.4 | 1128.6 | 32.35 | 84.13 | 67.37 | 11.84 |
| **M168_F** | 7.7 | 2156.1 | 242.2 | 278.9 | 1634.9 | 18.21 | 68.62 | 45.18 | 6.14 |
| **M168_M** | 5.6 | 1555.1 | 262.2 | 221.5 | 1071.3 | 30.82 | 84.79 | 66.34 | 10.26 |
| **M168_S** | 4.8 | 1328.9 | 262.3 | 203.7 | 863 | 36.65 | 86.35 | 69.91 | 13.7 |
| **M5_F** | 9.1 | 2544 | 287.7 | 330.8 | 1925.5 | 17.37 | 67.87 | 44.52 | 5.16 |
| **M5_H** | 7.7 | 2156.7 | 249.4 | 279.2 | 1628.1 | 18.1 | 70.14 | 46.41 | 5.27 |
| **M5_M** | 8.7 | 2422 | 273 | 317.4 | 1831.6 | 16.51 | 66.96 | 43.3 | 4.34 |
| **M5_S** | 7.1 | 1990.6 | 226.2 | 256 | 1508.4 | 18.9 | 69.35 | 45.99 | 6.73 |
| **Average** | 6.64 | 1856.86 | 257.03 | 252.29 | 1347.54 | 24.90 | 76.39 | 56.11 | 8.11 |

Note: “C_covgMean” represents average coverage depth of all the cytosine sites on the genome; “C (Mb)” represents the number of bases aligned to the C position of the genome; “CG (Mb)”, “CHG (Mb)”, and “CHH (Mb)” represents the number of bases aligned to the cytosine in CG, CHG and CHH subcontexts of the genome, respectively. Mb represents megabase. “meanC (%)” represents average methylation level of all the cytosine sites on the genome. “MeanCG (%)”, “MeanCHG (%)”, and “MeanCHH (%)” represent average methylation levels in CG, CHG and CHH subcontexts of the genome, respectively.

**Table S22 Differential expression of DNA methyltransferase and demethylase genes between female (or hermaphroditic) and male floral buds in different comparative combinations**

| Gene ID | Functional annotation | Differential expression ($Log_2FC$) | | | |
|---|---|---|---|---|---|
| | | M_F vs. M_M | G_F vs. A_M | AM_M vs. AM_H | G_S vs. A_S |
| evm.model.Chr8.1809 | DNA (cytosine-5)-methyltransferase 4 OS=Arabidopsis thaliana GN=*MET4* PE=2 SV=1 | / | / | / | / |
| evm.model.Chr8.1810 | DNA (cytosine-5)-methyltransferase 1B OS=Oryza sativa subsp. japonica GN=*MET1B* PE=2 SV=1 | / | / | / | / |
| evm.model.Chr13.1774 | DNA (cytosine-5)-methyltransferase 2 OS=Zea mays GN=*ZMET5* PE=2 SV=1 | / | / | / | / |
| evm.model.Chr10.460 | DNA (cytosine-5)-methyltransferase CMT2 OS=Arabidopsis thaliana GN=*CMT2* PE=2 SV=3 | / | 1.18747 | -1.26831 | / |
| evm.model.Chr10.461 | DNA (cytosine-5)-methyltransferase CMT2 OS=Arabidopsis thaliana GN=*CMT2* PE=2 SV=3 | / | / | / | / |
| evm.model.Chr1.187 | DNA (cytosine-5)-methyltransferase CMT2 OS=Arabidopsis thaliana GN=*CMT2* PE=2 SV=3 | / | 1.20473 | / | / |
| evm.model.Chr2.2059 | DNA (cytosine-5)-methyltransferase DRM2 OS=Arabidopsis thaliana GN=*DRM2* PE=1 SV=1 | / | / | / | / |
| evm.model.Chr4.1896 | DNA (cytosine-5)-methyltransferase DRM2 OS=Arabidopsis thaliana GN=*DRM2* PE=1 SV=1 | / | / | / | / |
| evm.model.Chr4.1897 | DNA (cytosine-5)-methyltransferase DRM2 OS=Arabidopsis thaliana GN=*DRM2* PE=1 SV=1 | / | / | / | / |
| evm.model.Chr6.1098 | DNA (cytosine-5)-methyltransferase DRM2 OS=Arabidopsis thaliana GN=*DRM2* PE=1 SV=1 | / | / | / | / |
| evm.model.Chr6.1447 | DNA (cytosine-5)-methyltransferase DRM2 OS=Arabidopsis thaliana GN=*DRM2* PE=1 SV=1 | / | / | / | / |
| evm.model.Chr1.1637.6 | DNA-directed RNA polymerase V subunit 1 OS=Arabidopsis thaliana GN=*NRPE1* PE=1 SV=1 | / | / | -0.825986 | / |

| | | | | | |
|---|---|---|---|---|---|
| evm.model.Chr14.632 | Putative disease resistance protein RGA4 OS=Solanum bulbocastanum GN=*RGA4* PE=2 SV=1 | / | / | / | / |
| evm.model.Chr2.1873 | Uncharacterized protein At5g39570 OS=Arabidopsis thaliana GN=*At5g39570* PE=1 SV=1 | / | / | / | / |
| evm.model.Chr4.947 | Protein DCL, chloroplastic OS=Solanum lycopersicum GN=*DCL* PE=2 SV=1 | / | / | / | / |
| evm.model.Chr7.334 | DNA-directed RNA polymerase V subunit 5A OS=Arabidopsis thaliana GN=*NRPE5A* PE=1 SV=1 | / | / | / | / |
| evm.model.Chr8.1527.1 | DNA-directed RNA polymerase V subunit 7 OS=Arabidopsis thaliana GN=*NRPE7* PE=1 SV=1 | / | / | / | / |
| evm.model.Chr9.1005 | Protein DCL, chloroplastic OS=Solanum lycopersicum GN=*DCL* PE=2 SV=1 | / | / | / | / |
| evm.model.Chr9.878 | Protein DCL, chloroplastic OS=Solanum lycopersicum GN=*DCL* PE=2 SV=1 | / | / | / | / |
| evm.model.Chr1.2976 | Protein argonaute 4 OS=Arabidopsis thaliana GN=*AGO4* PE=1 SV=2 | / | / | / | / |
| evm.model.Chr13.1173 | Protein argonaute 4A OS=Oryza sativa subsp. japonica GN=*AGO4A* PE=2 SV=1 | / | / | -1.15965 | / |
| evm.model.Chr13.1174 | Protein argonaute 4 OS=Arabidopsis thaliana GN=*AGO4* PE=1 SV=2 | / | / | -1.22971 | / |
| evm.model.Chr9.935 | Protein argonaute 4A OS=Oryza sativa subsp. japonica GN=*AGO4A* PE=2 SV=1 | / | / | / | / |
| evm.model.Chr15.1689 | Protein ROS1 OS=Arabidopsis thaliana GN=*ROS1* PE=1 SV=2 | / | / | / | / |
| evm.model.Chr7.1501 | Protein ROS1 OS=Arabidopsis thaliana GN=*ROS1* PE=1 SV=2 | / | / | -0.808349 | / |
| evm.model.Chr5.929 | Transcriptional activator DEMETER OS=Arabidopsis thaliana GN=*DME* PE=1 SV=2 | / | / | 1.39878 | / |
| evm.model.Chr5.930 | Transcriptional activator DEMETER OS=Arabidopsis thaliana GN=*DME* PE=1 SV=2 | / | / | 1.68125 | / |
| evm.model.Chr15.491 | 2,3-dimethylmalate lyase OS=Eubacterium barkeri | / | / | / | / |

| | | | | | |
|---|---|---|---|---|---|
| | GN=*Dml* PE=1 SV=1 | | | | |
| evm.model.Chr8.1861 | 2,3-dimethylmalate lyase OS=Eubacterium barkeri GN=*Dml* PE=1 SV=1 | / | / | / | / |

Note: "/" represents no significant expression difference. "$Log_2FC$" represents $Log_2$(fold-change) in the whole text. "M_F vs. M_M", "G_F vs. A_M", and "AM_M vs. AM_H" represent M_F compared with M_M, G_F compared with A_M, and AM_M compared with AM_H, respectively, in the whole text.

**Table S23 Significant enrichment of genes whose gene body overlapped with DMRs in both single- and co-sex comparative combinations**

| Term type | GO terms | Significant enrichment of genes whose gene body overlapped with the following DMRs (**gene number; corrected *P*-value**) | | | | | |
|---|---|---|---|---|---|---|---|
| | | CG_hypo DMRs in M_F vs. M_M | CG_hypo DMRs in G_F vs. A_M | CHG_hypo DMRs in G_F vs. A_M | CHH_hypo DMRs in G_F vs. A_M | CG_hypo DMRs in AM_H vs. AM_M | CHG_hypo DMRs in AM_H vs. AM_M |
| Molecular function | RNA glycosylase activity | 4; 0.020 | / | / | / | / | / |
| | rRNA N-glycosylase activity | 4; 0.020 | / | / | / | / | / |
| | hydrolase activity, hydrolyzing N-glycosyl compounds | 4, 0.046 | / | / | / | / | / |
| | hydrolase activity | / | 219; 0.043 | / | 383; 0.001 | / | / |
| | protein tyrosine kinase activity | / | 79; 0.043 | / | / | / | / |
| | protein binding | / | 245; 0.046 | 36; 0.010 | / | / | / |
| | ADP binding | / | / | 14; 0.010 | / | / | / |
| | catalytic activity | / | / | / | 1055; 0.000 | / | / |
| | nucleoside-triphosphatase activity | / | / | / | 154; 0.007 | / | / |
| | pyrophosphatase activity | / | / | / | 156; 0.007 | / | / |
| | hydrolase activity, acting on acid anhydrides, in phosphorus-containing anhydrides | / | / | / | 158; 0.007 | / | / |
| | hydrolase activity, acting on acid anhydrides | / | / | / | 160; 0.007 | / | / |
| | oxidoreductase activity, acting on paired donors, with incorporation or | / | / | / | / | 23; 0.007 | / |

| | | | | | | | |
|---|---|---|---|---|---|---|---|
| | reduction of molecular oxygen | | | | | | |
| | iron ion binding | / | / | / | / | 19; 0.024 | / |
| | transcription factor activity, transcription factor binding | / | / | / | / | / | 7; 0.036 |
| | transcription cofactor activity | / | / | / | / | / | 7; 0.036 |
| Biological process | negative regulation of translation | 4, 0.020 | / | / | / | / | / |
| | negative regulation of cellular amide metabolic process | 4, 0.020 | / | / | / | / | / |
| | negative regulation of biosynthetic process | 4, 0.020 | / | / | / | / | / |
| | negative regulation of macromolecule biosynthetic process | 4, 0.023 | / | / | / | / | / |
| | negative regulation of cellular biosynthetic process | 4, 0.023 | / | / | / | / | / |
| | negative regulation of cellular macromolecule biosynthetic process | 4, 0.023 | / | / | / | / | / |
| | negative regulation of nitrogen compound metabolic process | 4, 0.026 | / | / | / | / | / |
| | negative regulation of cellular protein metabolic process | 4, 0.026 | / | / | / | / | / |
| | negative regulation of protein metabolic process | 4, 0.026 | / | / | / | / | / |
| | regulation of translation | 4, 0.026 | / | / | / | / | / |
| | posttranscriptional regulation of gene expression | 4, 0.026 | / | / | / | / | / |
| | regulation of cellular amide metabolic process | 4, 0.026 | / | / | / | / | / |
| | negative regulation of cellular metabolic process | 4, 0.000 | / | / | / | / | / |
| | ubiquitin-dependent protein catabolic process | / | / | 22; 0.000 | / | / | / |
| | modification-dependent protein catabolic process | / | / | 22; 0.000 | / | / | / |
| | modification-dependent | / | / | 22; 0.000 | / | / | / |

| | | | | | | | |
|---|---|---|---|---|---|---|---|
| | macromolecule catabolic process | | | | | | |
| | cellular protein catabolic process | / | / | 24; 0.001 | / | / | / |
| | proteolysis involved in cellular protein catabolic process | / | / | 24; 0.001 | / | / | / |
| | cellular macromolecule catabolic process | / | / | 25; 0.001 | / | / | / |
| | macromolecule catabolic process | / | / | 28; 0.003 | / | / | / |
| | protein catabolic process | / | / | 24; 0.005 | / | / | / |
| | organic substance catabolic process | / | / | 36; 0.010 | / | / | / |
| | catabolic process | / | / | 38; 0.023 | / | / | / |
| | macromolecule methylation | / | / | | / | / | 10; 0.036 |
| | methylation | / | / | | / | / | 10; 0.038 |
| Cellular component | cytoskeleton | 10; 0.038 | / | | / | / | |
| | nucleus | / | / | | / | / | 54; 0.032 |
| | membrane-bounded organelle | / | / | | / | / | 66; 0.036 |
| | intracellular membrane-bounded organelle | / | / | | / | / | 66; 0.036 |

Note: “/” represents no significant enrichment.

**Table S24 Statistics of transcript sequencing and mapping**

| Sample name | Raw reads | Clean reads | clean bases(G) | Error rate(%) | Q20(%) | Q30(%) | GC content(%) | Total mapped reads(%) | Uniquely mapped reads(%) | Non-splice reads(%) | Splice reads(%) |
|---|---|---|---|---|---|---|---|---|---|---|---|
| G11_S | 98,868,068 | 97,787,712 | 14.67 | 0.02 | 98.09 | 94.41 | 47.26 | 72.85 | 62.22 | 43.03 | 19.19 |
| G21_S | 81,822,254 | 80,768,974 | 12.12 | 0.02 | 98.1 | 94.48 | 47.96 | 72.76 | 65.37 | 46.8 | 18.58 |
| G186_S | 88,683,254 | 87,197,418 | 13.08 | 0.03 | 98.01 | 94.23 | 46.22 | 72.16 | 64.64 | 46.13 | 18.51 |
| G11_F | 85,683,726 | 84,569,460 | 12.69 | 0.03 | 97.99 | 94.22 | 47.77 | 72.28 | 64.46 | 46.14 | 18.32 |
| G21_F | 86,030,630 | 84,867,366 | 12.73 | 0.03 | 97.93 | 94.05 | 47.24 | 72.48 | 63.42 | 45.68 | 17.74 |
| G186_F | 87,308,606 | 86,284,288 | 12.94 | 0.03 | 97.7 | 93.5 | 47.99 | 72.81 | 62.9 | 47.09 | 15.81 |
| A13_S | 84,465,068 | 82,418,836 | 12.36 | 0.02 | 98.03 | 94.33 | 49.41 | 72.1 | 58.23 | 42.56 | 15.67 |
| A65_S | 105,061,238 | 103,137,842 | 15.47 | 0.02 | 98.04 | 94.31 | 49.16 | 74.21 | 64.21 | 47.94 | 16.27 |
| A188_S | 84,457,734 | 83,374,706 | 12.51 | 0.03 | 97.87 | 93.88 | 46.62 | 73.32 | 65.67 | 45.35 | 20.32 |
| A13_M | 92,985,930 | 91,507,886 | 13.73 | 0.02 | 98.11 | 94.46 | 49.32 | 68.87 | 60.34 | 45.3 | 15.04 |
| A65_M | 91,687,298 | 90,264,794 | 13.54 | 0.02 | 98.02 | 94.28 | 47.58 | 70.44 | 62.82 | 45.53 | 17.28 |
| A188_M | 96,545,780 | 95,038,466 | 14.26 | 0.02 | 98.1 | 94.5 | 49.63 | 71.88 | 63.65 | 46.67 | 16.98 |
| M5_F | 85,463,624 | 83,829,884 | 12.57 | 0.03 | 97.83 | 93.78 | 45.45 | 71.54 | 62.22 | 43.52 | 18.69 |
| M108_F | 100,535,034 | 99,004,114 | 14.85 | 0.03 | 97.95 | 94.1 | 45.81 | 69.99 | 62.31 | 43.55 | 18.76 |
| M136_F | 85,961,618 | 83,305,204 | 12.5 | 0.03 | 97.83 | 93.85 | 46.28 | 69.37 | 60.48 | 42.83 | 17.66 |
| M168_F | 90,119,882 | 88,283,026 | 13.24 | 0.03 | 97.85 | 93.92 | 45.48 | 70.84 | 62.32 | 44.07 | 18.25 |
| M5_M | 91,763,228 | 90,668,530 | 13.6 | 0.02 | 98.01 | 94.24 | 45.85 | 73.8 | 60.68 | 41.61 | 19.07 |
| M108_M | 93,231,950 | 91,255,670 | 13.69 | 0.02 | 98.07 | 94.33 | 47.66 | 70.65 | 57.81 | 43.59 | 14.22 |
| M136_M | 86,853,918 | 84,923,322 | 12.74 | 0.02 | 98.2 | 94.67 | 46.7 | 68.29 | 61.59 | 43.86 | 17.73 |
| M168_M | 83,739,456 | 81,732,790 | 12.26 | 0.02 | 98.15 | 94.55 | 47.17 | 69.87 | 62.65 | 45.54 | 17.11 |
| AM206_M1 | 85,441,398 | 82,610,270 | 12.39 | 0.02 | 98.13 | 94.53 | 47.49 | 70.5 | 65.54 | 45.26 | 20.28 |
| AM206_M2 | 87,539,372 | 86,069,394 | 12.91 | 0.02 | 98.2 | 94.69 | 46.84 | 70.7 | 65.57 | 44.48 | 21.09 |
| AM206_M3 | 82,042,672 | 80,760,548 | 12.11 | 0.02 | 98.1 | 94.46 | 47.7 | 70.66 | 65.69 | 44.92 | 20.77 |
| AM206_H1 | 81,265,378 | 80,349,558 | 12.05 | 0.03 | 97.97 | 93.99 | 46.22 | 69.77 | 63.11 | 43.18 | 19.93 |
| AM206_H2 | 111,500,364 | 110,398,622 | 16.56 | 0.03 | 97.85 | 93.81 | 46.31 | 69.49 | 62.15 | 43.12 | 19.03 |
| AM206_H3 | 82,562,330 | 81,865,868 | 12.28 | 0.03 | 97.93 | 94.07 | 46.2 | 69.95 | 63.37 | 43.39 | 19.97 |
| Average | 89,677,685 | 88,164,406 | 13.23 | 0.02 | 98 | 94.22 | 47.2 | 71.21 | 62.82 | 44.66 | 18.16 |
| Total | 2,331,619,810 | 2,292,274,548 | 343.85 | / | / | / | / | / | / | / | / |

**Table S25 Statistics of small RNA (smRNA) sequencing**

| Sample | Reads | Bases (G) | Error rate (%) | Q20 (%) | Q30 (%) | GC content (%) |
|---|---|---|---|---|---|---|
| G11_S | 20,753,281 | 1.038 | 0.01 | 97.75 | 95.16 | 50.47 |
| G21_S | 22,183,003 | 1.109 | 0.01 | 97.77 | 95.16 | 51.01 |
| G186_S | 21,053,296 | 1.053 | 0.01 | 97.85 | 95.40 | 49.43 |
| G11_F | 20,521,327 | 1.026 | 0.01 | 97.71 | 95.06 | 50.32 |
| G21_F | 23,726,752 | 1.186 | 0.01 | 97.49 | 94.51 | 49.98 |
| G186_F | 21,499,262 | 1.075 | 0.01 | 97.88 | 95.42 | 50.80 |
| A13_S | 20,803,542 | 1.040 | 0.01 | 97.84 | 95.32 | 50.98 |
| A65_S | 17,893,824 | 0.895 | 0.01 | 97.85 | 95.37 | 51.07 |
| A188_S | 16,771,483 | 0.839 | 0.01 | 97.90 | 95.43 | 51.98 |
| A13_M | 23,281,643 | 1.164 | 0.01 | 97.79 | 95.22 | 51.46 |
| A65_M | 18,425,519 | 0.921 | 0.01 | 97.77 | 95.19 | 50.50 |
| A188_M | 18,346,157 | 0.917 | 0.01 | 97.91 | 95.46 | 51.73 |
| M5_F | 22,995,555 | 1.150 | 0.01 | 97.88 | 95.46 | 48.23 |
| M108_F | 19,607,315 | 0.980 | 0.01 | 97.90 | 95.50 | 48.94 |
| M136_F | 24,117,369 | 1.206 | 0.01 | 97.81 | 95.31 | 49.22 |
| M168_F | 17,338,667 | 0.867 | 0.01 | 97.72 | 95.18 | 48.54 |
| M5_M | 17,439,006 | 0.872 | 0.01 | 97.78 | 95.26 | 47.88 |
| M108_M | 19,139,740 | 0.957 | 0.01 | 97.85 | 95.37 | 49.44 |
| M136_M | 16,783,027 | 0.839 | 0.01 | 97.82 | 95.33 | 49.31 |
| M168_M | 19,538,642 | 0.977 | 0.01 | 97.80 | 95.25 | 49.75 |
| AM206_M1 | 16,815,807 | 0.841 | 0.01 | 97.64 | 95.00 | 49.25 |
| AM206_M2 | 11,814,813 | 0.591 | 0.01 | 96.67 | 92.77 | 49.30 |
| AM206_M3 | 22,495,367 | 1.125 | 0.01 | 96.18 | 90.93 | 49.92 |
| AM206_H1 | 15,126,405 | 0.756 | 0.01 | 97.70 | 95.09 | 49.62 |
| AM206_H2 | 15,469,387 | 0.773 | 0.01 | 97.92 | 95.46 | 49.66 |
| AM206_H3 | 14,986,549 | 0.749 | 0.01 | 97.90 | 95.41 | 49.87 |
| Avarege | 19,189,489.92 | 0.959 | 0.01 | 97.7 | 95.00 | 49.95 |
| Total | 498,926,738 | 24.946 | / | / | / | / |

**Table S26 Clean reads of smRNA processing**

| Sample | total_reads | N% > 10% | low quality | 5_adapter_contamine | 3_adapter_null or insert_null | with ployA/T/G/C | clean reads |
|---|---|---|---|---|---|---|---|
| G11_S | 20,753,281 | 4482 (0.02%) | 44,965 (0.22%) | 65,588 (0.32%) | 3,379,399 (16.28%) | 53,554 (0.26%) | 17,205,293 (82.90%) |
| G21_S | 22,183,003 | 4844 (0.02%) | 51,267 (0.23%) | 45,076 (0.20%) | 332,708 (1.50%) | 74,420 (0.34%) | 21,674,688 (97.71%) |
| G186_S | 21,053,296 | 8348 (0.04%) | 53,455 (0.25%) | 31,050 (0.15%) | 771,219 (3.66%) | 31,093 (0.15%) | 20,158,131 (95.75%) |
| G11_F | 20,521,327 | 4555 (0.02%) | 44,705 (0.22%) | 39,425 (0.19%) | 1,122,804 (5.47%) | 67,406 (0.33%) | 19,242,432 (93.77%) |
| G21_F | 23,726,752 | 9550 (0.04%) | 65,007 (0.27%) | 37,442 (0.16%) | 724,493 (3.05%) | 79,241 (0.33%) | 22,811,019 (96.14%) |
| G186_F | 21,499,262 | 8644 (0.04%) | 51,434 (0.24%) | 53,442 (0.25%) | 352,244 (1.64%) | 53,468 (0.25%) | 20,980,030 (97.58%) |
| A13_S | 20,803,542 | 8361 (0.04%) | 54,187 (0.26%) | 38,158 (0.18%) | 372,161 (1.79%) | 65,836 (0.32%) | 20,264,839 (97.41%) |
| A65_S | 17,893,824 | 7090 (0.04%) | 46,544 (0.26%) | 40,159 (0.22%) | 347,825 (1.94%) | 47,114 (0.26%) | 17,405,092 (97.27%) |
| A188_S | 16,771,483 | 6693 (0.04%) | 38,845 (0.23%) | 47,157 (0.28%) | 288,495 (1.72%) | 40,809 (0.24%) | 16,349,484 (97.48%) |
| A12_M | 23,281,643 | 9253 (0.04%) | 52,829 (0.23%) | 52,201 (0.22%) | 1,016,877 (4.37%) | 54,885 (0.24%) | 22,095,598 (94.91%) |
| A65_M | 18,425,519 | 7299 (0.04%) | 46,357 (0.25%) | 57,479 (0.31%) | 426,804 (2.32%) | 42,910 (0.23%) | 17,844,670 (96.85%) |
| A188_M | 18,346,157 | 7300 (0.04%) | 45,458 (0.25%) | 31,497 (0.17%) | 873,432 (4.76%) | 39,748 (0.22%) | 17,348,722 (94.56%) |
| M5_F | 22,995,555 | 9184 (0.04%) | 60,638 (0.26%) | 23,473 (0.10%) | 210,678 (0.92%) | 72,735 (0.32%) | 22,618,847 (98.36%) |
| M108_F | 19,607,315 | 7787 (0.04%) | 53,098 (0.27%) | 24,183 (0.12%) | 209,901 (1.07%) | 46,867 (0.24%) | 19,265,479 (98.26%) |
| M136_F | 24,117,369 | 9618 (0.04%) | 54,949 (0.23%) | 21,589 (0.09%) | 292,397 (1.21%) | 15,501 (0.06%) | 23,723,315 (98.37%) |
| M168_F | 17,338,667 | 6345 (0.04%) | 70,855 (0.41%) | 24,960 (0.14%) | 324,173 (1.87%) | 44,682 (0.26%) | 16,867,652 (97.28%) |
| M5_M | 17,439,006 | 6899 (0.04%) | 46,058 (0.26%) | 18,244 (0.10%) | 168,246 (0.96%) | 66,801 (0.38%) | 17,132,758 (98.24%) |
| M108_M | 19,139,740 | 7798 (0.04%) | 42,697 (0.22%) | 13,209 (0.07%) | 440,488 (2.30%) | 6,011 (0.03%) | 18,629,537 (97.33%) |
| M136_M | 16,783,027 | 6602 (0.04%) | 38,479 (0.23%) | 15,128 (0.09%) | 664,293 (3.96%) | 11,334 (0.07%) | 16,047,191 (95.62%) |
| M168_M | 19,538,642 | 4219 (0.02%) | 42,570 (0.22%) | 19,856 (0.10%) | 514,863 (2.64%) | 13,418 (0.07%) | 18,943,716 (96.96%) |
| AM206_M1 | 16,815,807 | 3799 (0.02%) | 46,924 (0.28%) | 16,041 (0.10%) | 249,235 (1.48%) | 24,968 (0.15%) | 16,474,840 (97.97%) |
| AM206_M2 | 11,814,813 | 945 (0.01%) | 53,534 (0.45%) | 12,989 (0.11%) | 301,943 (2.56%) | 25,822 (0.22%) | 11,419,580 (96.65%) |
| AM206_M3 | 22,495,367 | 4849 (0.02%) | 61,155 (0.27%) | 19,829 (0.09%) | 404,512 (1.80%) | 26,332 (0.12%) | 21,978,690 (97.70%) |
| AM206_H1 | 15,126,405 | 3252 (0.02%) | 43,197 (0.29%) | 17,159 (0.11%) | 353,727 (2.34%) | 22,375 (0.15%) | 14,686,695 (97.09%) |
| AM206_H2 | 15,469,387 | 959 (0.01%) | 47,204 (0.31%) | 22,246 (0.14%) | 258,200 (1.67%) | 19,484 (0.13%) | 15,121,294 (97.75%) |
| AM206_H3 | 14,986,549 | 959 (0.01%) | 44,592 (0.30%) | 17,627 (0.12%) | 163,779 (1.09%) | 23,907 (0.16%) | 14,735,685 (98.33%) |
| Total | 498,926,738 | / | / | / | / | / | 481,025,277 (96.41%) |

**Table S27 Statistics of smRNA clean reads mapping**

| Sample | Total small RNA clean reads | Mapped smRNA | + Mapped sRNA | - Mapped sRNA |
|---|---|---|---|---|
| G11_S | 7,160,573 | 6,088,851 (85.03%) | 4,802,350 (67.07%) | 1,286,501 (17.97%) |
| G21_S | 15,343,724 | 13,072,196 (85.20%) | 9,785,419 (63.77%) | 3,286,777 (21.42%) |
| G186_S | 12,416,072 | 11,449,022 (92.21%) | 9,732,332 (78.38%) | 1,716,690 (13.83%) |
| G11_F | 10,150,467 | 8,615,952 (84.88%) | 6,398,538 (63.04%) | 2,217,414 (21.85%) |
| G21_F | 13,683,887 | 11,506,877 (84.09%) | 8,401,352 (61.40%) | 3,105,525 (22.69%) |
| G186_F | 13,785,791 | 11,814,579 (85.70%) | 9,083,181 (65.89%) | 2,731,398 (19.81%) |
| A13_S | 14,554,222 | 12,555,325 (86.27%) | 9,586,332 (65.87%) | 2,968,993 (20.40%) |
| A65_S | 12,597,917 | 11,076,168 (87.92%) | 8,694,168 (69.01%) | 2,382,000 (18.91%) |
| A188_S | 12,071,984 | 10,461,769 (86.66%) | 8,320,263 (68.92%) | 2,141,506 (17.74%) |
| A13_M | 15,749,154 | 13,528,838 (85.90%) | 10,239,860 (65.02%) | 3,288,978 (20.88%) |
| A65_M | 9,164,660 | 8,026,581 (87.58%) | 6,573,969 (71.73%) | 1,452,612 (15.85%) |
| A188_M | 12,023,291 | 10,232,268 (85.10%) | 7,705,657 (64.09%) | 2,526,611 (21.01%) |
| M5_F | 18,283,531 | 15,656,727 (85.63%) | 10,627,010 (58.12%) | 5,029,717 (27.51%) |
| M108_F | 16,245,072 | 14,059,966 (86.55%) | 9,882,133 (60.83%) | 4,177,833 (25.72%) |
| M136_F | 16,414,374 | 15,638,935 (95.28%) | 14,453,230 (88.05%) | 1,185,705 (7.22%) |
| M168_F | 12,482,195 | 10,683,974 (85.59%) | 7,587,615 (60.79%) | 3,096,359 (24.81%) |
| M5_M | 13,876,787 | 11,923,229 (85.92%) | 7,892,456 (56.88%) | 4,030,773 (29.05%) |
| M108_M | 12,271,609 | 11,625,042 (94.73%) | 10,853,440 (88.44%) | 771,602 (6.29%) |
| M136_M | 8,941,691 | 8,496,542 (95.02%) | 7,941,639 (88.82%) | 554,903 (6.21%) |
| M168_M | 12,527,033 | 11,853,682 (94.62%) | 10,847,783 (86.59%) | 1,005,899 (8.03%) |
| AM206_M1 | 11,213,075 | 10,288,560 (91.76%) | 8,687,362 (77.48%) | 1,601,198 (14.28%) |
| AM206_M2 | 7,146,184 | 6,344,395 (88.78%) | 5,016,140 (70.19%) | 1,328,255 (18.59%) |
| AM206_M3 | 14,529,796 | 13,381,058 (92.09%) | 10,725,693 (73.82%) | 2,655,365 (18.28%) |
| AM206_H1 | 9,757,517 | 8,760,438 (89.78%) | 7,067,391 (72.43%) | 1,693,047 (17.35%) |
| AM206_H2 | 10,840,203 | 9,812,926 (90.52%) | 7,984,091 (73.65%) | 1,828,835 (16.87%) |
| AM206_H3 | 11,058,384 | 9,776,262 (88.41%) | 7,506,348 (67.88%) | 2,269,914 (20.53%) |
| Total | 324,289,193 | 286,730,162 (88.42%) | 226,395,752 (69.81%) | 60,334,410 (18.61%) |

Note: “+ Mapped sRNA” and “- Mapped sRNA” represent reads mapped to the forward and reverse sequences of the reference genome, respectively.

**Table S28 Common up regulated miRNAs in female (or hermaphroditic) floral buds and their down regulated targets in single- and co-sex systems**

| miRNAs | Comparison of expression levels ($Log_2FC$) | | | Target mRNAs and functional annotation | Comparison of expression levels ($Log_2FC$) | | |
|---|---|---|---|---|---|---|---|
| | M_F vs. M_M | G_F vs. A_M | AM_M vs. AM_H | | M_F vs. M_M | G_F vs. A_M | AM_M vs. AM_H |
| osa-miR159a.1<br>ath-miR159a<br>lus-miR159b | 1.6679<br>1.7551<br>1.3304 | 1.2786<br>1.1429<br>1.0134 | -0.71849<br>-0.941<br>-0.73138 | evm.model.Chr14.73 (Exonuclease mut-7 homolog) | -0.885885 | -1.12888 | 1.24313 |
| osa-miR159a.1<br>ath-miR159a<br>lus-miR159b | 1.6679<br>1.7551<br>1.3304 | 1.2786<br>1.1429<br>1.0134 | -0.71849<br>-0.941<br>-0.73138 | evm.model.Chr11.1415 (NOZZLE) | -1.20511 | -2.5791 | 2.00008 |
| lus-miR159b<br>osa-miR159f<br>ptc-miR319e<br>ath-miR159a<br>ath-miR159b-3p<br>osa-miR159a.1 | 1.3304<br>/<br>1.6508<br>1.7551<br>1.3851<br>1.6679 | 1.0134<br>1.5704<br>1.3031<br>1.1429<br>1.0997<br>1.2786 | -0.73138<br>-0.92136<br>-1.953<br>-0.941<br>-0.75237<br>-0.71849 | evm.model.Chr13.1162; evm.model.Chr11.1032 (GAMYB) | -1.22572;<br>-4.80432 | -1.08237; / | 1.31184;<br>3.182 |
| osa-miR160e-5p<br>gma-miR160b<br>ath-miR160a-5p<br>csi-miR160 | /<br>/<br>/<br>/ | 1.7188<br>1.8218<br>1.832<br>1.8218 | -1.1056<br>-1.2671<br>-1.2981<br>-1.2736 | evm.model.Chr1.2406_evm.model.Chr1.2407; evm.model.Chr15.1795 (Auxin response factor 18) | / | -1.61897;<br>-1.5837 | 1.05192;<br>1.16987 |
| ath-miR159b-3p | 1.3851 | 1.0997 | -0.75237 | evm.model.Chr14.639 (Cinnamoyl-CoA reductase 2) | / | -2.10974 | 2.83561 |
| ptc-miR319e | 1.6508 | 1.3031 | -1.953 | evm.model.Chr1.16 (Myosin-11) | / | -1.87145 | 3.16934 |
| osa-miR159a.1<br>ath-miR159b-3p<br>ath-miR159a<br>lus-miR159b | 1.6679<br>1.3851<br>1.7551<br>1.3304 | 1.2786<br>1.0997<br>1.1429<br>1.0134 | -0.71849<br>-0.75237<br>-0.941<br>-0.73138 | evm.model.Chr12.1832.1 (NA) | -1.76297 | -1.1655 | / |

**Table S29 Common down regulated miRNAs in female (or hermaphroditic) floral buds and their up regulated targets in single- and co-sex systems**

| miRNAs | Comparison of expression | Target mRNAs | Comparison of expression levels |
|---|---|---|---|

| | levels ($Log_2FC$) | | | | ($Log_2FC$) | | |
|---|---|---|---|---|---|---|---|
| | M_F vs. M_M | G_F vs. A_M | AM_M vs. AM_H | | M_F vs. M_M | G_F vs. A_M | AM_M vs. AM_H |
| hbr-miR156<br>ath-miR157a-5p | /<br>/ | -4.9913<br>-4.9913 | 2.2267<br>2.2267 | evm.model.Chr14.920 (*SPL7*) | 1.68142 | 1.9433 | -2.86595 |
| hbr-miR156<br>ath-miR157a-5p<br>ath-miR156a-5p | /<br>/<br>-2.767 | -4.9913<br>-4.9913<br>-3.1252 | 2.2267<br>2.2267<br>1.0797 | evm.model.Chr7.129 (*SPL9*) | 1.33446 | 1.53419 | / |
| hbr-miR156<br>ath-miR157a-5p<br>ath-miR156a-5p | /<br>/<br>-2.767 | -4.9913<br>-4.9913<br>-3.1252 | 2.2267<br>2.2267<br>1.0797 | evm.model.Chr7.171 (*SPL17*) | 1.17788 | 1.20547 | / |
| ath-miR156a-5p | -2.767 | -3.1252 | 1.0797 | evm.model.Chr2.176 (*JMJ25*) | / | 1.93947 | -4.28718 |
| hbr-miR156<br>ath-miR157a-5p | /<br>/ | -4.9913<br>-4.9913 | 2.2267<br>2.2267 | evm.model.Chr12.2581 (*MFP1-1*) | / | 1.12136 | -1.90103 |
| osa-miR396g | -1.6262 | -1.3292 | 0.95526 | evm.model.Chr4.1040 (Growth-regulating factor 6) | / | 1.35027 | -0.992217 |
| hbr-miR156 | / | -4.9913 | 2.2267 | evm.model.Chr1.2836 (*ORCS-1A*) | / | 1.80364 | -1.59282 |
| hbr-miR156<br>ath-miR156a-5p | /<br>-2.767 | -4.9913<br>-3.1252 | 2.2267<br>1.0797 | evm.model.Chr6.168 (*BHLH25*) | / | 1.87414 | -2.31547 |
| hbr-miR156 | / | -4.9913 | 2.2267 | evm.model.Chr6.1694 (*PCS1*) | / | 1.952 | -2.53158 |
| tae-miR395b<br>ath-miR395a | /<br>/ | -6.0795<br>-6.3978 | 1.3101<br>1.3609 | evm.model.Chr4.294 (ATP sulfurylase 1, chloroplastic) | / | 1.27484 | -1.07331 |
| hbr-miR156 | / | -4.9913 | 2.2267 | evm.model.Chr9.754 (Fanconi anemia group M protein homolog) | / | 1.12718 | -1.86662 |
| hbr-miR156<br>ath-miR157a-5p | /<br>/ | -4.9913<br>-4.9913 | 2.2267<br>2.2267 | evm.model.Chr1.2968 (Cucumisin) | / | 2.07241 | -2.0509 |
| hbr-miR156<br>ath-miR157a-5p<br>ath-miR156a-5p | /<br>/<br>-2.767 | -4.9913<br>-4.9913<br>-3.1252 | 2.2267<br>2.2267<br>1.0797 | evm.model.Chr12.765 (*SPL16*) | / | 1.17253 | -2.7081 |
| vvi-miR396b<br>mdm-miR396a | /<br>/ | -1.1362<br>-2.2734 | 0.67247<br>0.58789 | evm.model.Chr12.864 (*SOBIR1*) | / | 0.99355<br>2 | -1.24859 |

Note: "/" represents no significant expression difference.

**Table S30 Samples for WGCNA analysis**

| Groups | Samples | Trait value (female and male) |
|---|---|---|
| Female | G_11F | 0 |
| | G_21F | 0 |
| | G_186F | 0 |
| | M108_F | 0 |
| | M136_F | 0 |
| | M168_F | 0 |
| | PM_SF | 0 |
| Male | A13_M | 1 |
| | A65_M | 1 |
| | A188_M | 1 |
| | M108_M | 1 |
| | M136_M | 1 |
| | M168_M | 1 |
| | AM206_M1 | 1 |
| | AM206_M2 | 1 |
| | AM206_M3 | 1 |
| | PM_LF | 1 |
| Hermaphrodite | AM206_H1 | / |
| | AM206_H2 | / |
| | AM206_H3 | / |

**Table S31 Location and Category of SNPs and indels before LD pruning of 90 plants with male flower production**

| Category | | Number of SNPs |
|---|---|---|
| Upstream (located in 1 Kb upstream of a gene) | | 840,435 |
| Exonic | Stop gain | 3,943 |
| | Stop loss | 593 |
| | Synonymous | 166,097 |
| | Non-synonymous | 184,950 |
| Intronic | | 5,285,375 |
| Splicing (located in a 2 bp region in intron close to the boundary of exon and intron) | | 2,215 |
| Downstream (located in 1 Kb downstream of a gene) | | 689,012 |
| Upstream/downstream (located in 1 Kb upstream of a gene and in 1Kb downstream of another gene) | | 80,246 |
| Intergenic | | 16,585,800 |
| ts (transitions) | | 17,889,632 |
| tv (transversions) | | 6,233,531 |
| ts/tv (ratio of transitions to transversions) | | 2.869 |
| Total SNPs and Indels | | 24,123,163 |

Note: Total SNPs and Indels after LD pruning was 3,502,197 and 311,512, respectively, which were used for GWAS.

**Table S32 Genotypes of top 10 SNPs with the proportion of female immature branches as phenotype in 90 plants with male flower production**

| Chromosome | Chr7 | Chr7 | Chr7 | Chr7 | Chr7 | Chr7 | Chr7 | Chr7 | Chr7 | Chr7 | Proportion of female shoots |
|---|---|---|---|---|---|---|---|---|---|---|---|
| Position | 29256366 | 29276632 | 29317389 | 29323533 | 29083270 | 29333185 | 29269527 | 29358184 | 29314787 | 29315847 | / |
| Ref | C | A | T | G | T | G | G | A | T | T | / |
| 10 | TC | G | C | T | T | C | G | G | C | C | 0.2 |
| 13 | T | G | C | T | T | C | G | G | C | C | 0 |
| 24 | T | G | C | T | TC | C | G | G | C | C | 0.1 |
| 26 | T | GC | C | T | T | C | G | G | C | C | 0.1 |
| 28 | T | GC | C | T | T | C | G | G | C | C | 0.2 |
| 31 | T | G | C | T | T | C | G | G | C | C | 0.2 |
| 34 | T | G | C | T | T | C | G | G | C | C | 0 |
| 40 | T | GC | C | T | T | C | G | G | C | C | 0.2 |
| 44 | T | G | C | T | T | C | G | G | C | C | 0.2 |
| 45 | T | G | C | T | T | C | G | G | C | C | 0 |
| 47 | T | GC | C | T | T | C | G | G | C | C | 0.5 |
| 50 | T | G | CA | T | T | C | G | G | C | C | 0.1 |
| 83 | T | G | C | T | T | C | G | G | C | C | 0 |
| 87 | TC | AG | TC | TG | CT | GC | G | AG | TC | TC | 0.5 |
| 8 | T | G | C | T | T | C | G | G | C | C | 0.2 |
| 108 | T | GC | C | T | T | C | G | G | C | C | 0.2 |
| 109 | T | G | C | T | T | C | G | G | C | C | 0.5 |
| 111 | T | G | C | T | T | C | G | G | C | C | 0.2 |
| 118 | T | G | C | T | T | C | G | G | C | C | 0.5 |
| 121 | TC | GA | CT | GT | CT | GC | G | GA | CT | TC | 0.5 |
| 122 | TC | GAC | TC | GT | CT | CG | G | GA | TC | TC | 0.5 |
| 125 | T | G | C | T | T | C | G | G | C | C | 0 |
| 127 | T | G | C | T | T | C | G | G | C | C | 0 |
| 133 | T | G | C | T | T | C | G | G | C | C | 0 |
| 136 | T | G | C | T | T | C | G | G | C | C | 0.4 |
| 52 | T | G | C | T | T | C | G | G | C | C | 0.2 |
| 53 | T | GC | C | T | T | C | G | G | C | C | 0.2 |
| 56 | T | G | C | T | T | C | G | G | C | C | 0 |
| 57 | T | G | C | T | T | C | G | G | C | C | 0 |

| | | | | | | | | | | |
|---|---|---|---|---|---|---|---|---|---|---|
| 59 | T | G | CA | T | T | C | G | G | C | C | 0 |
| 65 | T | G | C | T | T | C | G | G | C | C | 0 |
| 68 | T | G | C | T | T | C | G | G | C | C | 0 |
| 69 | T | G | C | T | T | C | G | G | C | C | 0 |
| 70 | T | G | C | T | T | C | G | G | C | C | 0 |
| 74 | T | GC | C | T | T | C | G | G | C | C | 0.2 |
| 90 | T | G | C | T | T | C | G | G | C | C | 0.2 |
| 93 | T | G | C | T | T | C | G | G | C | C | 0 |
| 95 | T | G | C | T | T | C | G | G | C | C | 0 |
| 96 | T | G | C | T | T | C | G | G | C | C | 0 |
| 97 | T | G | C | T | T | C | G | G | C | C | 0 |
| 98 | T | G | C | T | T | C | G | G | C | C | 0 |
| 99 | T | G | C | T | T | C | G | G | C | C | 0 |
| 135 | TC | AG | TC | GT | TC | CG | G | AG | CT | TC | 0.5 |
| 142 | T | GC | C | T | T | C | G | G | C | C | 0 |
| 145 | T | GC | C | T | T | C | G | G | C | C | 0 |
| 146 | T | G | C | T | T | C | G | G | C | C | 0 |
| 149 | T | G | C | T | T | C | G | G | C | C | 0 |
| 155 | T | GC | C | T | T | C | G | G | C | C | 0 |
| 157 | TC | GA | TC | GT | TC | GC | G | GA | CT | TC | 0 |
| 159 | T | G | C | T | T | C | G | G | C | C | 0 |
| 161 | T | GC | C | T | T | C | G | G | C | C | 0 |
| 162 | T | G | C | T | T | C | G | G | C | C | 0 |
| 167 | T | GC | C | T | T | C | G | G | C | C | 0 |
| 168 | T | GC | C | T | T | C | G | G | C | C | 0.4 |
| 173 | CT | GAC | CT | GT | TC | CG | G | AG | CT | TC | 0.4 |
| 174 | T | G | C | T | T | C | G | G | C | C | 0 |
| 177 | T | G | C | T | T | C | G | G | C | C | 0 |
| 179 | TC | AG | CT | TG | CT | CG | G | AG | TC | CT | 0.7 |
| 180 | TC | GA | CT | GT | CT | GC | G | A | CT | CT | 0.6 |
| 185 | T | GC | C | T | T | C | G | G | C | C | 0 |
| 188 | T | GC | C | T | T | C | G | G | C | C | 0 |
| 193 | T | G | C | T | T | C | G | G | C | C | 0 |
| 201 | T | G | C | T | T | C | G | G | C | C | 0 |

| | | | | | | | | | | |
|---|---|---|---|---|---|---|---|---|---|---|
| 206 | T | G | C | T | T | C | G | G | C | C | 0.01 |
| 208 | T | G | C | T | T | C | G | G | C | C | 0 |
| 211 | T | G | C | T | T | C | G | G | C | C | 0.3 |
| 113 | T | GC | C | T | T | C | G | G | C | C | 0 |
| 128 | T | G | C | T | T | C | G | G | C | C | 0 |
| 130 | T | G | C | T | T | C | G | GA | C | C | 0 |
| 139 | T | G | CA | T | T | C | G | G | C | C | 0 |
| 154 | T | G | C | T | T | C | G | G | C | C | 0 |
| 164 | T | GC | CA | T | T | C | G | G | C | C | 0 |
| 169 | T | G | C | T | T | C | G | G | C | C | 0 |
| 170 | T | G | C | T | T | C | G | G | C | C | 0 |
| 182 | T | GT | C | T | T | C | G | G | C | C | 0.05 |
| 187 | T | G | C | T | T | C | G | G | C | C | 0 |
| 189 | T | G | C | T | T | C | G | G | C | C | 0.02 |
| 19 | T | G | C | T | T | C | G | G | C | C | 0.1 |
| 204 | T | G | C | T | T | C | G | G | C | C | 0 |
| 210 | T | G | C | T | T | C | G | G | C | C | 0.1 |
| 41 | T | G | C | T | T | C | G | G | C | C | 0.2 |
| 107 | CT | GA | TC | TG | TC | CG | G | AG | CT | CT | 0.1 |
| 137 | T | G | C | T | T | C | G | G | C | C | 0.1 |
| 150 | T | G | C | T | T | C | G | G | C | C | 0 |
| 166 | T | G | C | T | T | C | G | G | C | C | 0 |
| 184 | T | G | C | T | T | C | G | G | C | C | 0 |
| 207 | T | G | C | T | T | C | G | G | C | C | 0 |
| 42 | T | G | C | T | T | C | G | G | C | C | 0.1 |
| 67 | T | G | C | T | T | C | G | G | C | C | 0.05 |
| 84 | T | G | C | T | T | C | G | G | C | C | 0.3 |

Note: the single letters represent homozygotes.

**Table S33 Genes in the genomic regions of Chr7: 29.0-29.4 Mb**

| Serial number | Gene ID | Genomic Region (Mb) | Gene name with Swissprot Annotation | Comparison of expression levels (M_F vs. M_M) | Comparison of expression levels (G_F vs. A_M) | Comparison of methylation levels (M_F vs. M_M) |
|---|---|---|---|---|---|---|
| 1 | evm.model.Chr7.976 | Chr7: 29.0-29.4 | *At3g47200* | / | / | / |
| 2 | evm.model.Chr7.977 | | *At3g47200* | / | / | / |
| 3 | evm.model.Chr7.978 | | *At3g47200* | / | 2.40182 | / |
| 4 | evm.model.Chr7.979 | | *At3g47200* | / | 1.35052 | / |
| 5 | evm.model.Chr7.980 | | *At3g47200* | / | 1.98539 | / |
| 6 | evm.model.Chr7.981 | | *At3g47200* | / | 1.02959 | / |
| 7 | evm.model.Chr7.983 | | *At3g47200* | 2.3662 | 2.72264 | / |

Note: "$Log_2FC$" values are used to represent the significant difference of expression levels in the comparative combinations, "/" represents no significant expression difference. This criterion is consistent in the following tables.

**Table S34 miRNA in the genomic regions of Chr7: 29.0-29.4 Mb**

| Serial number | Gene ID | Genomic Region (Mb) | Comparison of expression levels (M_F vs M_M) |
|---|---|---|---|
| 1 | pab-miR3711 | Chr7: 29.0-29.4 | -2.2242 |

**Table S35 Genotypes of top 13 SNPs in the peaks of Manhattan plot with the proportion of hermaphroditic immature floral branches as phenotype in 90 male-biased samples**

| Chromosome | Chr2 | Chr2 | Chr11 | Chr11 | Chr11 | Chr11 | Chr11 | Chr11 | Chr14 | Chr14 | Chr14 | Chr14 | Chr14 | Proportion of hermaphroditic shoots |
|---|---|---|---|---|---|---|---|---|---|---|---|---|---|---|
| Pos | 30821709 | 30845253 | 20659872 | 21065164 | 21061959 | 21063195 | 21079272 | 21083058 | 31311437 | 30967061 | 30940866 | 31300726 | 30979995 | / |
| Ref | G | C | G | G | G | G | A | C | A | C | G | A | C | / |
| 10 | G | C | G | G | G | G | A | C | G | TC | AG | C | C | 0 |
| 13 | GA | CA | G | G | G | G | A | C | G | T | GA | C | C | 0.2 |
| 24 | G | C | GC | G | G | G | A | C | G | C | G | C | C | 0.1 |
| 26 | AG | CA | G | G | G | G | A | C | G | CT | AG | C | GC | 0.1 |
| 28 | G | C | G | G | G | G | A | C | G | C | G | C | C | 0 |
| 31 | G | C | G | G | G | G | A | C | G | C | G | CA | C | 0 |
| 34 | G | C | G | G | G | G | A | C | G | C | G | C | C | 0.2 |
| 40 | G | C | G | G | G | G | A | C | G | C | G | C | C | 0 |
| 44 | G | C | G | G | G | G | A | C | G | C | G | C | C | 0 |
| 45 | G | C | CG | G | G | G | A | C | G | C | G | C | C | 0 |
| 47 | G | C | G | G | G | G | A | C | G | C | G | C | C | 0 |
| 50 | G | C | G | G | G | G | A | C | G | C | G | C | C | 0.1 |
| 83 | G | C | G | G | G | G | A | C | G | C | G | C | C | 0.1 |
| 87 | G | C | GC | G | G | G | A | C | G | C | G | C | C | 0 |
| 8 | G | C | G | G | G | G | A | C | G | TC | GA | C | C | 0 |
| 108 | G | C | CG | G | G | G | A | C | G | C | G | C | C | 0 |
| 109 | G | C | GC | G | G | G | A | C | G | C | G | C | C | 0 |
| 111 | G | C | G | CG | AG | AG | GA | GC | G | C | G | C | C | 0 |
| 118 | G | C | G | G | G | G | A | C | G | C | G | C | C | 0 |
| 121 | G | C | G | G | G | G | A | C | G | C | G | CA | C | 0 |
| 122 | G | C | G | G | G | G | A | C | G | C | G | CA | C | 0 |
| 125 | G | C | G | CG | AG | AG | AG | GC | G | C | G | C | C | 0 |
| 127 | G | C | G | CG | GA | AG | GA | GC | G | C | G | C | C | 0 |
| 133 | G | C | G | CG | GA | AG | GA | GC | G | C | G | C | C | 0 |
| 136 | G | C | GC | C | A | A | G | G | G | C | G | C | C | 0.2 |
| 52 | G | C | G | G | G | G | A | C | G | C | G | C | C | 0 |

| | | | | | | | | | | | | | | |
|---|---|---|---|---|---|---|---|---|---|---|---|---|---|---|
| 53 | G | C | G | G | G | G | A | C | G | C | G | C | C | 0 |
| 56 | G | C | G | G | G | GA | A | C | G | C | G | C | C | 0 |
| 57 | * | C | G | G | G | G | A | C | G | C | G | C | C | 0 |
| 59 | G | C | G | G | G | G | A | C | G | C | G | C | C | 0 |
| 65 | G | C | G | G | G | G | A | C | G | C | G | C | C | 0 |
| 68 | G | C | G | G | G | G | A | C | G | C | G | C | C | 0 |
| 69 | G | C | G | G | G | G | A | C | G | C | G | C | C | 0 |
| 70 | G | C | G | G | G | G | A | C | G | C | G | C | C | 0 |
| 74 | G | C | G | G | G | G | A | C | G | C | G | C | C | 0 |
| 90 | G | C | G | G | G | G | A | C | G | C | G | C | C | 0 |
| 93 | G | C | G | G | G | G | A | C | G | C | G | C | C | 0 |
| 95 | G | C | G | G | G | G | A | C | G | C | G | C | C | 0 |
| 96 | G | C | G | G | G | G | A | C | G | C | G | * | C | 0 |
| 97 | G | C | G | G | G | G | A | C | G | C | G | * | C | 0 |
| 98 | G | C | G | G | G | G | A | C | G | C | G | C | C | 0 |
| 99 | G | C | G | G | G | G | A | C | G | C | G | C | C | 0 |
| 135 | G | C | G | GC | GAT | AG | GA | GC | G | C | G | C | C | 0 |
| 142 | G | C | G | G | G | G | A | C | G | C | G | C | C | 0 |
| 145 | G | C | G | G | GT | G | A | C | G | C | G | C | C | 0 |
| 146 | G | C | G | G | G | G | A | C | G | C | G | C | C | 0 |
| 149 | G | C | G | G | G | G | A | C | G | C | G | C | C | 0 |
| 155 | G | C | G | G | G | G | A | C | G | C | G | * | C | 0 |
| 157 | G | C | G | G | G | G | A | C | G | C | G | * | C | 0 |
| 159 | G | C | G | G | G | G | A | C | G | C | G | C | C | 0 |
| 161 | G | C | G | G | G | G | A | C | GA | TC | AG | AC | CG | 0 |
| 162 | G | C | G | G | G | G | A | C | G | C | G | C | C | 0 |
| 167 | GA | AC | G | G | G | G | A | C | AG | CT | AG | CA | CG | 0.2 |
| 168 | AG | AC | G | G | G | G | A | C | AG | CT | AG | A | GC | 0.2 |
| 173 | G | C | CG | G | G | G | A | C | AG | CT | GA | CA | CG | 0.2 |
| 174 | G | C | G | G | G | G | A | C | G | C | G | C | C | 0 |
| 177 | G | C | G | G | G | G | A | C | AG | CT | AG | A | CG | 0 |
| 179 | G | C | G | G | G | G | A | C | G | CT | A | C | CG | 0 |
| 180 | G | C | G | G | G | G | A | C | G | C | G | C | C | 0 |
| 185 | G | C | G | G | G | G | A | C | G | C | G | C | C | 0 |

| | | | | | | | | | | | | | | |
|---|---|---|---|---|---|---|---|---|---|---|---|---|---|---|
| 188 | G | C | G | G | G | G | A | C | G | C | G | C | C | 0 |
| 193 | AG | CA | G | G | G | G | A | C | G | C | G | CA | C | 0 |
| 201 | G | C | G | G | G | G | A | C | G | C | G | C | C | 0 |
| 206 | GA | AC | C | C | A | A | G | G | AG | T | A | A | GC | 0.33 |
| 208 | G | C | G | G | G | G | A | C | G | C | G | C | C | 0 |
| 211 | G | C | CG | G | G | G | A | C | G | CT | GA | C | C | 0.2 |
| 113 | G | C | G | G | G | G | A | C | G | C | G | C | C | 0 |
| 128 | G | C | G | G | G | G | A | C | G | C | G | C | C | 0 |
| 130 | GA | CA | G | C | A | A | G | G | GA | C | G | AC | C | 0.3 |
| 139 | G | C | G | G | G | G | A | C | G | C | G | C | C | 0 |
| 154 | G | C | G | G | G | G | A | C | G | C | G | C | C | 0 |
| 164 | G | C | G | G | G | G | A | C | G | C | G | C | C | 0 |
| 169 | G | C | G | G | G | G | A | C | G | C | G | C | C | 0 |
| 170 | G | C | G | G | G | G | A | C | G | C | G | C | C | 0 |
| 182 | G | C | G | G | G | G | A | C | G | CT | GA | C | CG | 0 |
| 187 | G | C | G | G | G | G | A | C | G | TC | GA | C | GC | 0 |
| 189 | G | C | G | G | G | G | A | C | G | TC | GA | C | GC | 0 |
| 19 | G | C | G | G | G | G | A | C | G | C | G | C | C | 0 |
| 204 | G | C | G | G | G | G | A | C | G | C | G | C | C | 0 |
| 210 | G | C | G | G | G | G | A | C | G | C | G | C | C | 0.1 |
| 41 | G | C | G | G | G | G | A | C | G | C | G | * | C | 0 |
| 107 | G | C | G | G | G | G | A | C | GA | C | G | C | C | 0.15 |
| 137 | G | C | G | G | G | G | A | C | G | C | G | C | C | 0.1 |
| 150 | G | C | G | G | G | GT | A | C | G | C | G | C | C | 0 |
| 166 | AG | AC | CG | GC | GA | GA | AG | CG | GA | CT | A | A | GC | 0.3 |
| 184 | G | C | G | G | G | G | A | C | G | CT | GA | C | C | 0 |
| 207 | G | C | G | G | G | G | A | C | G | C | G | C | C | 0 |
| 42 | G | C | G | G | G | G | A | C | G | C | G | C | C | 0 |
| 67 | G | C | G | G | G | G | A | C.+1A | G | CT | G | C | C | 0 |
| 84 | G | C | G | G | G | G | A | C | G | C | G | C | C | 0 |

Note: the single letters represent homozygotes, and "*" represent missing.

**Table S36 Genes in the hermaphrodite production-associated regions**

| Serial number | Gene ID | Genomic Region (Mb) | Gene name | Comparison of expression levels (AM_M vs. AM_H) |
|---|---|---|---|---|
| 1 | evm.model.Chr2.407 | Chr2: 25.4-25.9 | *NA* | / |
| 2 | evm.model.Chr2.489 | Chr2: 28.6-28.75 | *NAC082* | / |
| 3 | evm.model.Chr2.490 | | *NA* | -4.0365 |
| 4 | evm.model.Chr2.491 | | *DCLRE1B* | / |
| 5 | evm.model.BG.188.p | | | / |
| 6 | evm.model.Chr2.493 | | *GONST1* | / |
| 7 | evm.model.Chr2.494 | | *NA* | / |
| 8 | evm.model.Chr2.495 | | *NA* | / |
| 9 | evm.model.Chr2.496 | | *NA* | / |
| 10 | evm.model.Chr2.497 | | *NA* | / |
| 11 | evm.model.Chr2.498 | | *NA* | / |
| 12 | evm.model.Chr2.499 | | *HDA14* | / |
| 13 | evm.model.Chr2.505 | Chr2: 28.9-29.2 | *NA* | / |
| 14 | evm.model.Chr2.507 | | *PMT26* | 1.62766 |
| 15 | evm.model.Chr2.509 | | *NA* | / |
| 16 | evm.model.Chr2.510 | | *PI4KB1* | / |
| 17 | evm.model.Chr2.511 | | *NA* | / |
| 18 | evm.model.Chr2.512 | | *At1g54610* | 2.23417 |
| 19 | evm.model.Chr2.513 | | *TKI1* | / |
| 20 | evm.model.Chr2.521 | Chr2: 29.62-29.83 | *SG1* | / |
| 21 | evm.model.Chr2.522 | | *NA* | / |
| 22 | evm.model.Chr2.523 | | *At1g10490* | / |
| 23 | evm.model.Chr2.524 | | *At1g10490* | / |
| 24 | evm.model.Chr2.526 | | *At2g21870* | / |
| 25 | evm.model.Chr2.564 | Chr2: 30.78-30.9 | Cytochrome P450 CYP736A12 | 1.65906 |
| 26 | evm.model.Chr2.567 | | *At5g10020* | / |
| 27 | evm.model.Chr2.568 | | Cytochrome P450 CYP736A12 | / |
| 28 | evm.model.Chr11.614 | Chr11: 20.6-20.7 | *At1g06470* | / |
| 29 | evm.model.Chr11.616 | | *At3g12360* | / |

| | | | | |
|---|---|---|---|---|
| 30 | evm.model.Chr11.617 | | *At3g12360* | -5.7288 |
| 31 | evm.model.Chr11.630 | Chr11: 21.0-21.12 | *ASP1* | / |
| 32 | evm.model.Chr11.631 | | *GSVIVT00023967 001* | / |
| 33 | evm.model.Chr11.633.1 | | *rplA* | / |
| 34 | evm.model.Chr11.634 | | *LBD12* | / |
| 35 | evm.model.Chr11.635 | | *LBD12* | / |
| 36 | evm.model.Chr14.711 | Chr14: 8.65-8.95 | Furcatin hydrolase | / |
| 37 | evm.model.Chr14.712 | | *BGLU12* | / |
| 38 | evm.model.Chr14.713 | | *BGLU27* | / |
| 39 | evm.model.Chr14.716 | | *BGLU12* | / |
| 40 | evm.model.Chr14.717 | | *BGLU12* | / |
| 41 | evm.model.BG.486 | Chr14: 9.80-9.95 | | / |
| 42 | evm.model.Chr14.766 | | *LHP1* | / |
| 43 | evm.model.Chr14.767 | | *NA* | / |
| 44 | evm.model.Chr14.768 | | *NA* | / |
| 45 | evm.model.Chr14.769 | | *trmB* | / |
| 46 | evm.model.Chr14.770 | | Glycine-rich protein A3 | / |
| 47 | evm.model.Chr14.771 | | Glycine-rich protein A3 | / |
| 48 | evm.model.Chr14.772 | | *Os04g0386900* | / |
| 49 | evm.model.Chr14.773 | | *NA* | / |
| 50 | evm.model.Chr14.774 | | *NA* | / |
| 51 | evm.model.Chr14.1526 | Chr14: 30.9-31.4 | *NA* | / |
| 52 | evm.model.Chr14.1528 | | *GGR* | / |
| 53 | evm.model.Chr14.1529 | | *FBL10* | / |
| 54 | evm.model.Chr14.1530 | | *HT1* | / |
| 55 | evm.model.Chr14.1531 | | *HT1* | / |
| 56 | evm.model.Chr14.1532 | | *NA* | / |
| 57 | evm.model.Chr14.1533 | | *MSL8* | / |
| 58 | evm.model.Chr14.1534 | | *ATR* | / |

| | | | | |
|---|---|---|---|---|
| 59 | evm.model.Chr14.1535 | | *At2g17033* | / |
| 60 | evm.model.Chr14.1536 | Chr14: 31.4-31.5 | *Dcaf8* | / |
| 61 | evm.model.Chr14.1537 | | *GLT3* | / |
| 62 | evm.model.Chr14.1538 | | *tetA* | / |
| 63 | evm.model.Chr14.1539 | | *NA* | / |
| 64 | evm.model.Chr14.1541 | | *NA* | / |
| 65 | evm.model.Chr14.1542 | | *SMT2* | / |

Note: “NA” represents not available. This criterion is consistent in the following tables.

**Table S37 lncRNAs in the hermaphrodite production-associated regions**

| Serial number | Gene ID | Genomic Region (Mb) | Comparison of expression levels (AM_M vs. AM_H) |
|---|---|---|---|
| 1 | LNC_004284 | Chr2: 25.4-25.9 | / |
| 2 | LNC_004283 | | / |
| 3 | LNC_004041 | | / |
| 4 | LNC_004301 | Chr2: 30.78-30.9 | / |
| 5 | LNC_003094 | Chr14: 8.65-8.95 | / |
| 6 | LNC_003095 | | / |
| 7 | LNC_003290 | | / |
| 8 | LNC_003098 | Chr14: 9.80-9.95 | / |
| 9 | LNC_003099 | | / |
| 10 | LNC_003102 | | / |
| 11 | LNC_003103 | | 3.04716 |
| 12 | LNC_003100 | | / |
| 13 | LNC_003101 | | / |
| 14 | LNC_003104 | | / |
| 15 | LNC_003397 | Chr14: 30.9-31.4 | / |
| 16 | LNC_003396 | | / |
| 17 | LNC_003395 | | / |
| 18 | LNC_003399 | | / |
| 19 | LNC_003398 | | / |
| 20 | LNC_003186 | | / |
| 21 | LNC_003187 | | 3.29828 |
| 22 | LNC_003184 | | / |
| 23 | LNC_003185 | | / |
| 24 | LNC_003401 | | / |
| 25 | LNC_003400 | | / |
| 26 | LNC_003188 | Chr14: 31.4-31.5 | / |
| 27 | LNC_003406 | | / |
| 28 | LNC_003405 | | / |
| 29 | LNC_003404 | | / |
| 30 | LNC_003403 | | / |
| 31 | LNC_003402 | | / |

**Table S38 miRNAs in the hermaphrodite production-associated regions**

| Serial number | Gene ID | Genomic Region (Mb) | Comparison of expression levels (AM_M vs AM_H) |
|---|---|---|---|
| 1 | zma-miR167j-3p | Chr2: 28.9-29.2 | / |
| 2 | novel_20 | Chr2: 29.62-29.83 | / |
| 3 | novel_216 | | / |
| 4 | aly-miR395e-5p | Chr2: 30.78-30.9 | / |
| 5 | ata-miR164b-3p | Chr14: 8.65-8.95 | / |
| 6 | novel_143 | | 1.2807 |
| 7 | cre-miR1171 | Chr14: 9.80-9.95 | / |
| 8 | atr-miR390.1 | Chr14: 30.9-31.4 | -2.8853 |
| 9 | pab-miR3711 | | / |

**Table S39 circRNAs in the hermaphrodite production-associated regions**

| Serial number | Gene ID | Genomic Region (Mb) | Comparison of expression levels (AM_M vs AM_H) |
|---|---|---|---|
| 1 | Dol_circ_0006799 | Chr2: 28.6-28.75 | / |
| 2 | Dol_circ_0006800 | | / |
| 3 | Dol_circ_0006801 | | / |
| 4 | Dol_circ_0006802 | | / |
| 5 | Dol_circ_0006803 | | / |

| | | | |
|---|---|---|---|
| 6 | Dol_circ_0006804 | | / |
| 7 | Dol_circ_0006805 | | / |
| 8 | Dol_circ_0006808 | Chr2: 28.9-29.2 | / |
| 9 | Dol_circ_0006809 | | / |
| 10 | Dol_circ_0006810 | | / |
| 11 | Dol_circ_0006811 | | / |
| 12 | Dol_circ_0006812 | | / |
| 13 | Dol_circ_0006813 | | / |
| 14 | Dol_circ_0006824 | Chr2: 29.62-29.83 | / |
| 15 | Dol_circ_0006825 | | / |
| 16 | Dol_circ_0006826 | | / |
| 17 | Dol_circ_0006827 | | / |
| 18 | Dol_circ_0006828 | | / |
| 19 | Dol_circ_0006829 | | / |
| 20 | Dol_circ_0006830 | | / |
| 21 | Dol_circ_0006831 | | / |
| 22 | Dol_circ_0006832 | | / |
| 23 | Dol_circ_0000693 | Chr11: 21.0-21.12 | / |
| 24 | Dol_circ_0000694 | | / |
| 25 | Dol_circ_0004511 | Chr14: 9.80-9.95 | / |
| 26 | Dol_circ_0004512 | | / |
| 27 | Dol_circ_0004513 | | / |
| 28 | Dol_circ_0004514 | | / |
| 29 | Dol_circ_0004515 | | / |
| 30 | Dol_circ_0004516 | | / |
| 31 | Dol_circ_0004517 | | / |
| 32 | Dol_circ_0004518 | | / |
| 33 | Dol_circ_0003833 | Chr14: 30.9-31.4 | / |
| 34 | Dol_circ_0003835 | | / |
| 35 | Dol_circ_0003836 | | / |
| 36 | Dol_circ_0003837 | | / |
| 37 | Dol_circ_0003838 | | / |
| 38 | Dol_circ_0003839 | | / |
| 39 | Dol_circ_0003841 | Chr14: 31.4-31.5 | / |

108

**Supplementary Table S40 Location and Category of SNPs and indels before LD pruning of 150 samples**

| Category | | Number of SNPs |
|---|---|---|
| Upstream (located in 1 Kb upstream of a gene) | | 833,595 |
| Exonic | Stop gain | 3,938 |
| | Stop loss | 573 |
| | Synonymous | 164,556 |
| | Non-synonymous | 182,984 |
| Intronic | | 5,238,872 |
| Splicing (located in a 2 bp region in intron close to the boundary of extron and intron ) | | 2,209 |
| Downstream (located in 1 Kb downstream of a gene) | | 683,392 |
| Upstream/downstream (located in 1 Kb upstream of a gene and in 1Kb downstream of another gene) | | 79,444 |
| Intergenic | | 16,456,349 |
| ts (transitions) | | 17,749,062 |
| tv (transversions) | | 6,178,944 |
| ts/tv (ratio of transitions to transversions) | | 2.876 |
| Total SNPs and Indels | | 23,928,006 |

Note: Total SNPs and Indels after LD pruning was 3,545,359 and 318,863, respectively, which were used for GWAS.

**Table S41 Summary of 12 individuals with inconsistency between sex expression and genotype in the top 10 SNPs associated with male expression**

| Individual ID | Description/putative phenotype | Position on chromosome 4 | | | | | | | | | | Proportion of female branches | *OGI* |
|---|---|---|---|---|---|---|---|---|---|---|---|---|---|
| | | 24,820,665 | 24,862,019 | 25,007,882 | 25,154,020 | 25,207,091 | 25,483,064 | 25,602,661 | 25,826,008 | 28,581,683 | 29,088,469 | | |
| 38 | female-biased monoecious plant | CT | CT | GT | GA | GA | GT | AG | GT | GA | GT | 1 | + |
| 62 | female-biased monoecious plant | CT | CT | GT | * | GA | GT | AG | GT | GA | GT | 1 | + |
| 80 | female-biased monoecious plant | CT | CT | GT | GA | GA | GT | AG | GT | GA | GT | 1 | + |
| 175 | female-biased monoecious plant | * | CT | GT | GA | GA | GT | AG | GT | GA | GT | 1 | + |
| 59 | male with haplotype recombined at OGI-flanking region | CT | CT | GT | GA | GA | GT | AA | GT | GA | GT | 0 | + |
| 157 | male with haplotype recombined at OGI-flanking region | * | CT | GT | GA | AA | GT | AG | GT | GA | GT | 0 | + |
| 161 | male with haplotype recombined at OGI-flanking region | CT | CT | GT | GA | GA | GT | * | GT | AA | TT | 0 | + |
| 188 | male with haplotype recombined at | CT | TT | GT | GA | GA | GT | * | GG | GA | GT | 0 | + |

| | | | | | | | | | | | | | |
|---|---|---|---|---|---|---|---|---|---|---|---|---|---|
| 204 | OGI-flanking region male with haplotype recombined at OGI-flanking region | TT | TT | GT | GA | GA | GT | * | GG | GA | GT | 0 | + |
| 5 | pseudo-monoecious plant | CC | CC | GG | * | AA | GG | AA | TT | GG | * | 0.8 | — |
| 6 | pseudo-monoecious plant | CC | CC | GG | * | AA | GG | AA | TT | GG | GG | 0.8 | — |
| 179 | monoecious plant | CC | CC | GG | GG | AA | GG | AA | TT | GG | GG | 0.7 | — |
| male | | CT | CT | GT | GA | GA | GT | AG | GT | GA | GT | | + |
| female | | CC | CC | GG | GG | AA | GG | AA | TT | GG | GG | | — |

Note: Yellow highlight represents Y allele-homozygous genotype; asterisk represents missing genotype.

117 **Table S42 Genes in the genomic region of Chr4: 22.0-32.0 Mb**

| Serial number | Gene ID | Genomic Region (Mb) | Gene name with Swissprot Annotation | Comparison of expression levels (M_F vs M_M) | Comparison of expression levels (G_F vs A_M) | Comparison of expression levels (AM_M vs AM_H) |
|---|---|---|---|---|---|---|
| 1 | evm.model.Chr4.1448 | Chr4: 22.0-24.4 | *SECA2* | / | / | / |
| 2 | evm.model.Chr4.1449 | | *At1g77330* | / | / | / |
| 3 | evm.model.Chr4.1450 | | *Os11g0191400* | / | / | / |
| 4 | evm.model.Chr4.1451 | | *NADK2* | / | -1.16668 | / |
| 5 | evm.model.Chr4.1452 | | *FBP24* | / | / | / |
| 6 | evm.model.Chr4.1453 | | *Pqbp1* | / | / | / |
| 7 | evm.model.Chr4.1454 | | NA | / | / | / |
| 8 | evm.model.Chr4.1455 | | NA | / | / | / |
| 9 | evm.model.Chr4.1456 | | *GLO* | -0.879045 | -2.37178 | 1.39711 |
| 10 | evm.model.Chr4.1458 | | *PERK9* | 1.00746 | / | / |
| 11 | evm.model.Chr4.1459 | | *USP* | / | / | / |
| 12 | evm.model.Chr4.1459.1 | | NA | / | / | / |
| 13 | evm.model.Chr4.1460 | | *At1g60770* | / | / | / |
| 14 | evm.model.Chr4.1461 | | *CML44* | / | / | / |
| 15 | evm.model.Chr4.1462 | | NA | / | / | / |
| 16 | evm.model.Chr4.1463 | | NA | / | 2.16666 | / |
| 17 | evm.model.Chr4.1470 | | *CYP71A6* | / | / | / |
| 18 | evm.model.Chr4.1473 | | NA | / | / | / |
| 19 | evm.model.Chr4.1475 | | *Kat6a* | / | / | / |
| 20 | evm.model.Chr4.1476 | | *tom1l2* | / | / | / |
| 21 | evm.model.Chr4.1478 | | NA | / | / | / |
| 22 | evm.model.Chr4.1479 | | NA | / | 2.61391 | / |
| 23 | evm.model.Chr4.1480 | | *At1g21400* | / | / | / |
| 24 | evm.model.Chr4.1481 | | NA | / | / | / |
| 25 | evm.model.Chr4.1481.3 | | NA | / | / | / |
| 26 | evm.model.Chr4.1487 | | *PERK2* | / | / | / |
| 27 | evm.model.Chr4.1489 | | NA | / | / | / |
| 28 | evm.model.Chr4.1490 | | NA | / | -1.11421 | / |
| 29 | evm.model.Chr4.1491 | | *RABC1* | / | / | / |
| 30 | evm.model.Chr4.1492 | | *FOLB2* | / | / | / |
| 31 | evm.model.Chr4.1493 | | NA | / | / | / |
| 32 | evm.model.Chr4.1496 | | *MYB44* | / | / | -2.95032 |
| 33 | evm.model.Chr4.1498 | Chr4: 24.4-26.0 | *SCL1* | / | / | -0.927144 |
| 34 | evm.model.Chr4.1499 | | *SDN5* | / | / | / |
| 35 | evm.model.Chr4.1505 | | NA | / | / | / |
| 36 | evm.model.Chr4.1506 | | NA | / | / | / |
| 37 | evm.model.Chr4.1507 | | *NAC002* | / | / | / |
| 38 | evm.model.Chr4.1508 | | *PUB4* | / | / | 1.50908 |
| 39 | evm.model.Chr4.1509 | | *ASP1* | / | -1.35933 | / |
| 40 | evm.model.Chr4.150 | | NA | / | / | / |
| 41 | evm.model.Chr4.1518 | | NA | / | / | / |
| 42 | evm.model.Chr4.1519 | | *PDI* | / | / | / |
| 43 | evm.model.Chr4.151 | | NA | / | / | / |
| 44 | evm.model.Chr4.152 | | NA | / | / | / |
| 45 | evm.model.Chr4.1521 | | *LACS9* | / | / | / |
| 46 | evm.model.Chr4.1523 | | NA | 2.11684 | / | / |
| 47 | evm.model.Chr4.1525 | | NA | / | / | / |
| 48 | evm.model.Chr4.1527 | | *CAO* | / | / | / |
| 49 | evm.model.Chr4.1528 | | NA | / | / | / |
| 50 | evm.model.Chr4.1529 | | *ILL6* | / | -3.26906 | / |

| | | | | | | |
|---|---|---|---|---|---|---|
| 51 | evm.model.Chr4.152 | | NA | / | / | / |
| 52 | evm.model.Chr4.1534 | Chr4: | *ARR9* | 2.17868 | 3.7477 | -3.26959 |
| 53 | evm.model.Chr4.1536 | 26.0-27.4 | *PDS5A* | / | / | / |
| | evm.model.Chr4.1537 | | *PDS5A* | / | / | / |
| 55 | evm.model.Chr4.1538 | | *UDP-GALT1* | / | 0.961092 | 0.919472 |
| 56 | evm.model.Chr4.153 | | NA | / | / | / |
| 57 | evm.model.Chr4.1540 | | *NFYA3* | / | / | / |
| 58 | evm.model.Chr4.1541 | | NA | / | / | / |
| 59 | evm.model.Chr4.1542 | | *TRS120* | / | / | / |
| 60 | evm.model.Chr4.1543 | | *PCMP-H32* | / | / | / |
| 61 | evm.model.Chr4.1546 | | *GALE* | / | / | / |
| 62 | evm.model.Chr4.1547 | | *UGE1* | / | / | / |
| 63 | evm.model.Chr4.1548.1 | | NA | / | / | / |
| 64 | evm.model.Chr4.1549 | | *UGE1* | / | 1.41701 | -1.0526 |
| 65 | evm.model.Chr4.1551 | | NA | / | / | / |
| 66 | evm.model.Chr4.1553 | | *TRS120* | / | / | / |
| 67 | evm.model.Chr4.1554 | 27.4-28.0 | NA | / | / | / |
| 68 | evm.model.Chr4.1558 | | NA | / | 1.75605 | / |
| 69 | evm.model.Chr4.1560 | 28.0-29.2 | *SULTR4;1* | / | / | / |
| 70 | evm.model.Chr4.1560.3 | | NA | / | / | / |
| 71 | evm.model.Chr4.1562 | | NA | / | / | / |
| 72 | evm.model.Chr4.1563 | | *DIR9* | / | / | / |
| 73 | evm.model.Chr4.1564 | | NA | / | / | / |
| 74 | evm.model.Chr4.1566 | | NA | / | 4.17624 | / |
| 75 | evm.model.Chr4.1567 | | NA | / | / | / |
| 76 | evm.model.Chr4.1568 | | NA | / | 4.86363 | / |
| 77 | evm.model.Chr4.1569 | | NA | / | / | / |
| 78 | evm.model.Chr4.1570 | | *MTP5* | / | / | / |
| 79 | evm.model.Chr4.1574 | | NA | / | / | / |
| 80 | evm.model.Chr4.1575 | | NA | / | / | / |
| 81 | evm.model.Chr4.1577 | | *POLD1* | / | / | / |
| 82 | evm.model.Chr4.1579 | | NA | / | / | / |
| 83 | evm.model.Chr4.1580 | | NA | / | / | / |
| 84 | evm.model.Chr4.1583 | 29.2-32.0 | *csd* | / | / | / |
| 85 | evm.model.Chr4.1584 | | *At1g04910* | / | / | / |
| 86 | evm.model.Chr4.1585 | | NA | / | -inf | 2.97094 |
| 87 | evm.model.Chr4.1586 | | NA | / | / | / |
| 88 | evm.model.Chr4.1587 | | NA | / | / | / |
| 89 | evm.model.Chr4.158 | | NA | / | / | / |
| 90 | evm.model.Chr4.1590 | | NA | / | / | / |
| 91 | evm.model.Chr4.1591 | | *NAC078* | / | / | / |
| 92 | evm.model.Chr4.1592 | | NA | / | / | / |
| 93 | evm.model.Chr4.1593 | | PRL1-IFG | / | / | / |
| 94 | evm.model.Chr4.1596 | | *CALS5* | -4.87137 | -6.32596 | 4.44529 |
| 95 | evm.model.Chr4.1597 | | *CALS5* | -5.19545 | -6.5116 | 4.41433 |
| 96 | evm.model.Chr4.1602 | | *YDA* | / | / | / |
| 97 | evm.model.Chr4.1603 | | *ndhO* | / | / | / |
| 98 | evm.model.Chr4.1604 | | *At5g09310* | / | / | / |
| 99 | evm.model.Chr4.1604.1 | | NA | / | / | / |
| 100 | evm.model.Chr4.1605 | | *PSAN* | / | / | / |
| 101 | evm.model.Chr4.1606 | | *GSVIVT00026 920001* | / | -2.976 | / |
| 102 | evm.model.Chr4.1607 | | NA | / | / | / |
| 103 | evm.model.Chr4.1610 | | *PUB25* | / | / | -1.98649 |
| 104 | evm.model.Chr4.1611 | | NA | / | / | / |
| 105 | evm.model.Chr4.1612 | | *PUMP4* | / | / | / |
| 106 | evm.model.Chr4.1614 | | *CHX15* | / | -7.8179 | 1.46584 |

| | | | | | |
|---|---|---|---|---|---|
| 107 | evm.model.Chr4.1618 | *SAL1* | / | / | / |
| 108 | evm.model.Chr4.1618.4 | NA | / | / | / |
| 109 | evm.model.Chr4.1619 | *PCMP-E76* | / | / | / |
| 110 | evm.model.Chr4.161 | NA | / | / | / |
| 111 | evm.model.Chr4.1622 | *At5g64030* | / | / | / |
| 112 | evm.model.BG.131.p | NA | / | / | / |
| 113 | evm.model.Chr4.1626 | *FPP* | / | / | / |
| 114 | evm.model.Chr4.1627 | NA | / | / | / |
| 115 | evm.model.Chr4.1628 | *CBSCBSPB3* | / | / | / |
| 116 | evm.model.Chr4.1631 | *At4g08330* | / | / | / |
| 117 | evm.model.Chr4.1632 | NA | / | / | / |
| 118 | evm.model.Chr4.1641 | *nep1* | 2.35702 | 1.79766 | -3.20557 |
| 119 | evm.model.Chr4.1642 | *SAUR32* | / | / | / |
| 120 | evm.model.Chr4.1645 | *NFYB7* | -inf | / | 5.3057 |
| 121 | evm.model.Chr4.1647 | *CDKE-1* | / | / | / |
| 122 | evm.model.Chr4.164 | NA | / | / | / |
| 123 | evm.model.Chr4.1653 | NA | / | / | / |
| 124 | evm.model.Chr4.1654 | NA | / | / | / |
| 125 | evm.model.Chr4.1656 | *NAC002* | / | 1.29508 | -1.03713 |
| 126 | evm.model.Chr4.1657 | *UVR8* | / | / | / |
| 127 | evm.model.Chr4.1660 | *KIWI* | / | / | / |
| 128 | evm.model.Chr4.1661 | *SRT2* | / | / | / |

Note: "inf" represents infinite, and this was attributed to that the expression level of one sample in the comparative combination was zero.
